## Supplemental data for "Glycan-induced protein dynamics in human norovirus P dimers depend on virus strain and deamidation status"

### Contents

1) Sequence alignment and homology modeling ........ S2
2) Identification of deamidation ........ S4
3) Native MS ........ S12
4) Analysis of bimodal spectra ........ S14
5) HDX-MS coverage maps ........ S15
6) HDX summary tables ........ S20
7) MS-Viewer search keys ........ S25
8) MD simulations ........ S26
9) Deuterium uptake plots ........ S34

### 1) Sequence alignment and homology modeling

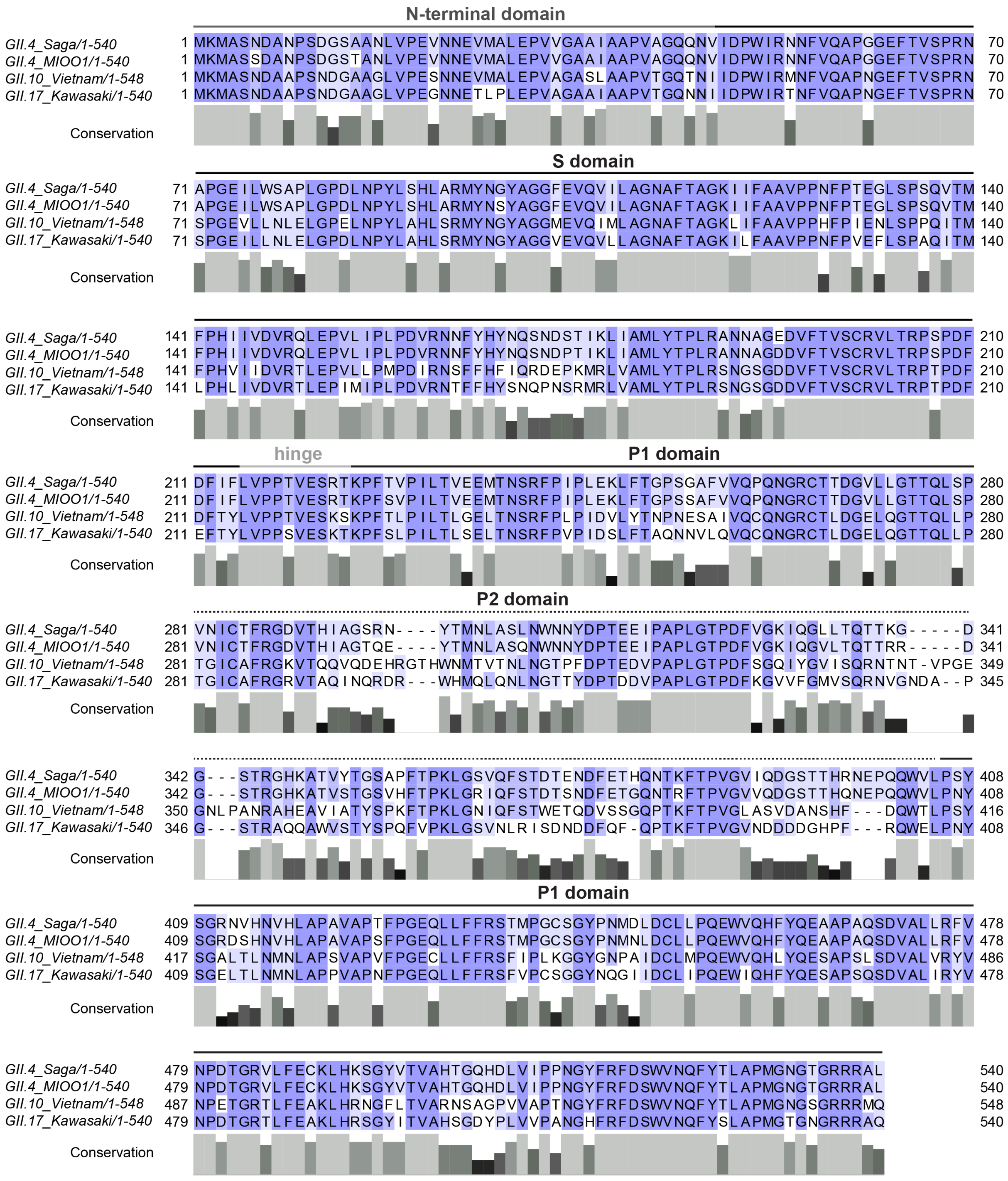

Figure S 1: Sequence alignment of GII.4 Saga, GII.4 MI001, GII.10 Vietnam and GII.17 Kawasaki major capsid protein (VP1) sequences. Color coding represents sequence identity (blue), similarity (light blue) or difference (white). Bars below the sequence indicate the degree of amino acid conservation among the strains at the respective position (low (black) to high (light grey)).

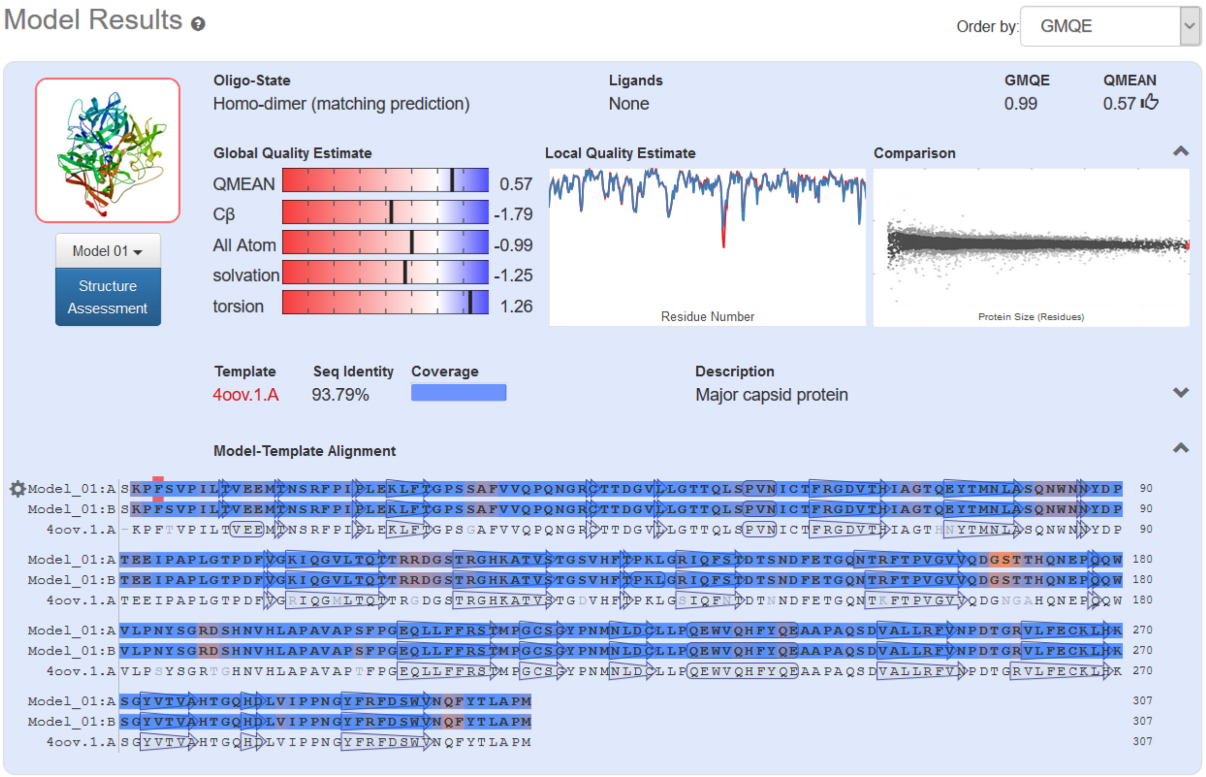


Figure S 2: SWISS MODEL result for the GII.4 MI001 P dimer homology model indicating good model accuracy.

### 2) Identification of deamidation

Table S 1: Identification of deamidation sites in GII.4 MI001 and GII.4 Saga P dimers stored in buffers of different pH. No deamidation sites were identified in GII.10 Vietnam and GII.17 Kawasaki P dimers. Deamidation positions and probability were calculated in MaxQuant based on peptide fragment spectra.

| Strain (storage pH) | Peptide sequence | Deamidated position | Deamidation position probability | deamidated fraction in % |
|---|---|---|---|---|
| **GII.4 MI001 (pH 4.9)** | RSTMPGCSGYPNM**N**L | N448 | 0.99 | 8 |
| **GII.4 MI001 pH (7.3)** | FRSTMPGCSGYPNM**N**L | N448 | 0.82 | 2 |
| | STDTS**N**DFETGQNTRF | N373 | 1 | approx. 64 |
| | **N**SRFPIPLEKL | N239 | 0.98 | 7 |
| **GII.4 Saga pH (7.3)** | STDTE**N**DFETHQ | N373 | 0.96 | approx. 88 |

**A**

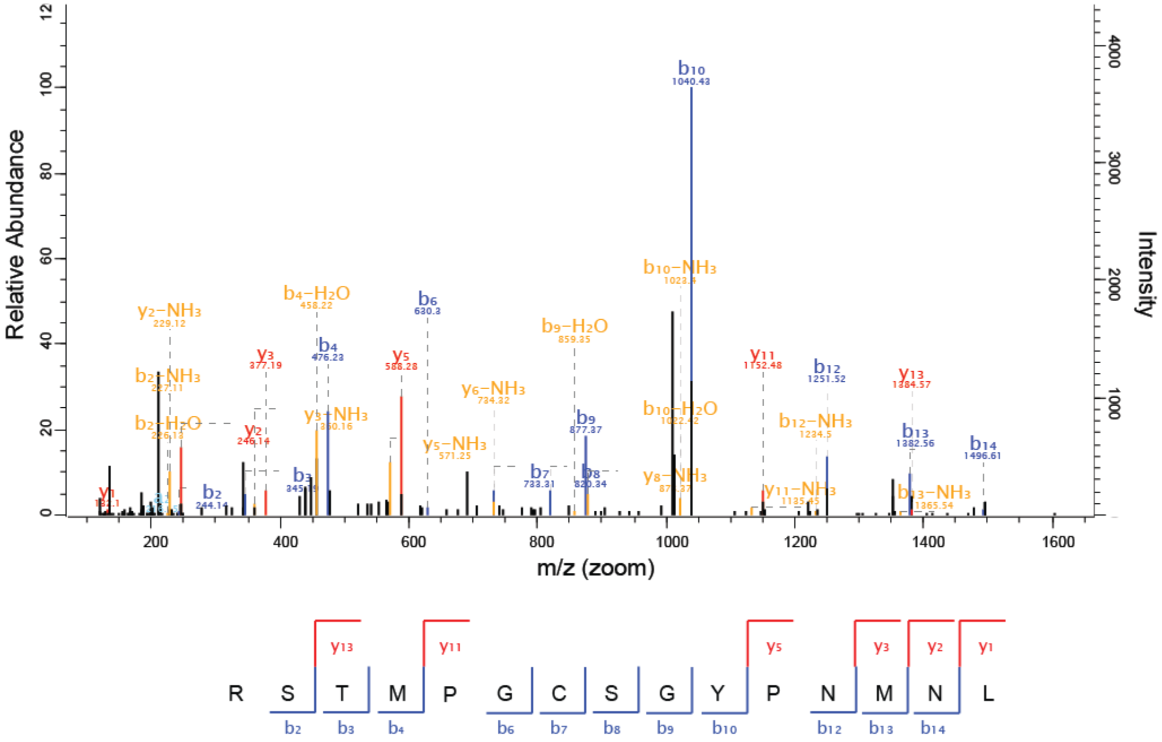


**B**

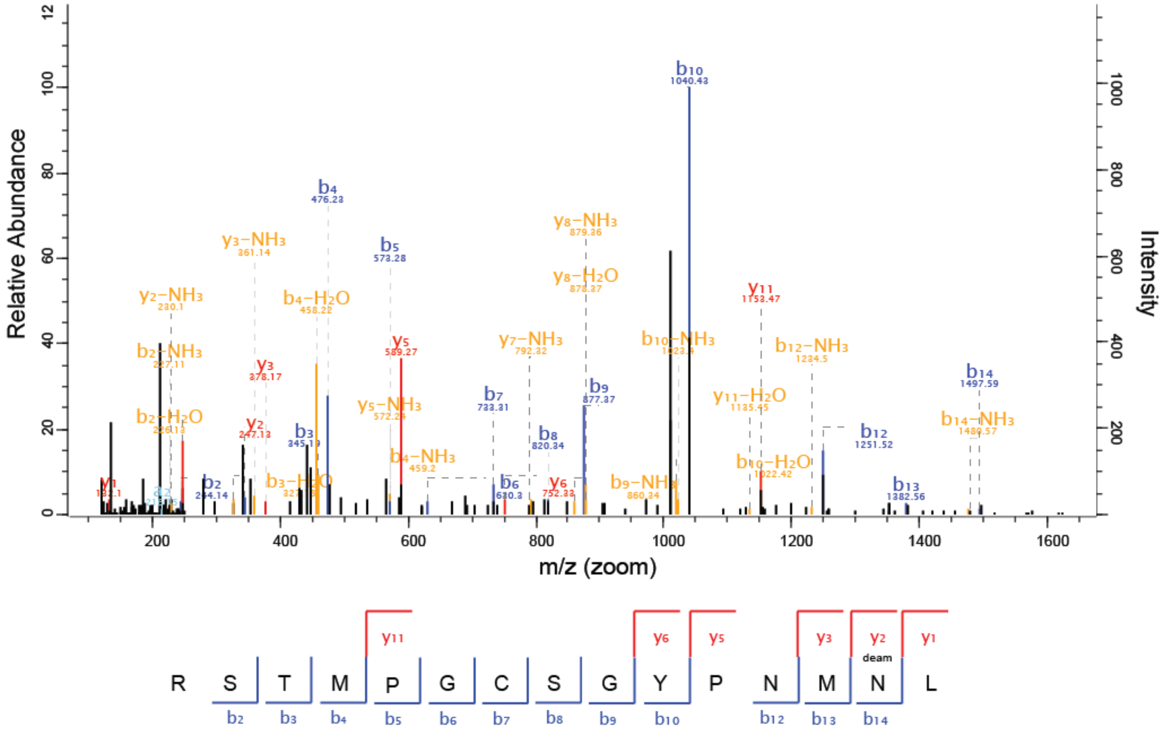


Figure S 3: Fragment spectra of wildtype (A) and deamidated (B) peptide RSTMPGCSGYPNMN$^{448}$L from GII.4 MI001 P dimer stored at pH 4.9 (5 months, 5°C).

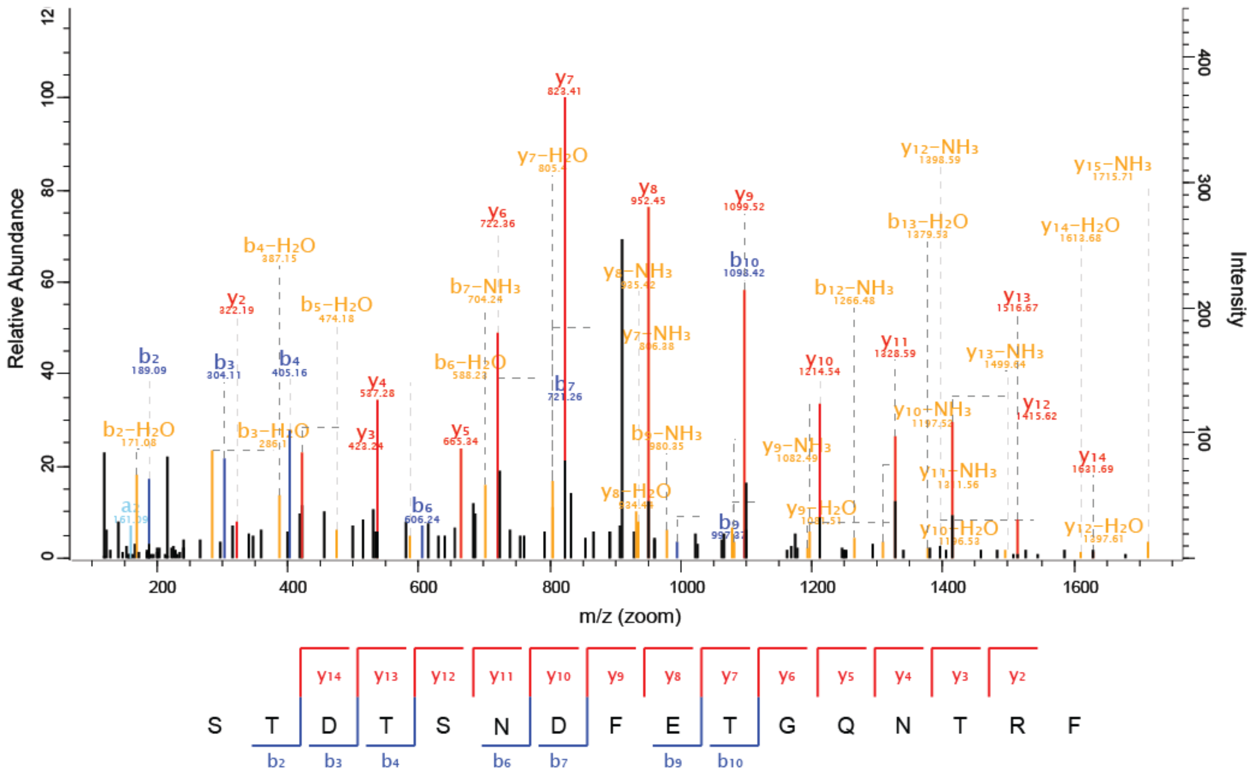

Figure S 4: Fragment spectra of wildtype peptide STDTSN[373]DFETGQNTRF from GII.4 MI001 P dimer stored at pH 4.9 (5 months, 5°C). No peptide carrying a deamidation at N373 could be detected.

**A**

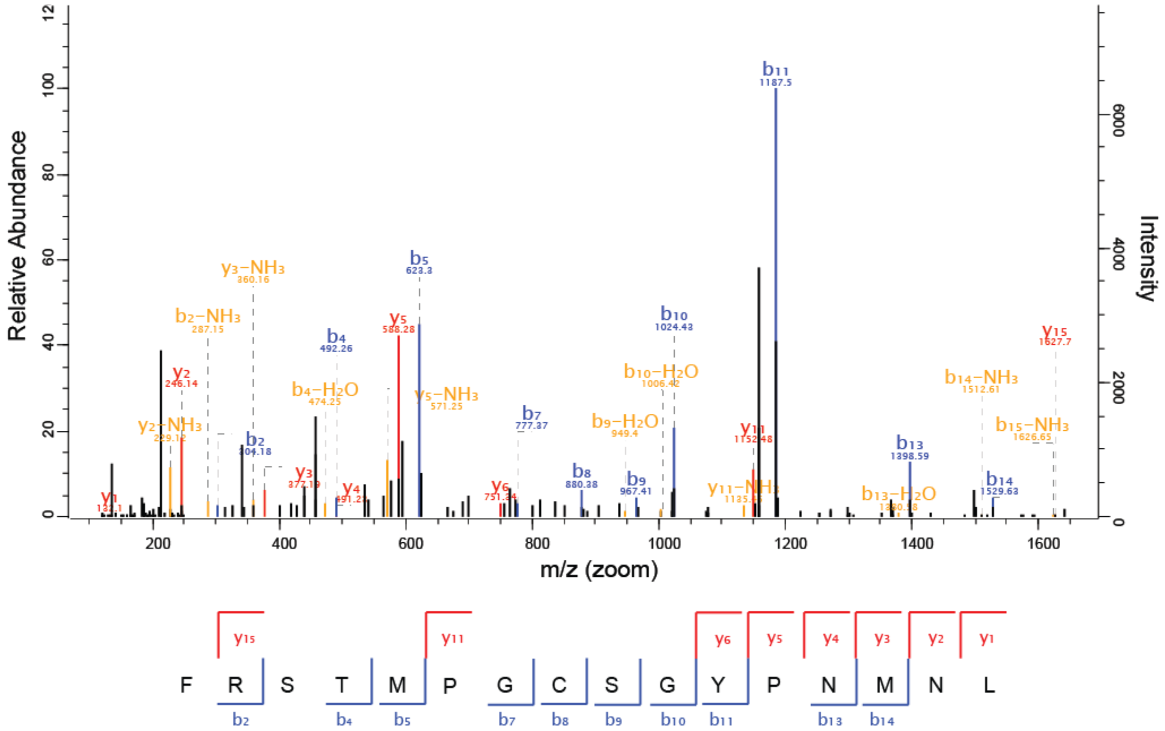


**B**

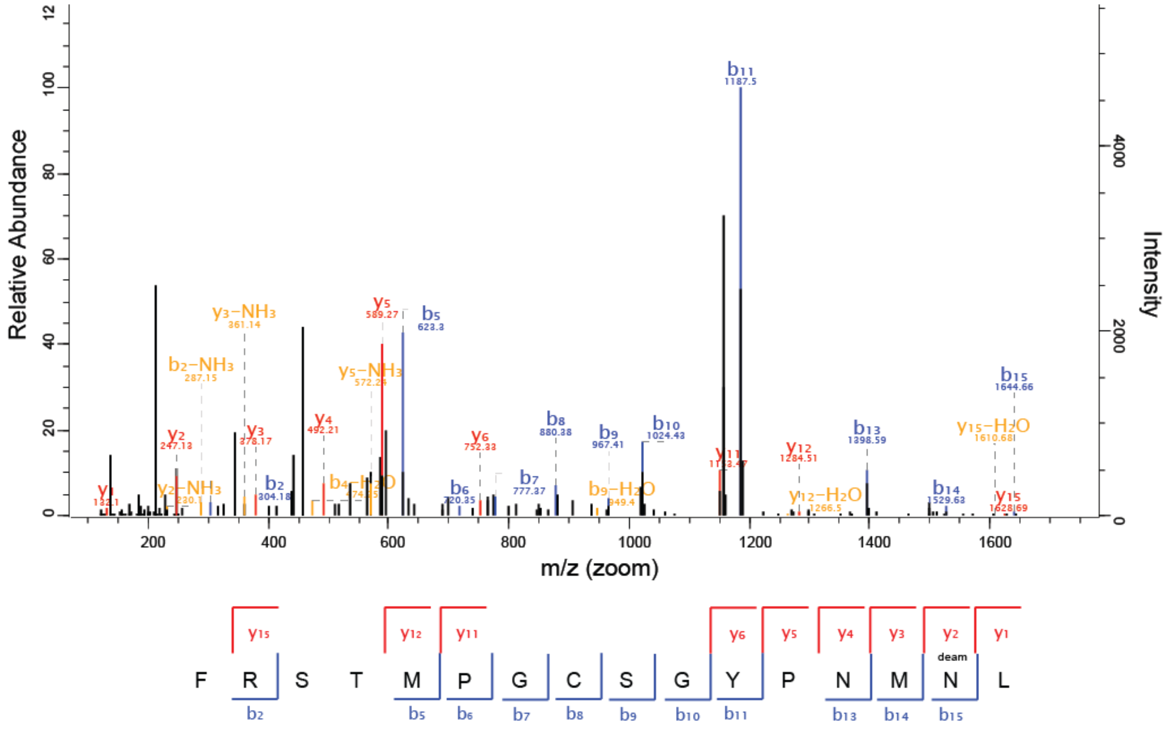


Figure S 5: Fragment spectra of wildtype (A) and deamidated (B) peptide FRSTMPGCSGYPNMN$^{448}$L from GII.4 MI001 P dimer stored at pH 7.3 (1 year, 5°C).

**A**

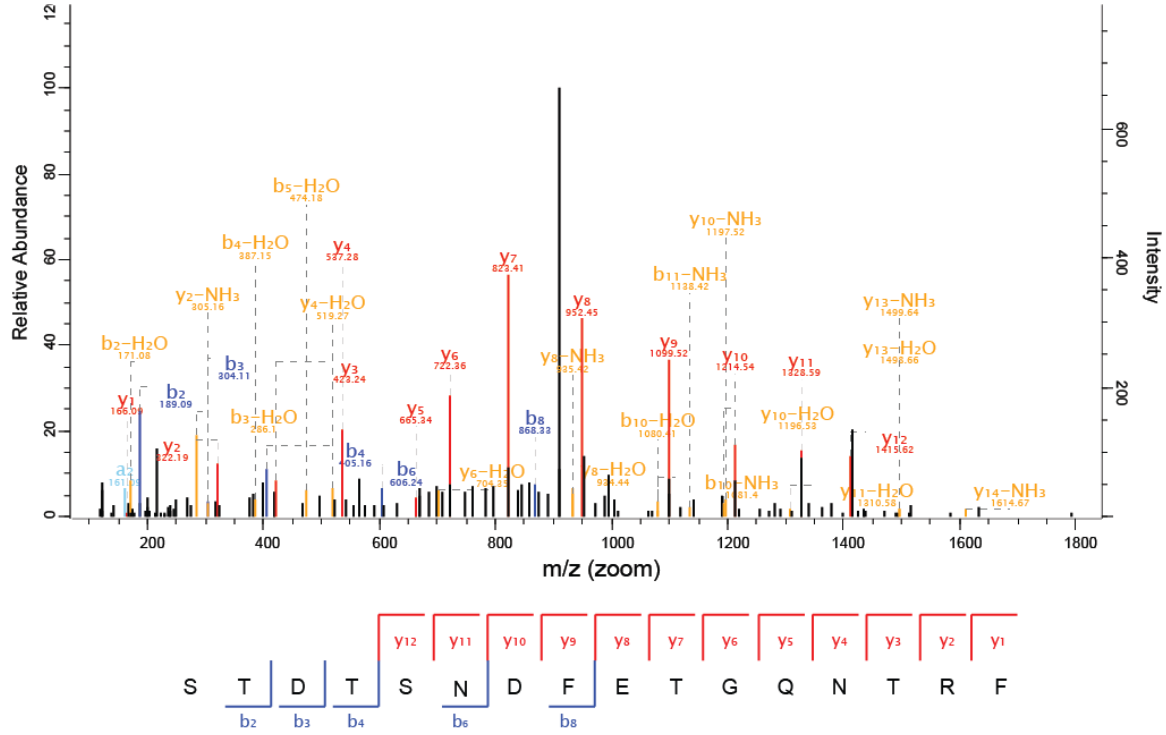


**B**

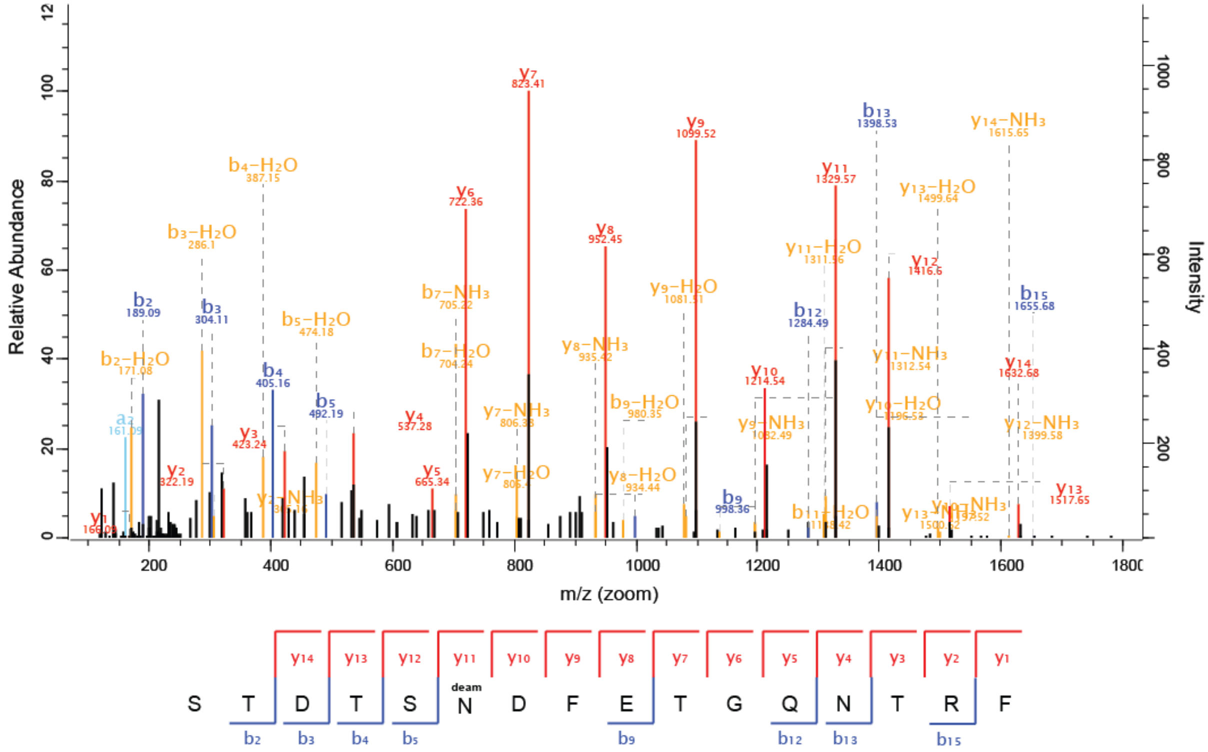


Figure S 6: Fragment spectra of wildtype (A) and deamidated (B) peptide STDTSN[373]DFETGQNTRF from GII.4 MI001 P dimer stored at pH 7.3 (1 year, 5°C).

**A**

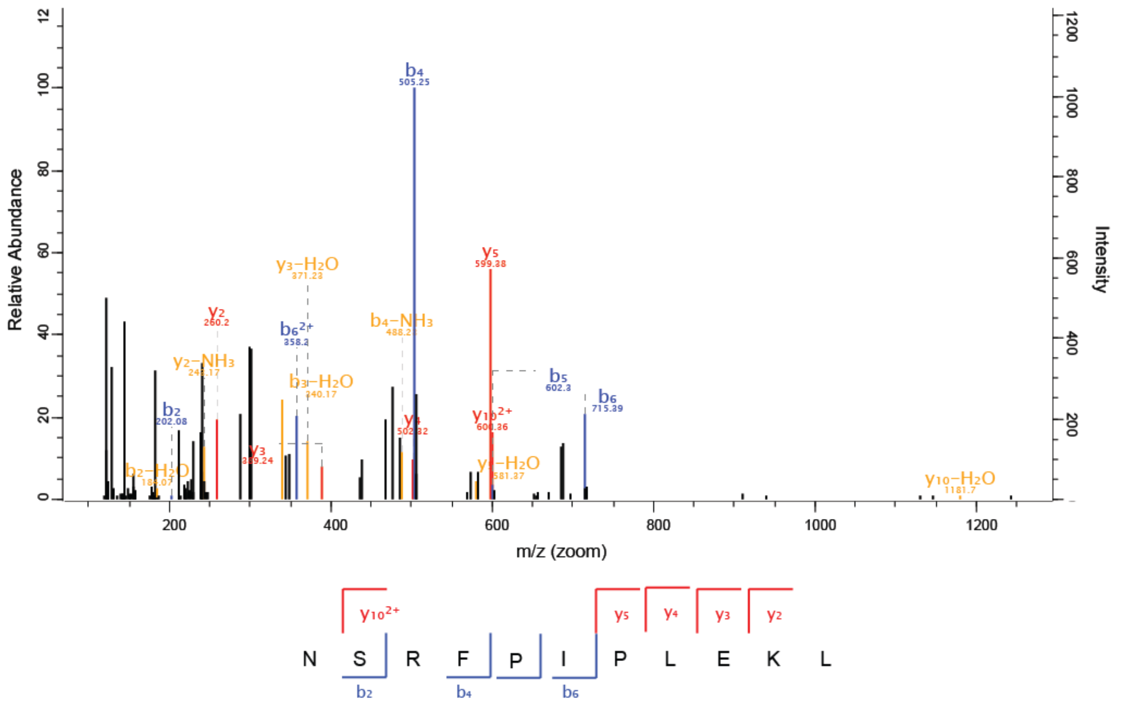


**B**

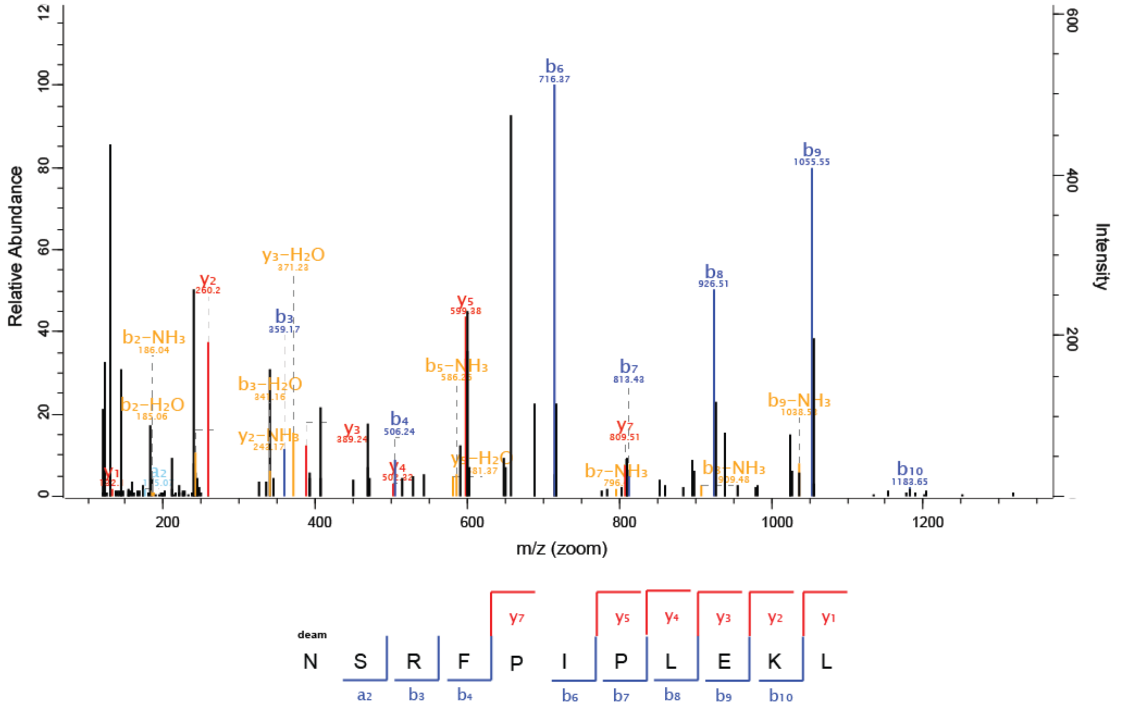


Figure S 7: Fragment spectra of wildtype (A) and deamidated (B) peptide N[239]SRFPIPLEKL from GII.4 MI001 P dimer stored at pH 7.3 (1 year, 5°C).

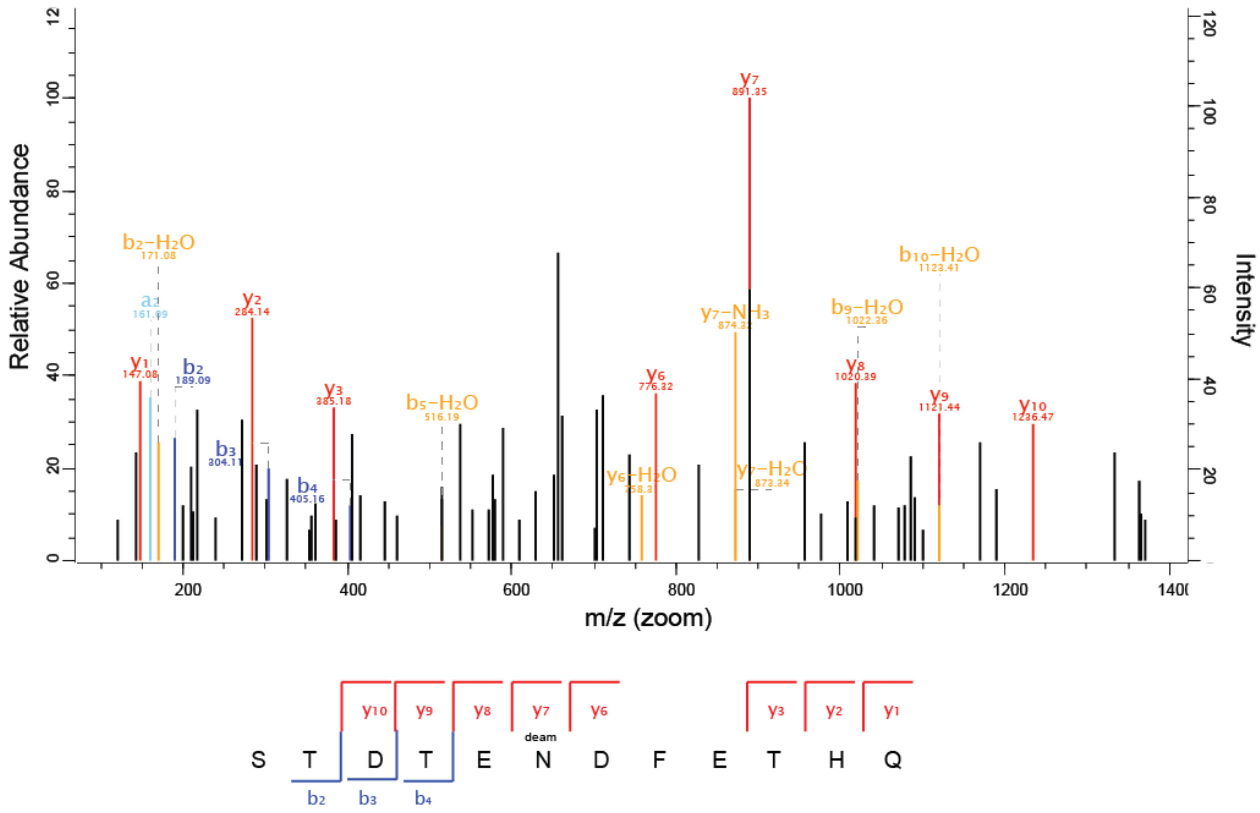


Figure S 8: Fragment spectra of deamidated peptide STDTEN$^{373}$DFETHQ from GII.4 Saga P dimer stored at pH 7.3 (more than 2 years, 5°C). Due to the low abundance wildtype N373 peptides did not get selected for fragmentation in the mass spectrometer and could therefore not be identified in the automated peptide search in MaxQuant. Therefore, MS data was searched manually for wildtype peptide precursors, which led to the identification of approximately 12 % N373 wildtype peptides.

**A**

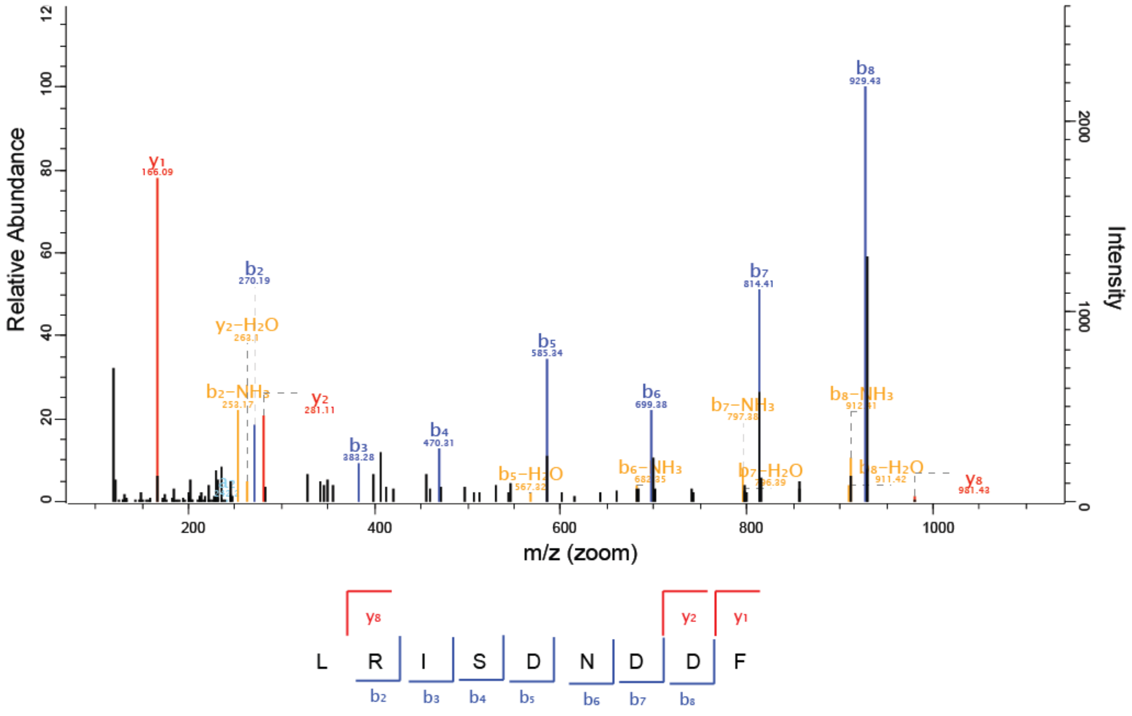


**B**

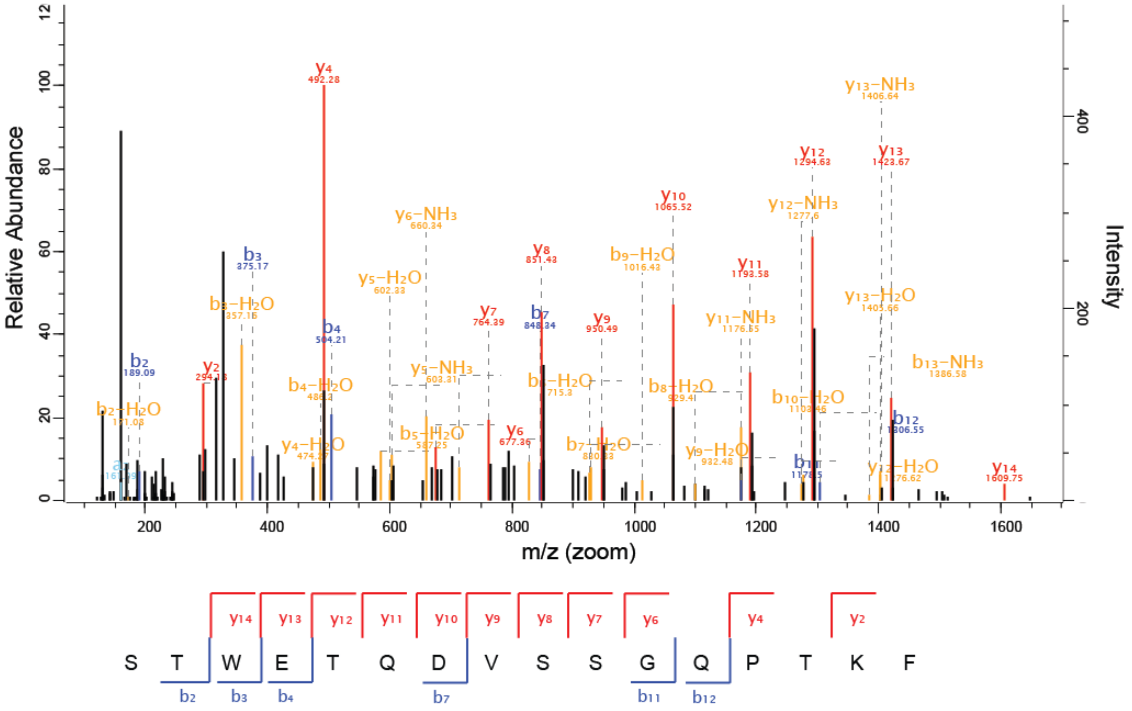


Figure S 9: Fragment spectra of peptides LRISDND[377]DF from GII.17 Kawasaki P dimer (A) and STWETQ[384]DVSSGQPTKF from GII.10 Vietnam P dimer (B). Both strains retain their wildtype sequence at the GII.4 N373 equivalent positon, even after more than a year of storage at 5°C.

### 3) Native MS

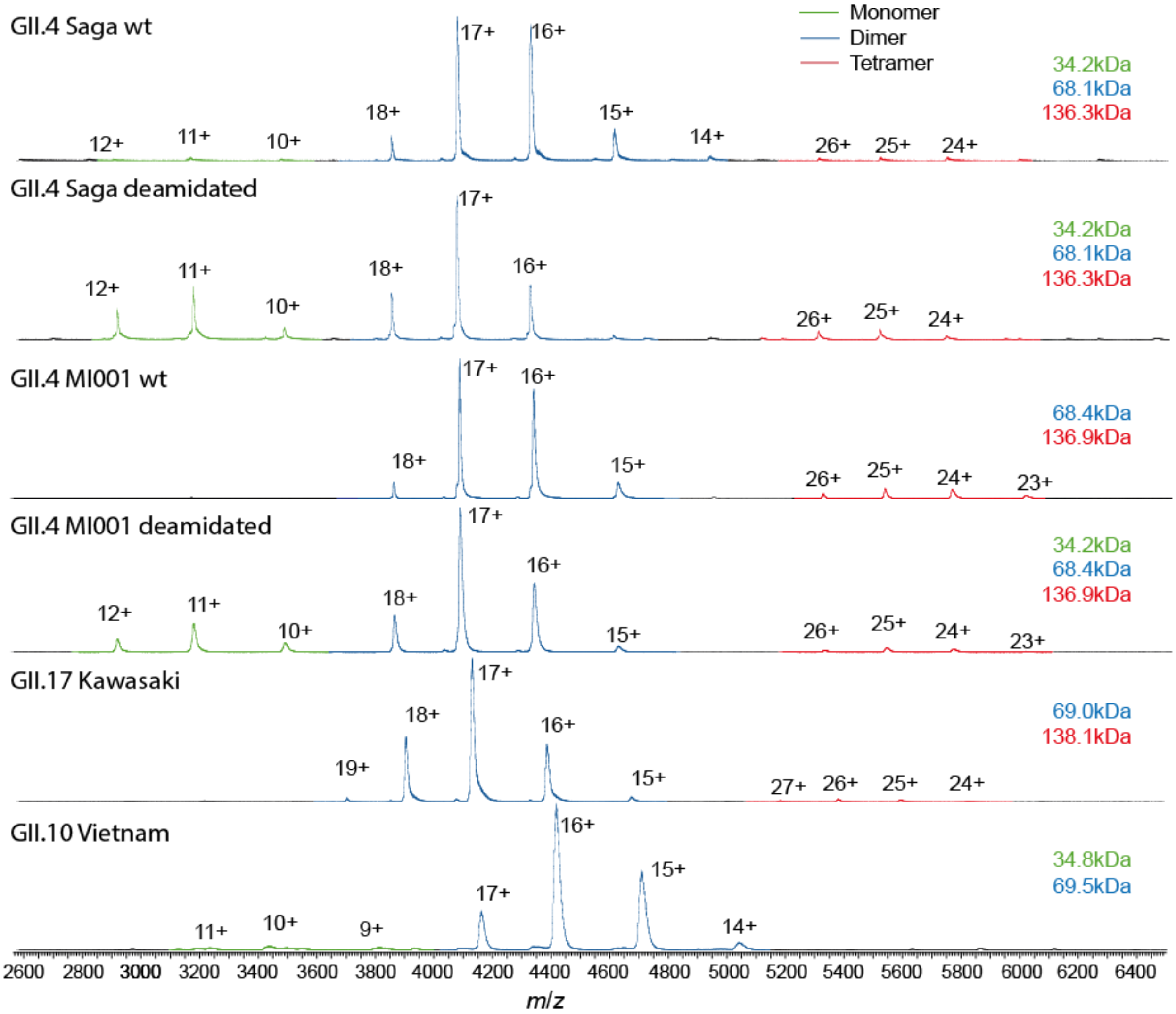


Figure S 10: Native mass spectra of different human norovirus P domain samples. GII.17 Kawasaki, GII.10 Vietnam, wildtype GII.4 MI001 and wildtype GII.4 Saga P domains are present as dimers with the expected molecular masses, apart from unspecific tetramers formed during the ESI process. Both deamidated GII.4 P domains (100 % deamidated GII.4 Saga and 64 % deamidated GII.4 MI001) are also present as monomers. No native MS data is available for the partially (88 %) deamidated GII.4 Saga P dimer. Quantification of the individual species in each strain can be found in Table S1.

Table S 2: Fractions of P monomers, dimers and tetramers from native MS (Figure S10)

| **Strain** | **Monomer fraction (%)** | **Dimer fraction (%)** | **Tetramer fraction (%)** |
|---|---|---|---|
| **GII.4 Saga deamidated (100%)** | **32** | **60** | 8 |
| **GII.4 Saga wildtype** | 0 | **100** | 0 |
| **GII.4 MI001 deamidated (64%)** | **16** | **80** | 4 |
| **GII.4 MI001 wildtype** | 0 | **91** | 9 |
| **GII.17 Kawasaki** | 0 | **100** | 0 |
| **GII.10 Vietnam** | 4 | **96** | 0 |

### 4) Analysis of bimodal spectra

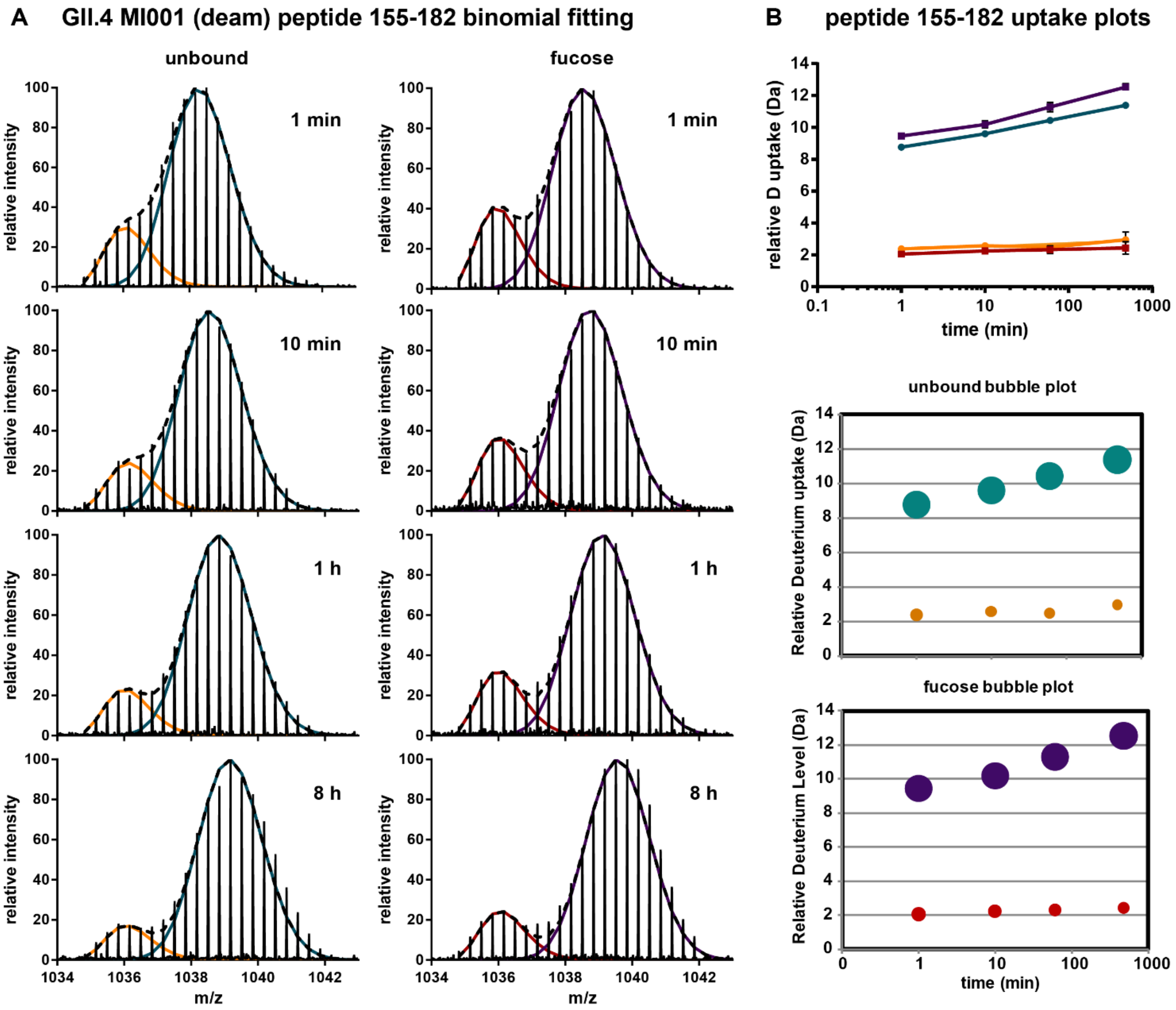


Figure S 11: Peptide HDX data analysis example for binomial fitting of bimodal spectra. A) Binomial fitting with bimodal deconvolution was applied in HXExpress resulting in a low (orange, red) and a high (green, violet) deuterated peak distribution for every state and time point. B) Deuteration values for the individual peak distributions can be visualized as uptake plot and significant differences between the unbound and ligand-bound state can be statistically analyzed. The relative deuterium uptake for each peak distribution can also be displayed as bubble plot, where the circle area corresponds to the relative intensity of the individual peak distribution at a given time point. Here, constant intensity ratios over time point towards the presence of two conformationally distinct protein populations.

### 5) HDX-MS coverage maps

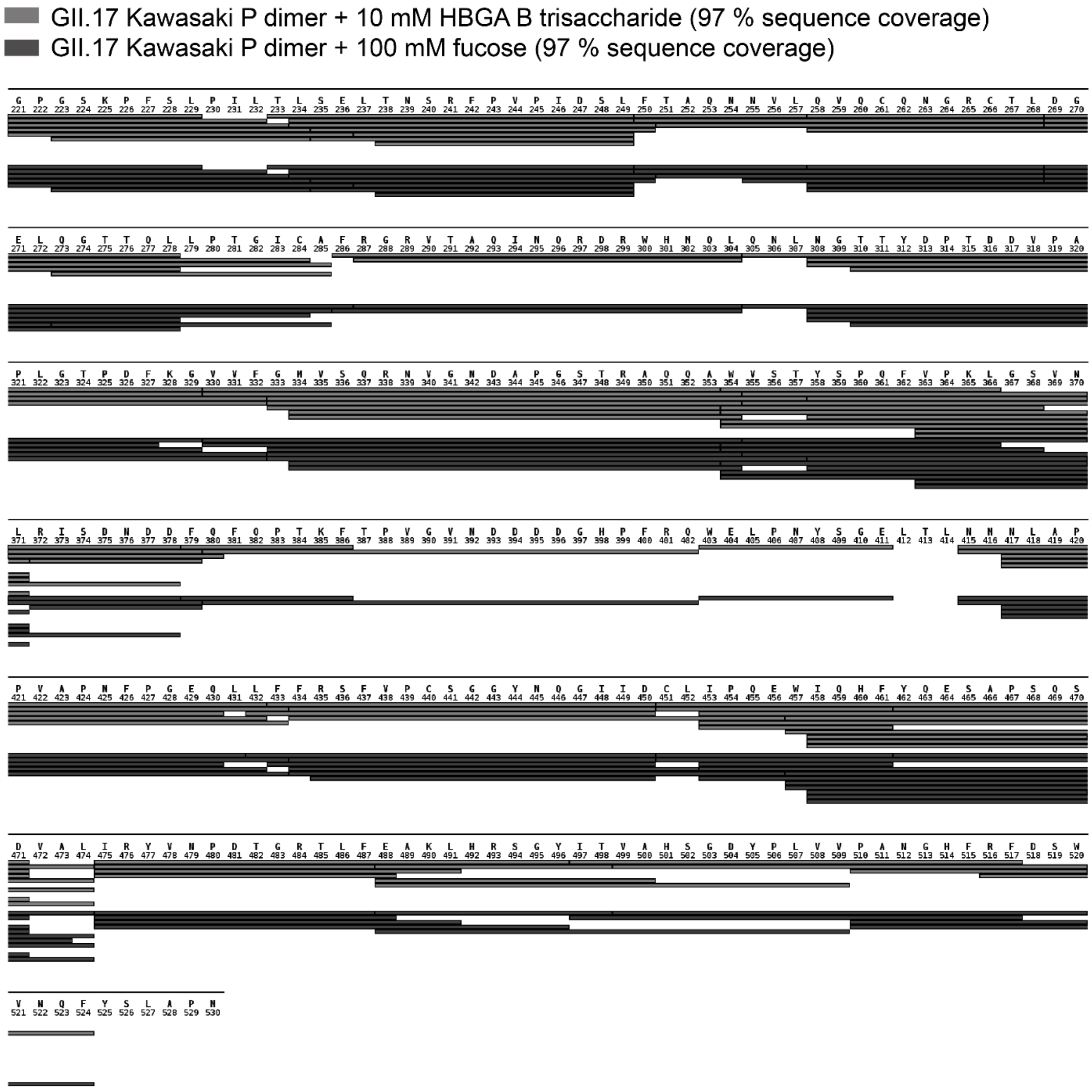

Figure S 12: GII.17 Kawasaki P domain effective peptide coverage map for HDX-MS experiments with HBGA B trisaccharide (grey) and fucose (dark grey).

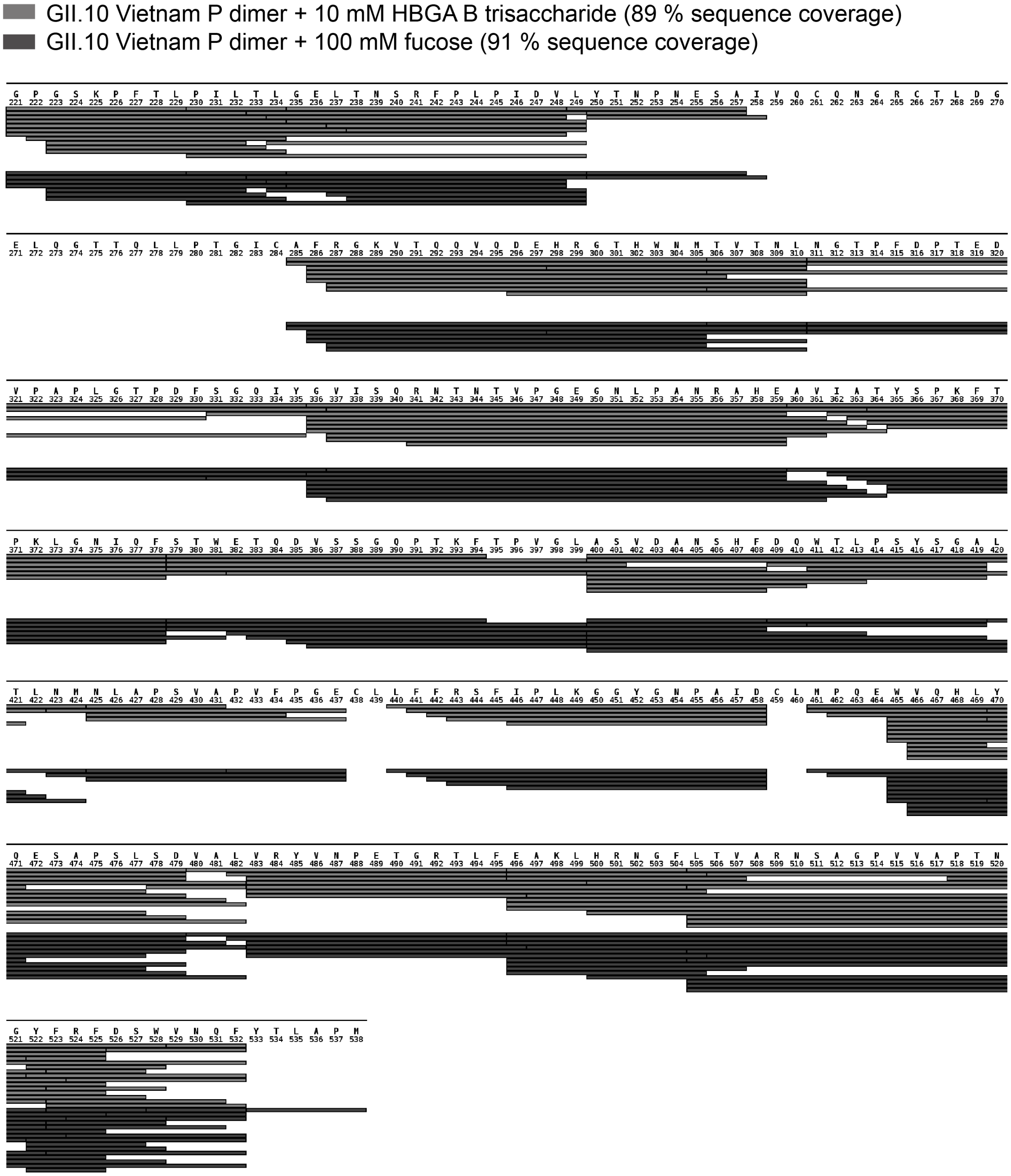


Figure S 13: GII.10 Vietnam P domain effective peptide coverage map for HDX-MS experiments with HBGA B trisaccharide (grey) and fucose (dark grey).

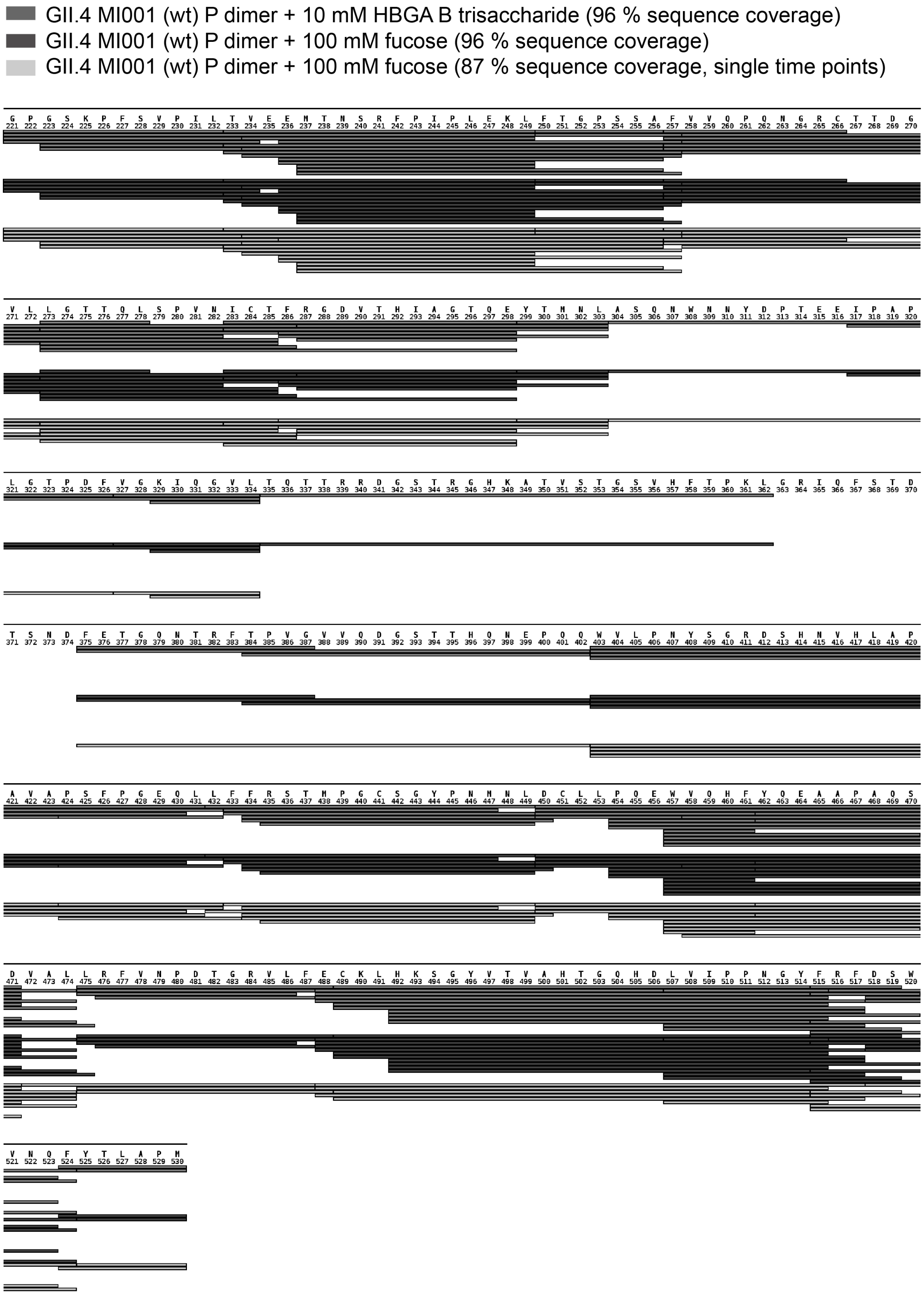


Figure S 14: GII.4 MI001 wildtype P domain effective peptide coverage map for HDX-MS experiments with HBGA B trisaccharide (grey) and fucose (triplicate measurement (dark grey) and single measurement (light grey)).

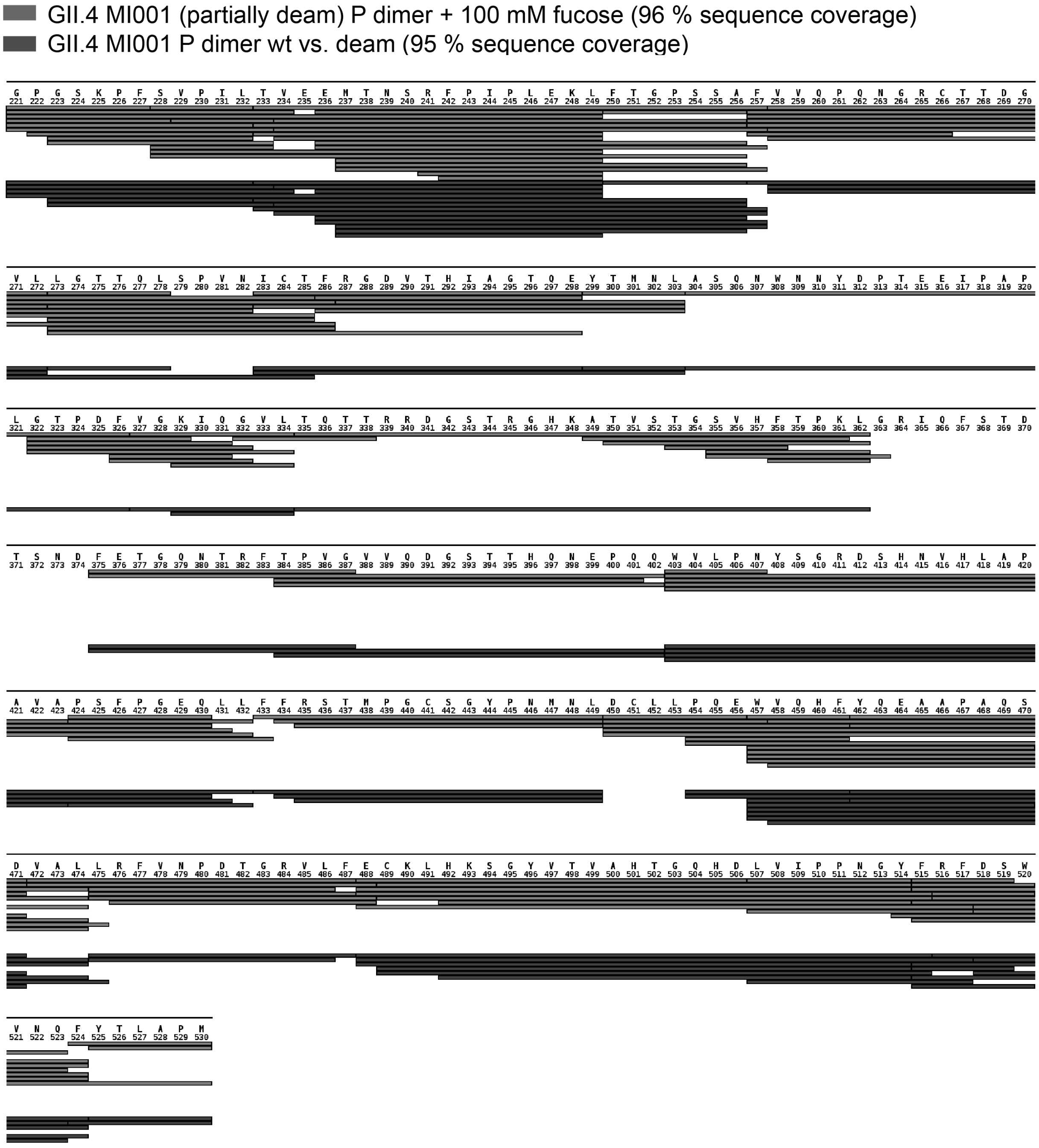


Figure S 15: GII.4 MI001 partially deamidated P domain effective peptide coverage map for HDX-MS experiments with fucose (grey) and wildtype vs. partially deamidated without ligand (dark grey).

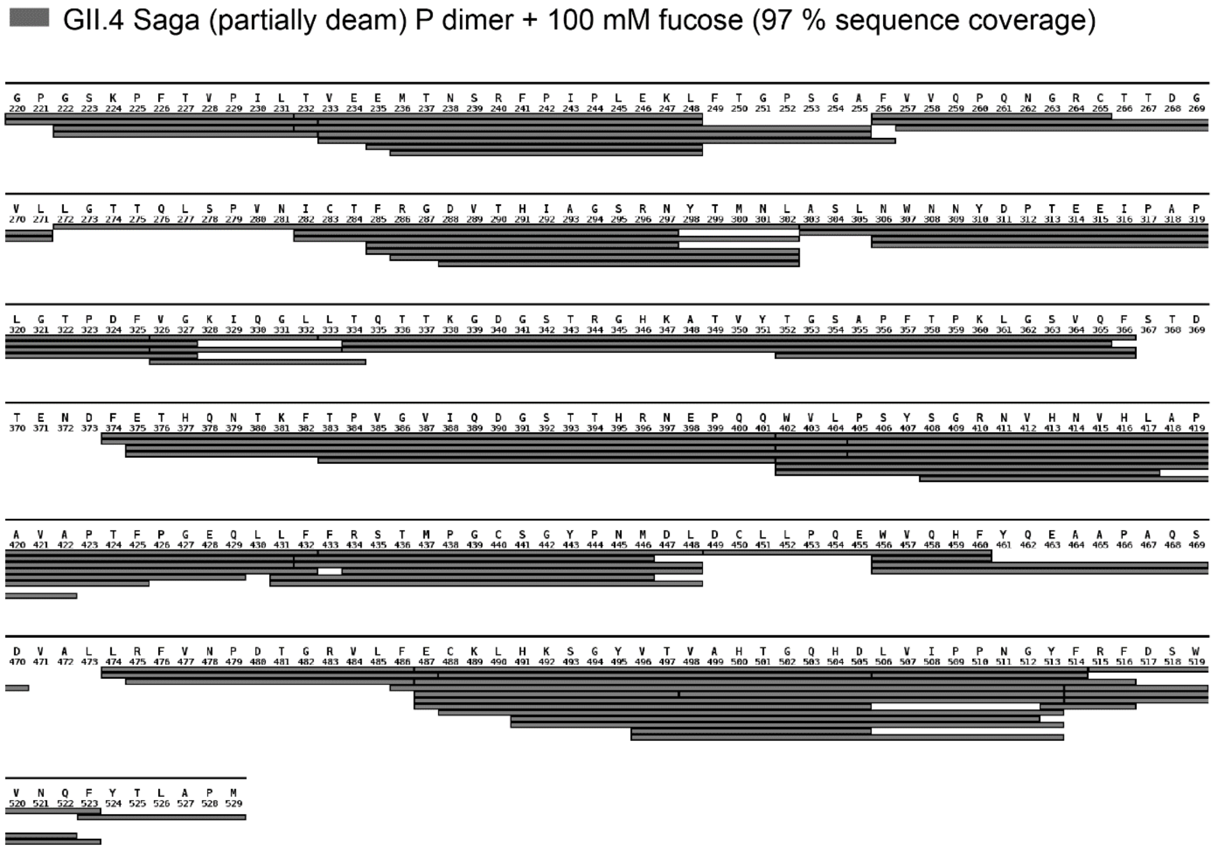


Figure S 16: GII.4 Saga partially deamidated P domain effective peptide coverage map for the HDX-MS experiment with fucose.

### 6) HDX summary tables

Table S 3: HDX summary tables for all datasets according to HDX-MS community recommendations (Masson, G.R., et al., Nat Methods, 2019. **16**(7): p. 595-602)

**T1** **GII.10 Vietnam P dimer**

| Data Set | unbound | 100 mM fucose |
|---|---|---|
| HDX reaction details | 20 mM Tris, 150 mM NaCl, pH 2.3, 25 °C | |
| HDX time course (min) | 1, 10, 60, 480 | |
| HDX control samples | FD model peptides in T9 | |
| Back-exchange (mean) | NA | |
| # of Peptides | 117 | 117 |
| Sequence coverage | 91% | 91% |
| Average peptide length / Redundancy | 15.9 / 5.8 | 15.9 / 5.8 |
| Replicates (biological or technical) | 3 (technical) | 3 (technical) |
| Repeatability (average standard deviation) | 0.1 | 0.073 |
| Significant differences in HDX (delta HDX > X D) | T test (p<0.05) and delta D > 0.25 (2x pooled average SD), delta D > 0.55 (99% percentile) for 8h time point single measurement, manual validation in case of bimodality | |

**T2** **GII.10 Vietnam P dimer**

| Data Set | unbound | 10 mM HBGA B trisaccharide |
|---|---|---|
| HDX reaction details | 20 mM Tris, 150 mM NaCl, pH 2.3, 25 °C | |
| HDX time course (min) | 1, 10, 60, 480 | |
| HDX control samples | Fully deuterated protein control, labeled for 24 h, buffer 20 mM Tris, 6 M urea, pH 7 + FD model peptides in T9 | |
| Back-exchange (mean) | 50 ± 15 % (FD protein) | |
| # of Peptides | 126 | 126 |
| Sequence coverage | 89% | 89% |
| Average peptide length / Redundancy | 16.4 / 6.49 | 16.4 / 6.49 |
| Replicates (biological or technical) | 3 (technical) | 3 (technical) |
| Repeatability (average standard deviation) | 0.12 | 0.178 |
| Significant differences in HDX (delta HDX > X D) | T test (p<0.05) and delta D > 0.43 (2x pooled average SD), manual validation in case of bimodality | |

**T3** **GII.17 Kawasaki P dimer**

| Data Set | unbound | 100 mM fucose |
|---|---|---|
| HDX reaction details | 20 mM Tris, 150 mM NaCl, pH 2.3, 25 °C | |
| HDX time course (min) | 1, 10, 60, 480 | |
| HDX control samples | Fully deuterated protein control, labeled for 72 h, buffer 20 mM Tris, 6 M urea, pH 7 + FD model peptides in T9 | |
| Back-exchange (mean) | 64 ± 11 % (FD protein) | |
| # of Peptides | 89 | 89 |
| Sequence coverage | 97% | 97% |
| Average peptide length / Redundancy | 16.3 / 4.69 | 16.3 / 4.69 |
| Replicates (biological or technical) | 3 (technical) | 3 (technical) |
| Repeatability (average standard deviation) | 0.057 | 0.083 |
| Significant differences in HDX (delta HDX > X D) | T test (p<0.05) and delta D > 0.20 (2x pooled average SD), manual validation in case of bimodality | |

**T4** **GII.17 Kawasaki P dimer**

| Data Set | unbound | 10 mM HBGA B trisasccharide |
|---|---|---|
| HDX reaction details | 20 mM Tris, 150 mM NaCl, pH 2.3, 25 °C | |
| HDX time course (min) | 1, 10, 60, 480 | |
| HDX control samples | Fully deuterated protein control, labeled for 72 h, buffer 20 mM Tris, 6 M urea, pH 7 + FD model peptides in T9 | |
| Back-exchange (mean) | 58 ± 14 % (FD protein) | |
| # of Peptides | 84 | 84 |
| Sequence coverage | 97% | 97% |
| Average peptide length / Redundancy | 16.1 / 4.36 | 16.1 / 4.36 |
| Replicates (biological or technical) | 3 (technical) | 3 (technical) |
| Repeatability (average standard deviation) | 0.115 | 0.157 |
| Significant differences in HDX (delta HDX > X D) | T test (p<0.05) and delta D > 0.39 (2x pooled average SD), manual validation in case of bimodality | |

**T5** **GII.4 MI001 P dimer wildtype**

| Data Set | unbound | 10 mM HBGA B trisasccharide |
|---|---|---|
| HDX reaction details | 20 mM Tris, 150 mM NaCl, pH 2.3, 25 °C | |
| HDX time course (min) | 1, 10, 60, 480 | |
| HDX control samples | Fully deuterated protein control, labeled for 72 h, buffer 20 mM Tris, 6 M urea, pH 7 + FD model peptides in T9 | |
| Back-exchange (mean) | 47 ± 12 % (FD protein) | |
| # of Peptides | 100 | 100 |
| Sequence coverage | 96% | 96% |
| Average peptide length / Redundancy | 16.9 / 5.43 | 16.9 / 5.43 |
| Replicates (biological or technical) | 3 (technical) | 3 (technical) |
| Repeatability (average standard deviation) | 0.08 | 0.073 |
| Significant differences in HDX (delta HDX > X D) | T test (p<0.05) and delta D > 0.22 (2x pooled average SD), manual validation in case of bimodality | |
| **Data Set** | **unbound** | **100 mM fucose** |
| HDX reaction details | 20 mM Tris, 150 mM NaCl, pH 2.3, 25 °C | |
| HDX time course (min) | 1, 10, 60, 480 | |
| HDX control samples | Fully deuterated protein control, labeled for 72 h, buffer 20 mM Tris, 6 M urea, pH 7 + FD model peptides in T9 | |
| Back-exchange (mean) | 47 ± 12 % (FD protein) | |
| # of Peptides | 100 | 100 |
| Sequence coverage | 96% | 96% |
| Average peptide length / Redundancy | 16.9 / 5.43 | 16.9 / 5.43 |
| Replicates (biological or technical) | 3 (technical) | 3 (technical) |
| Repeatability (average standard deviation) | 0.08 | 0.09 |
| Significant differences in HDX (delta HDX > X D) | T test (p<0.05) and delta D > 0.23 (2x pooled average SD), manual validation in case of bimodality | |

**T6** **GII.4 MI001 P dimer wildtype**

| Data Set | unbound | 100 mM fucose |
|---|---|---|
| HDX reaction details | 20 mM Tris, 150 mM NaCl, pH 2.3, 25 °C | |
| HDX time course (min) | 0.25, 1, 10, 60, 480 | |
| HDX control samples | Fully deuterated protein control, labeled for 24 h, buffer 20 mM Tris, 6 M urea, pH 7 + FD model peptides in T9 | |
| Back-exchange (mean) | 40 ± 8 % (FD protein) | |
| # of Peptides | 81 | 81 |
| Sequence coverage | 87% | 87% |
| Average peptide length / Redundancy | 16.6 / 4.34 | 16.6 / 4.34 |
| Replicates (biological or technical) | 1 (technical) | 1 (technical) |
| Repeatability (average standard deviation) | 0.1 for FD (2 technical replicates) | - |
| Significant differences in HDX (delta HDX > X D) | delta D > 0.5 (99% percentile of dataset T5 with similar SD), manual validation in case of bimodality | |

**T7** **GII.4 MI001 P dimer partially deamidated**

| Data Set | unbound | 100 mM fucose |
|---|---|---|
| HDX reaction details | 20 mM Tris, 150 mM NaCl, pH 2.3, 25 °C | |
| HDX time course (min) | 1, 10, 60, 480 | |
| HDX control samples | Fully deuterated protein control, labeled for 72 h, buffer 20 mM Tris, 6 M urea, pH 7 + FD model peptides in T9 | |
| Back-exchange (mean) | 41 ± 13 % (FD protein) | |
| # of Peptides | 123 | 123 |
| Sequence coverage | 96% | 96% |
| Average peptide length / Redundancy | 15.0 / 5.94 | 15.0 / 5.94 |
| Replicates (biological or technical) | 3 (technical) | 3 (technical) |
| Repeatability (average standard deviation) | 0.074 | 0.079 |
| Significant differences in HDX (delta HDX > X D) | T test (p<0.05) and delta D > 0.21 (2x pooled average SD), manual validation in case of bimodality | |

**T8** **GII.4 MI001 P dimer wildtype vs. partially deamidated**

| Data Set | unbound (wt) (T5) | unbound (deam) (T7) |
|---|---|---|
| HDX reaction details | 20 mM Tris, 150 mM NaCl, pH 2.3, 25 °C | |
| HDX time course (min) | 1, 10, 60, 480 | |
| HDX control samples | fully deuterated protein control (FD T5 and T7) + FD model peptides in T9 | |
| Back-exchange (mean) | see T5 and T7 for FD protein | |
| # of Peptides | 69 | 69 |
| Sequence coverage | 95% | 95% |
| Average peptide length / Redundancy | 16.4 / 3.65 | 16.4 / 3.65 |
| Replicates (biological or technical) | 3 (technical) | 3 (technical) |
| Repeatability (average standard deviation) | 0.081 | 0.072 |
| Significant differences in HDX (delta HDX > X D) | delta D normalized with FD controls ratio, T test (p<0.05) and delta D > 0.42 (99% percentile) | |

**T9** **fully deuterated model peptides for HDX workflow quality control**

| Data Set | Angiotensin I | Bradykinin |
|---|---|---|
| HDX reaction details | 20 mM deuterated Tris, 6 M GndHCl, pH 2.3, 25 °C, labeled for 24 h | |
| HDX time course (min) | - | |
| HDX control samples | fully deuterated peptide mix serves as general back exchange control for the standard bottom-up HDX MS workflow | |
| Back-exchange (mean) | 23 ± 1% (FD peptide) | 32 ± 2 % (FD peptide) |
| # of Peptides | 1 | 1 |
| Sequence coverage | 100% | 100% |
| Average peptide length / Redundancy | 10 | 9 |
| Replicates (biological or technical) | 2 (technical) | 2 (technical) |
| Repeatability (average standard deviation) | 0.1 | 0.08 |
| Significant differences in HDX (delta HDX > X D) | - | |

**T10** **GII.4 Saga P dimer partially deamidated**

| Data Set | unbound | 100 mM fucose |
|---|---|---|
| HDX reaction details | 20 mM Tris, 150 mM NaCl, pH 2.3, 25 °C | |
| HDX time course (min) | 0.25. 1, 10 | |
| HDX control samples | Fully deuterated protein control, labeled for 24 h, buffer 20 mM Tris, 6 M urea, pH 7 + FD model peptides in T9 | |
| Back-exchange (mean) | 37 ± 9 % (FD protein) | |
| # of Peptides | 78 | 78 |
| Sequence coverage | 97% | 97% |
| Average peptide length / Redundancy | 18.2 / 4.59 | 18.2 / 4.59 |
| Replicates (biological or technical) | 3 (technical) | 3 (technical) |
| Repeatability (average standard deviation) | 0.084 | 0.109 |
| Significant differences in HDX (delta HDX > X D) | T test (p<0.05) and delta D > 0.27 (2x pooled average SD) | |

### 7) MS-Viewer search keys

Table S 4: MS-Viewer search keys. Annotated fragment ion spectra of identified peptides can be viewed in the MS-Viewer online tool at http://msviewer.ucsf.edu/prospector/cgi-bin/msform.cgi?form=msviewer

| Protein | Deamidation status | MS-Viewer search key |
|---|---|---|
| GII.4 MI001 P domain | wildtype | widghx5vtx |
| GII.4 MI001 P domain | partially deamidated | svwlky0tqx |
| GII.4 Saga P domain | partially deamidated | adolnt1njh |
| GII.10 Vietnam 026 | wildtype | ycj8werllz |
| GII.17 Kawasaki 308 | wildtype | 9n5zuotk1a |

### 8) MD simulations

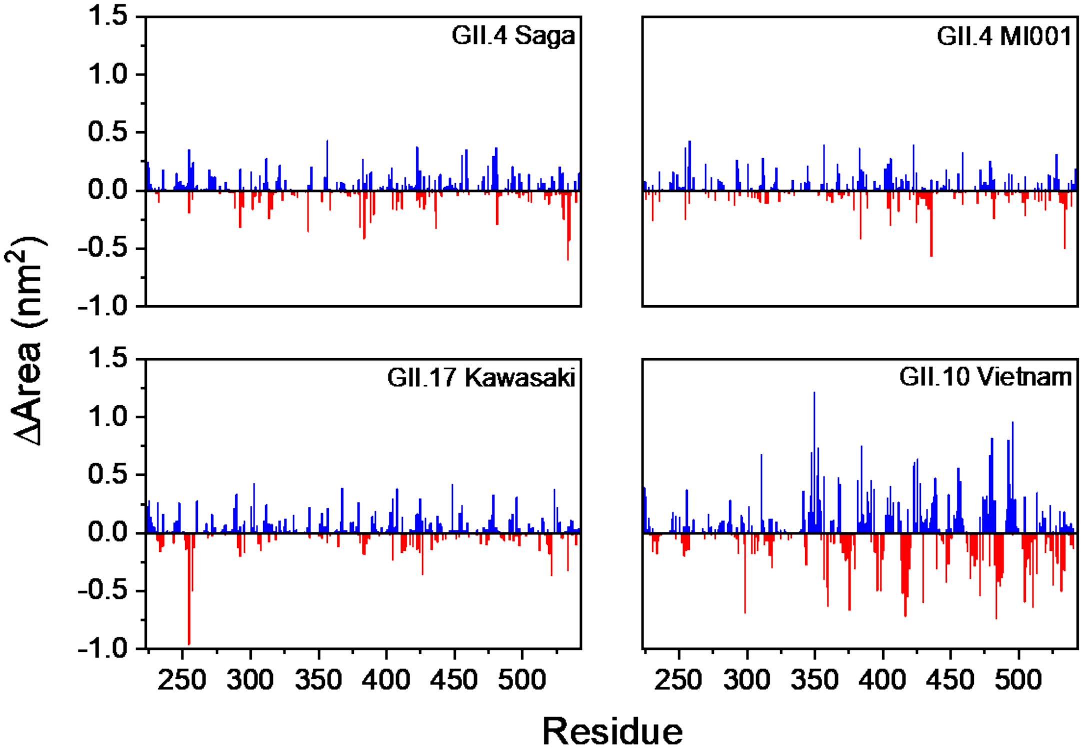


Figure S 17: Solvent accessible surface area calculations for GII.4 Saga, GII.4 MI001, GII.17 Kawasaki and GII.10 Vietnam. Blue peaks describe an increase of surface area per residue, whilst red peaks represent a decrease of the area throughout the simulations as compared to the initial structure.

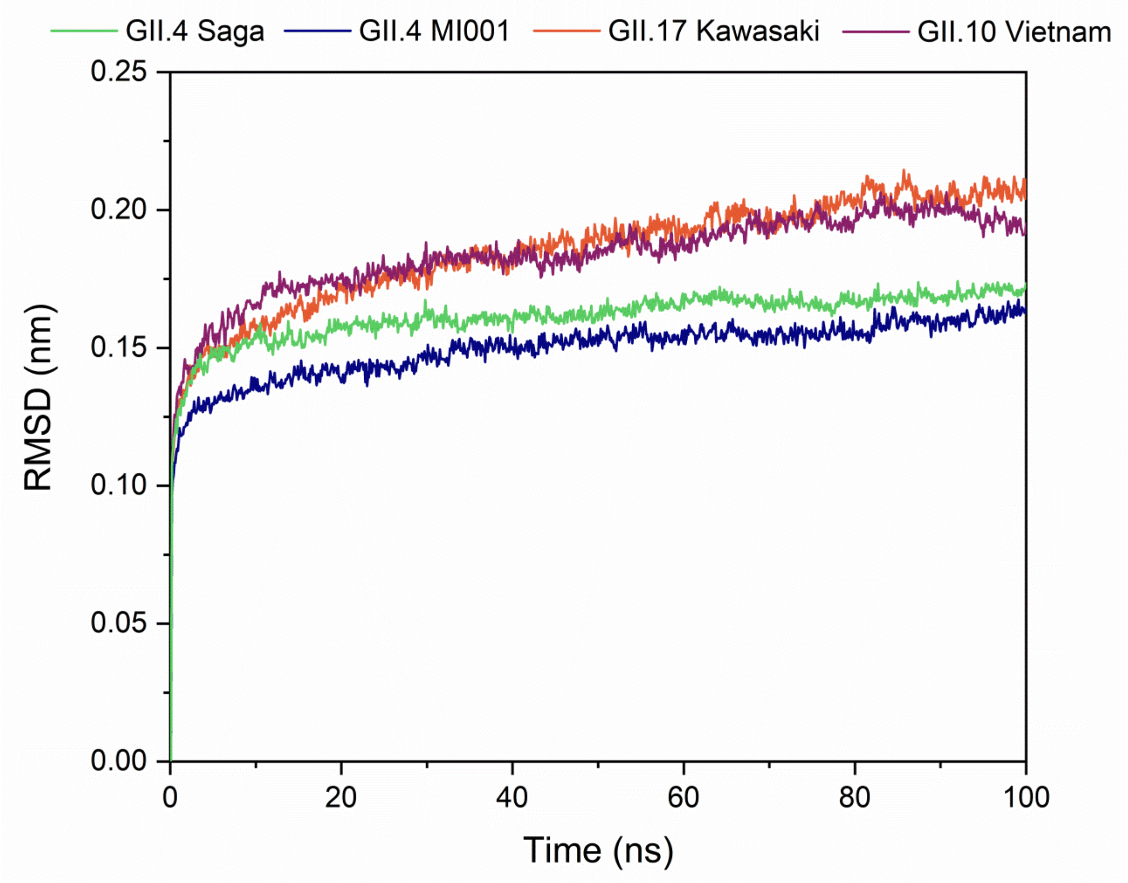


Figure S 18: RMSD of the GII.4 Saga, GII.4 MI001, GII.17 Kawasaki and GII.10 Vietnam P dimer strains.

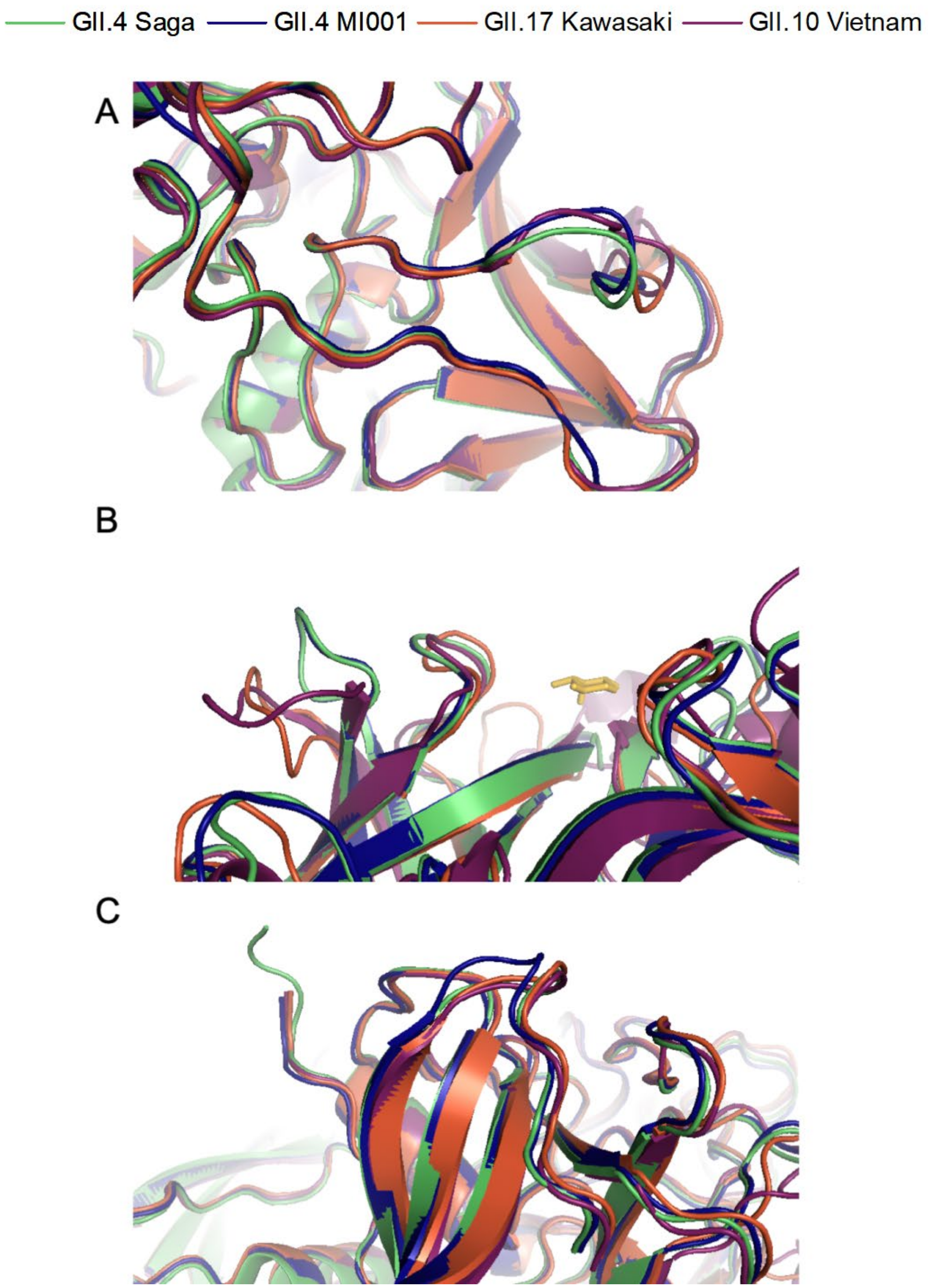


Figure S 19: Overlaid peptide chains of GII.4 Saga, GII.4 MI001, GII.17 Kawasaki and GII.10 Vietnam reveal differences in the peptide chains around residue 250 (A), 300 (B) and 424 (C). These areas of interest were chosen according to higher fluctuation observed in the RMSF data (figure 5 in the main manuscript). Unlike the loop around residue 350 (figure 6 in the main manuscript), which features a small helix in the GII.10 Vietnam strain, conformations depicted in this image show only smaller differences between the different strains. Fucose can be observed in panel B yellow-colored.

A

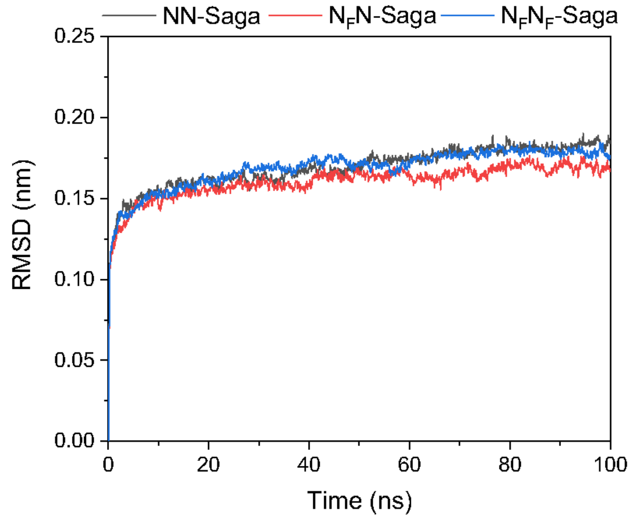
NN-Saga
$N_FN$-Saga
$N_FN_F$-Saga
0.25
0.20
0.15
0.10
0.05
0.00
RMSD (nm)
0
20
40
60
80
100
Time (ns)

B

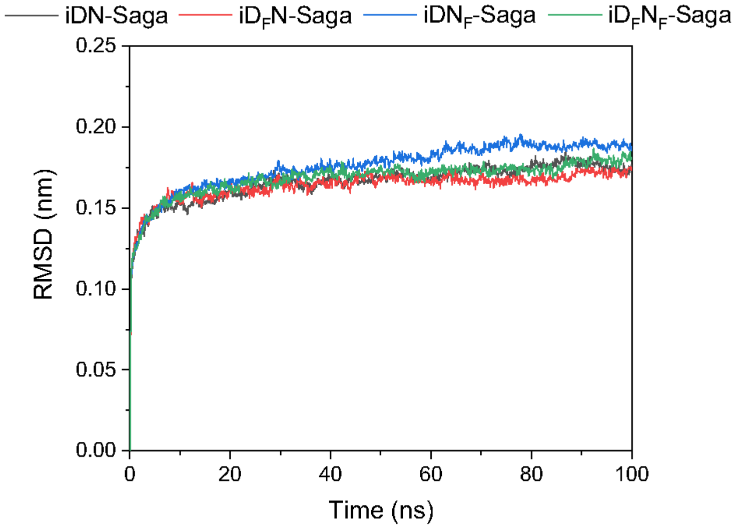
iDN-Saga
$iD_FN$-Saga
$iDN_F$-Saga
$iD_FN_F$-Saga
0.25
0.20
0.15
0.10
0.05
0.00
RMSD (nm)
0
20
40
60
80
100
Time (ns)

C

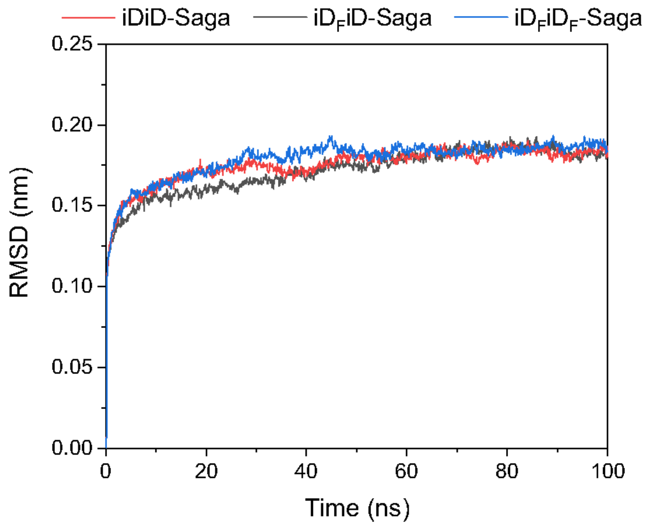
iDiD-Saga
$iD_FiD$-Saga
$iD_FiD_F$-Saga
0.25
0.20
0.15
0.10
0.05
0.00
RMSD (nm)
0
20
40
60
80
100
Time (ns)

Figure S 20: RMSD of GII.4 Saga NN (A), iDN (B) and iDiD (C) P dimers, pristine and complexed by fucose.

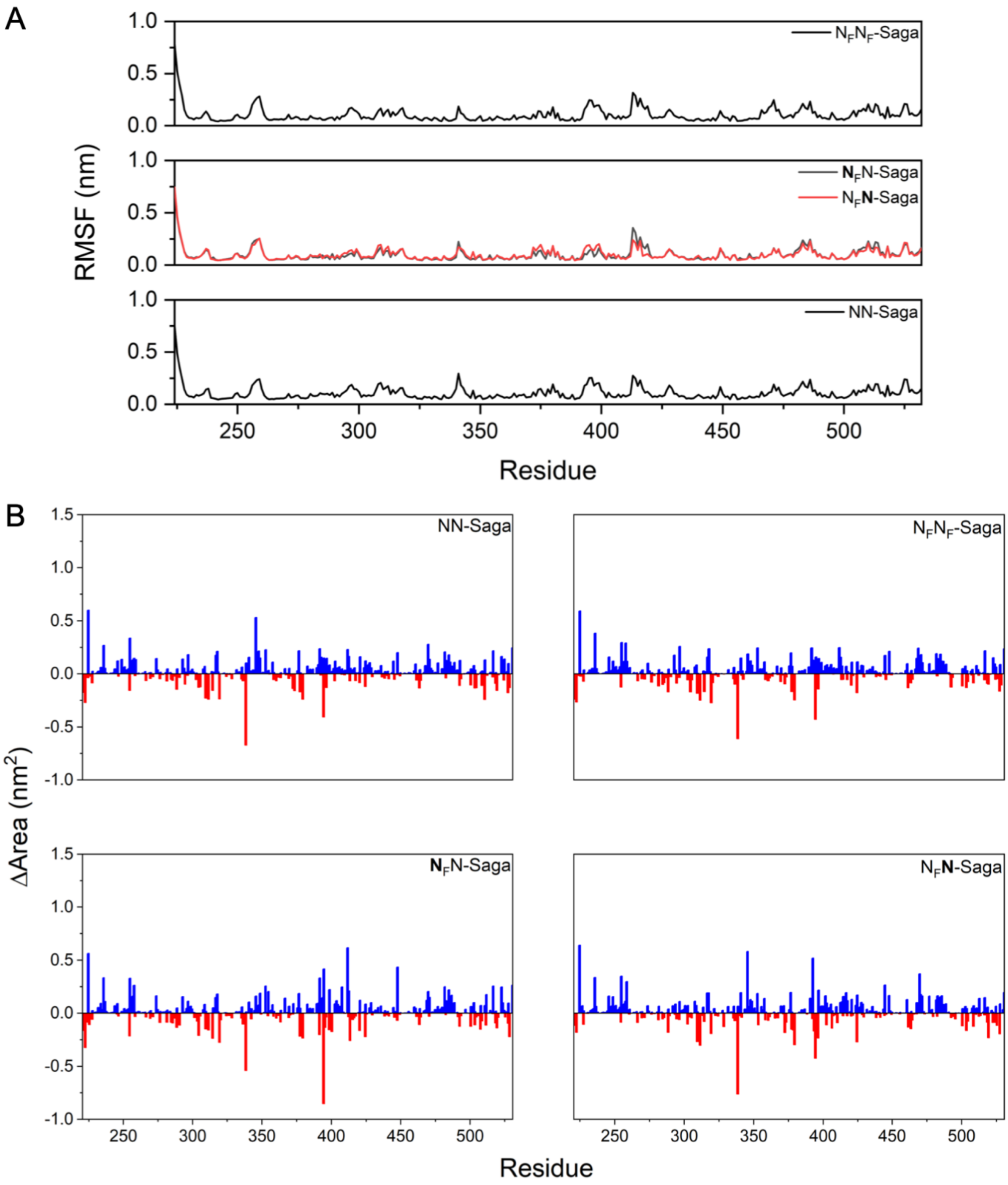


Figure S 21: RMSF (A) and the solvent accessible surface area (B) of NN-Saga without and complexed by fucose. Bold characters in the legends in panel B relate to the protein chain presented by the data.

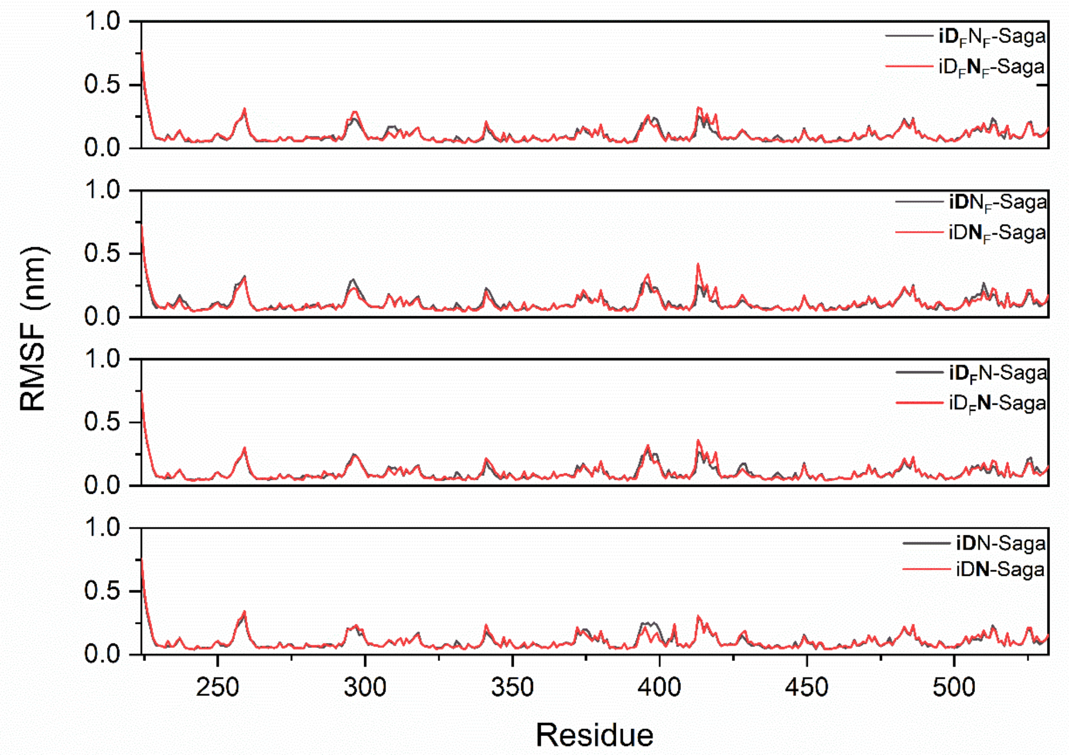


Figure S22: RMSF (A) and the solvent accessible surface area (B) of iDN-Saga without and complexed by fucose. Bold characters in the legends in panel B relate to the protein chain presented by the data.

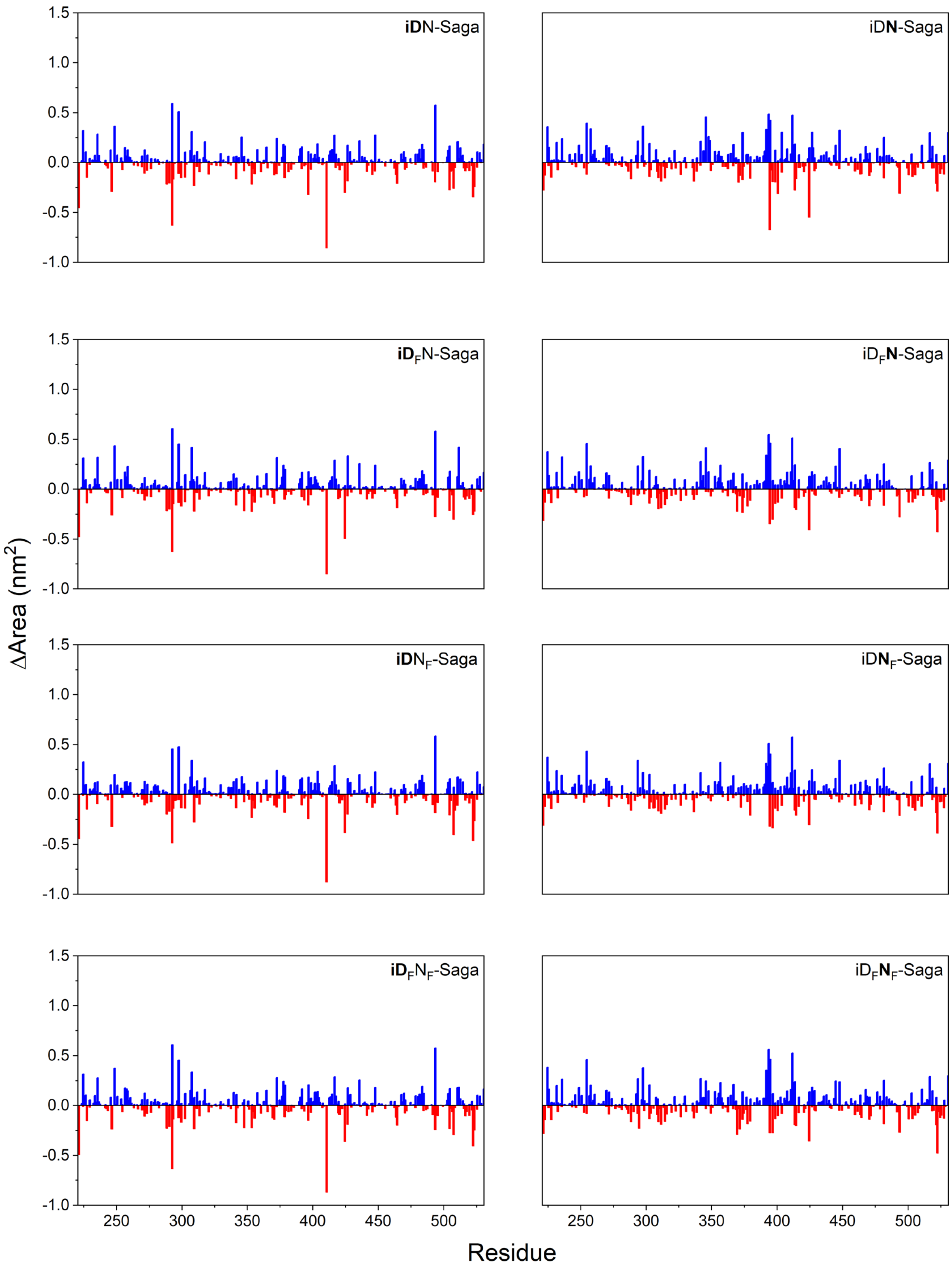


Figure S 22: continued.

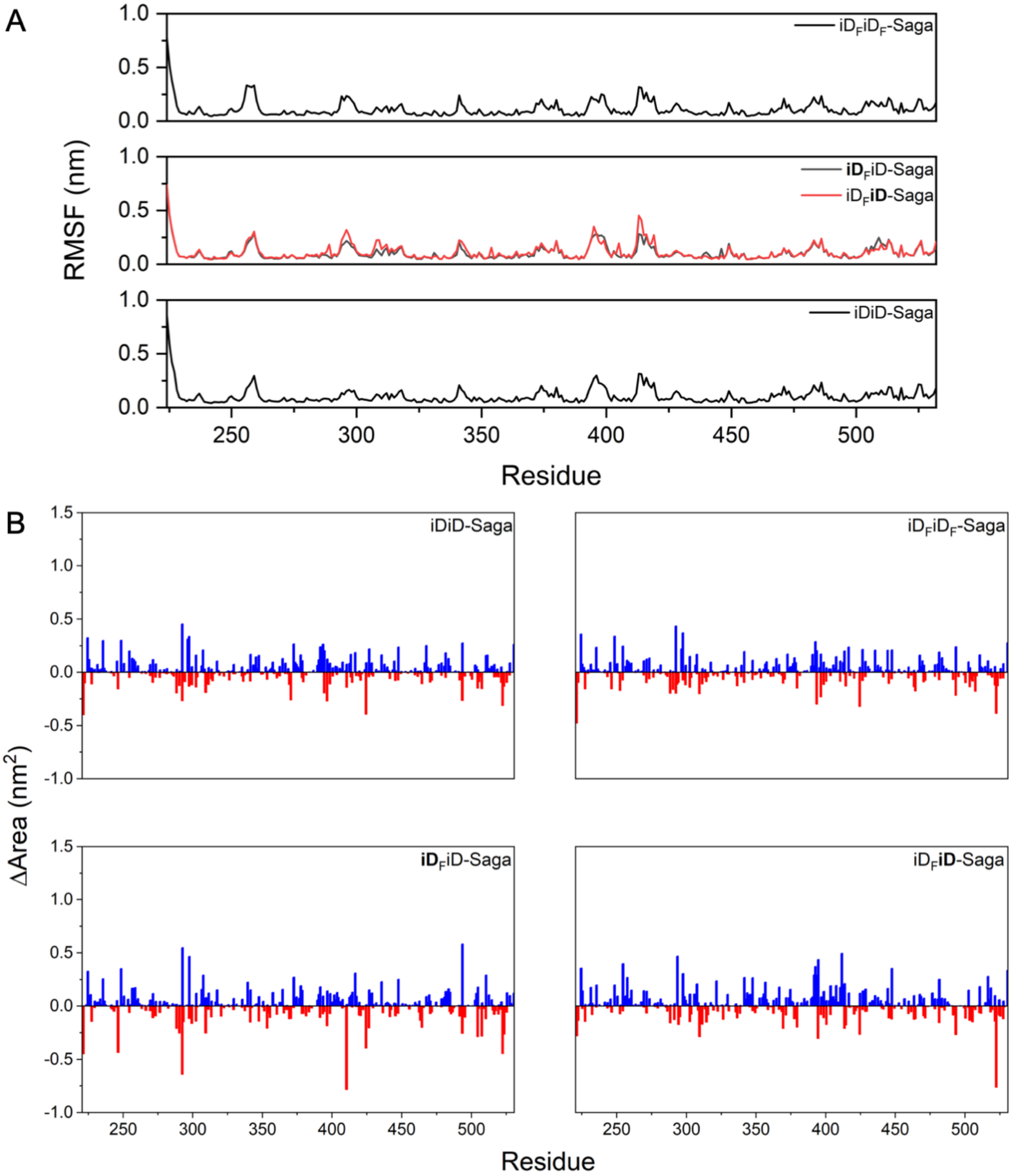


Figure S 23: RMSF (A) and the solvent accessible surface area (B) of iDiD-Saga without and complexed by fucose. Bold characters in the legends in panel B relate to the protein chain presented by the data.

### 9) Deuterium uptake plots

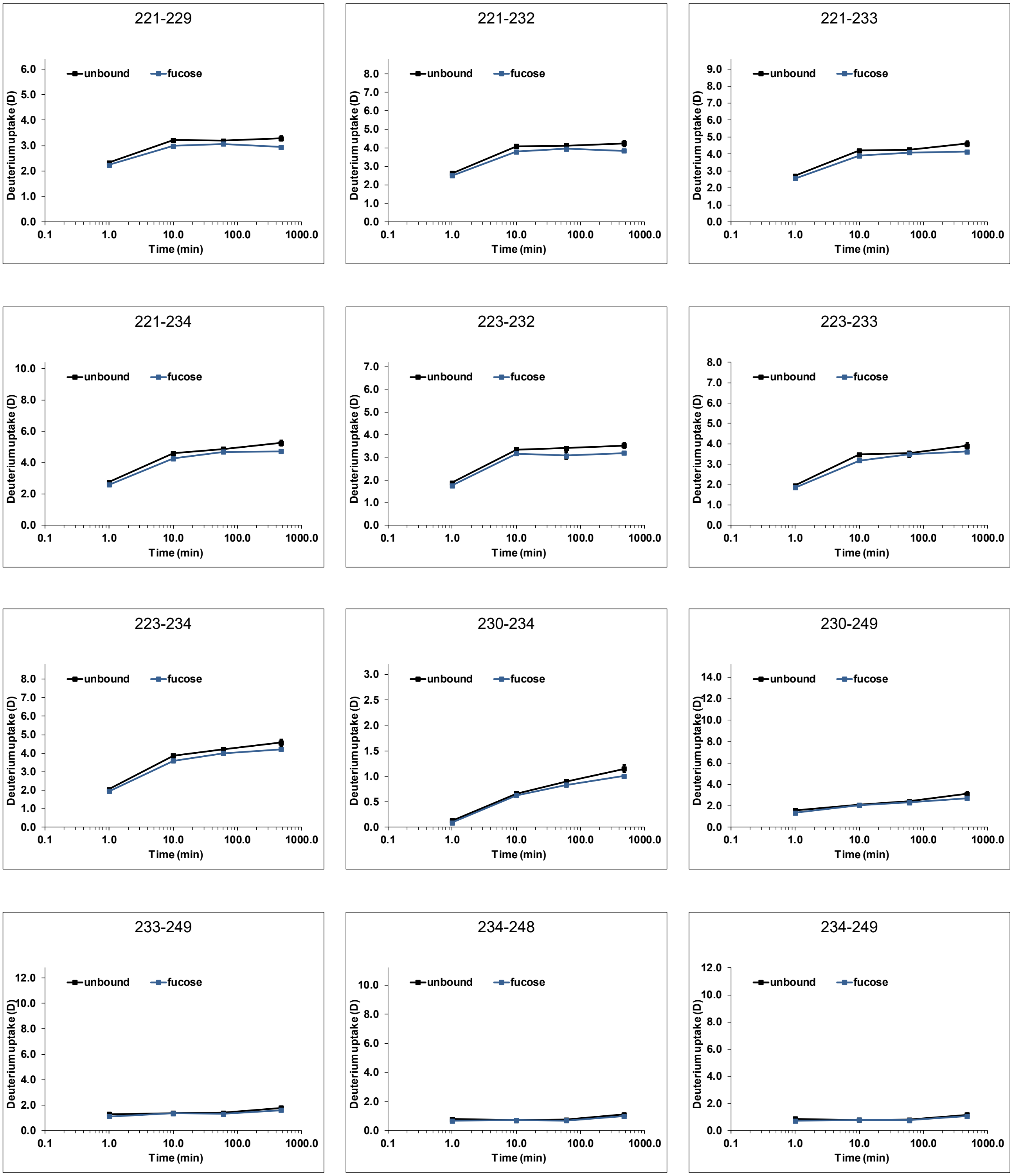


Figure S 24: Deuterium uptake plots for Vietnam P dimers (black) with 100 mM fucose (blue) (dataset T1). Time points 1 min, 10 min, and 1 h were performed in triplicate, the 8 h time point with fucose represents a single measurement. Error bars indicate the standard deviation (SD) of each triplicate analysis.

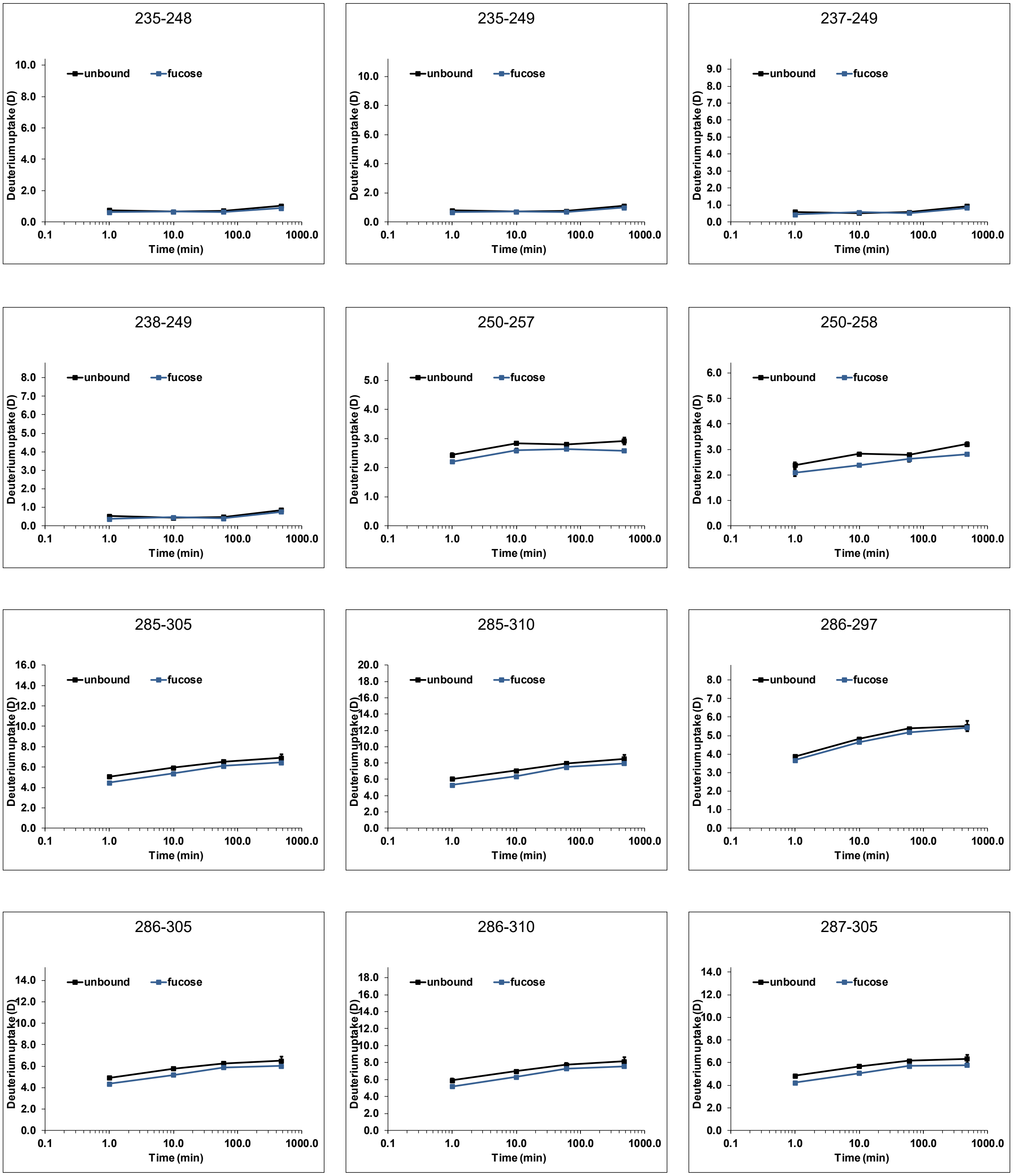


Figure S 24: continued

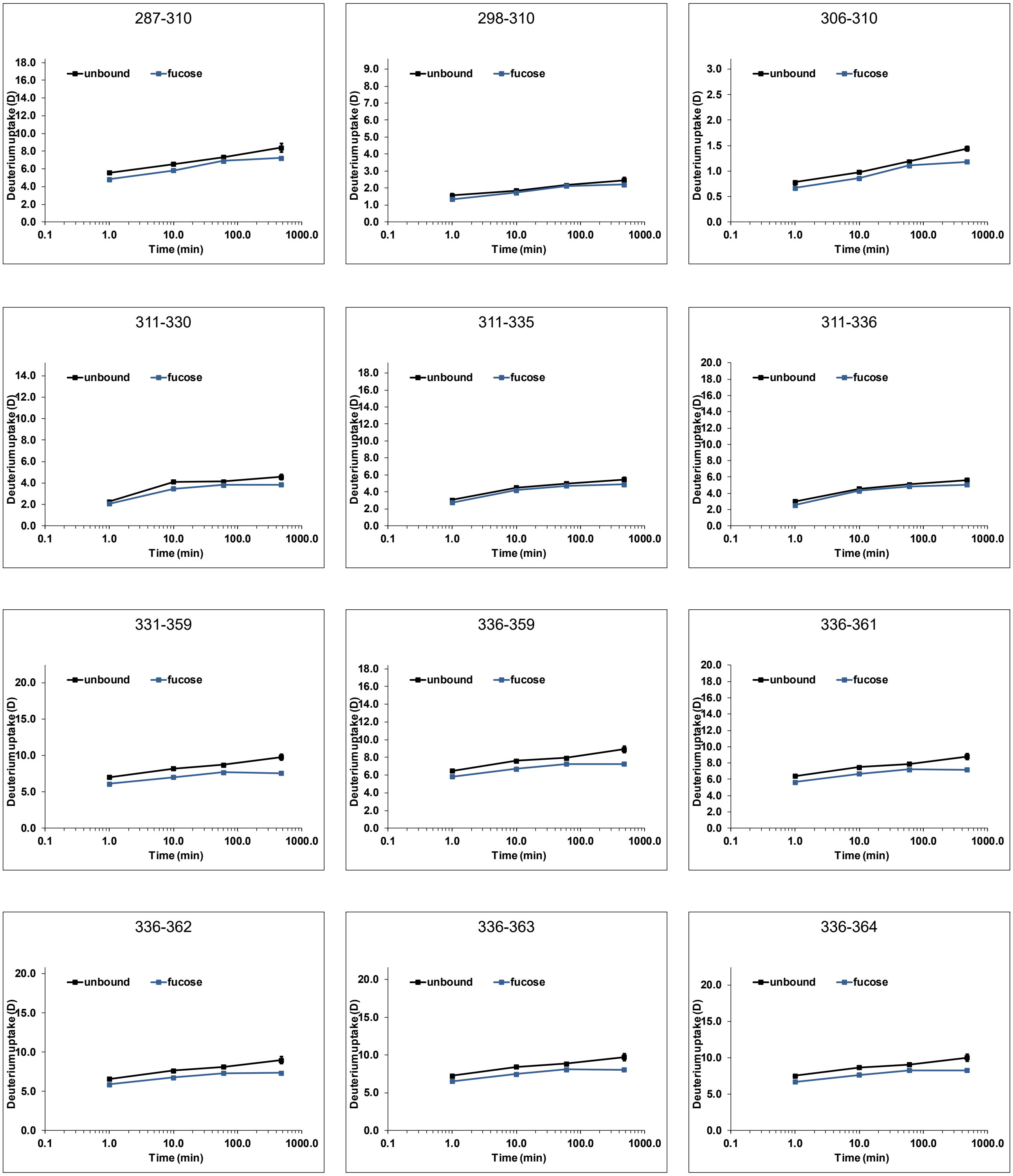


Figure S 24: continued

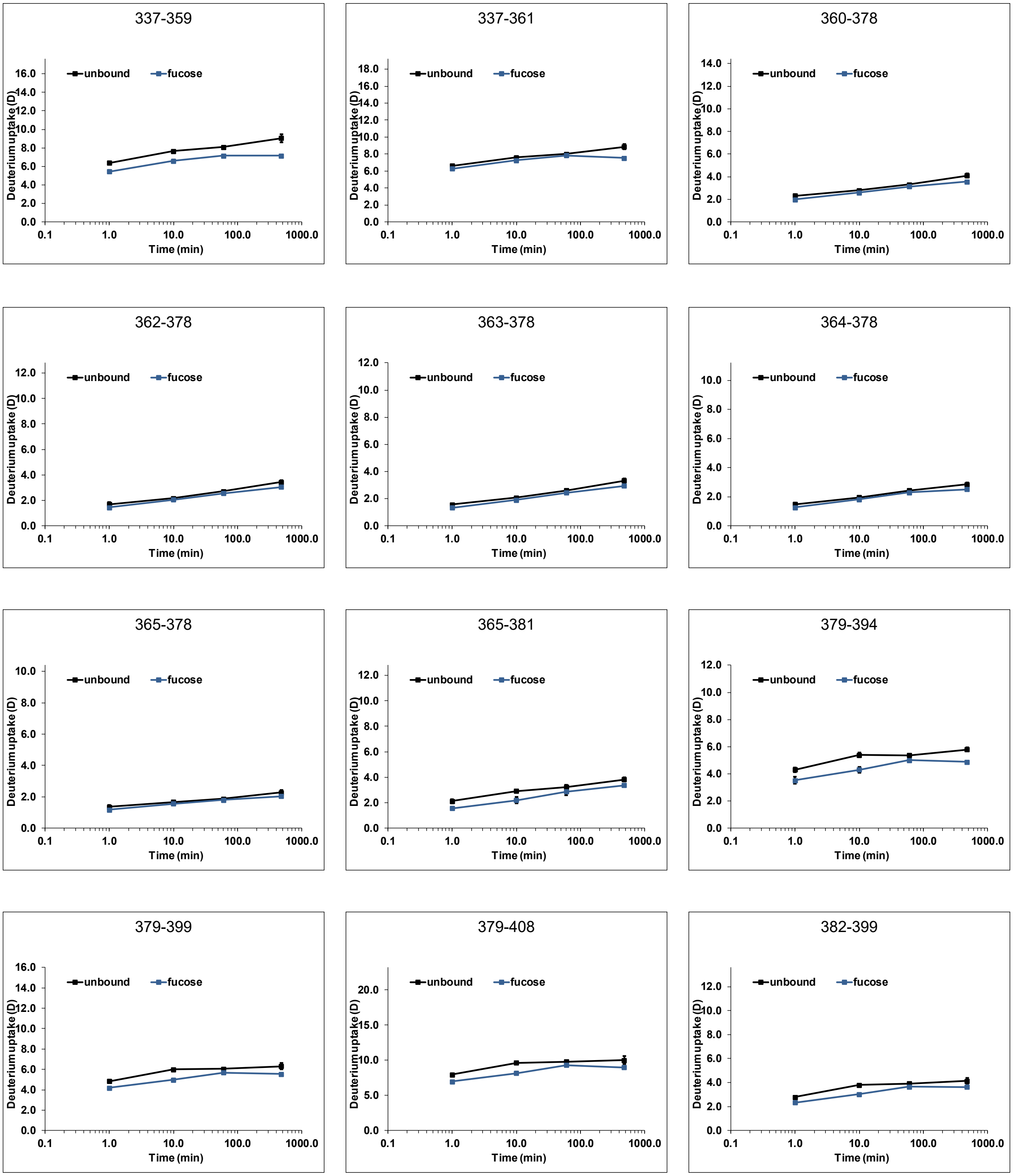


Figure S 24: continued

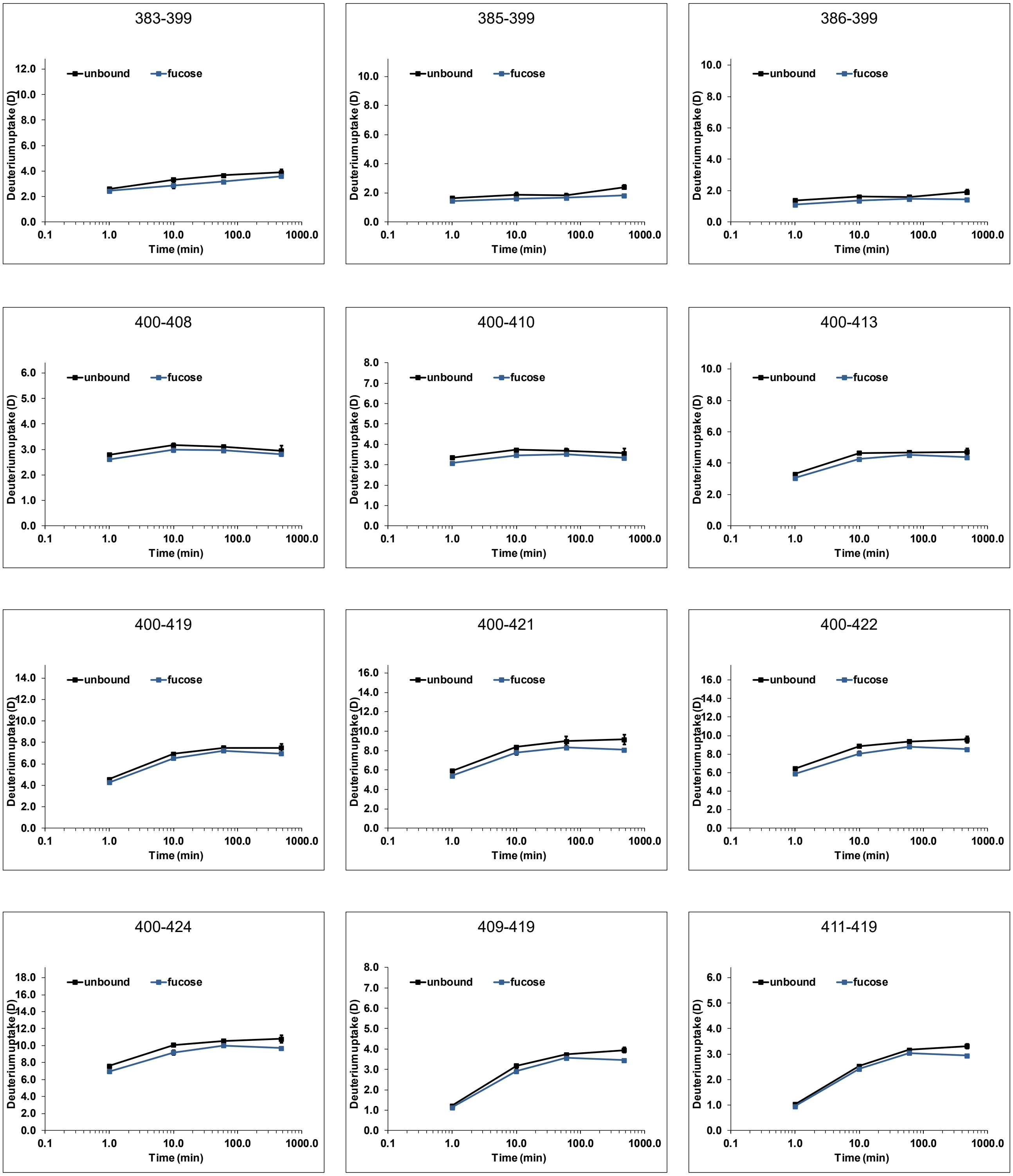


Figure S 24: continued

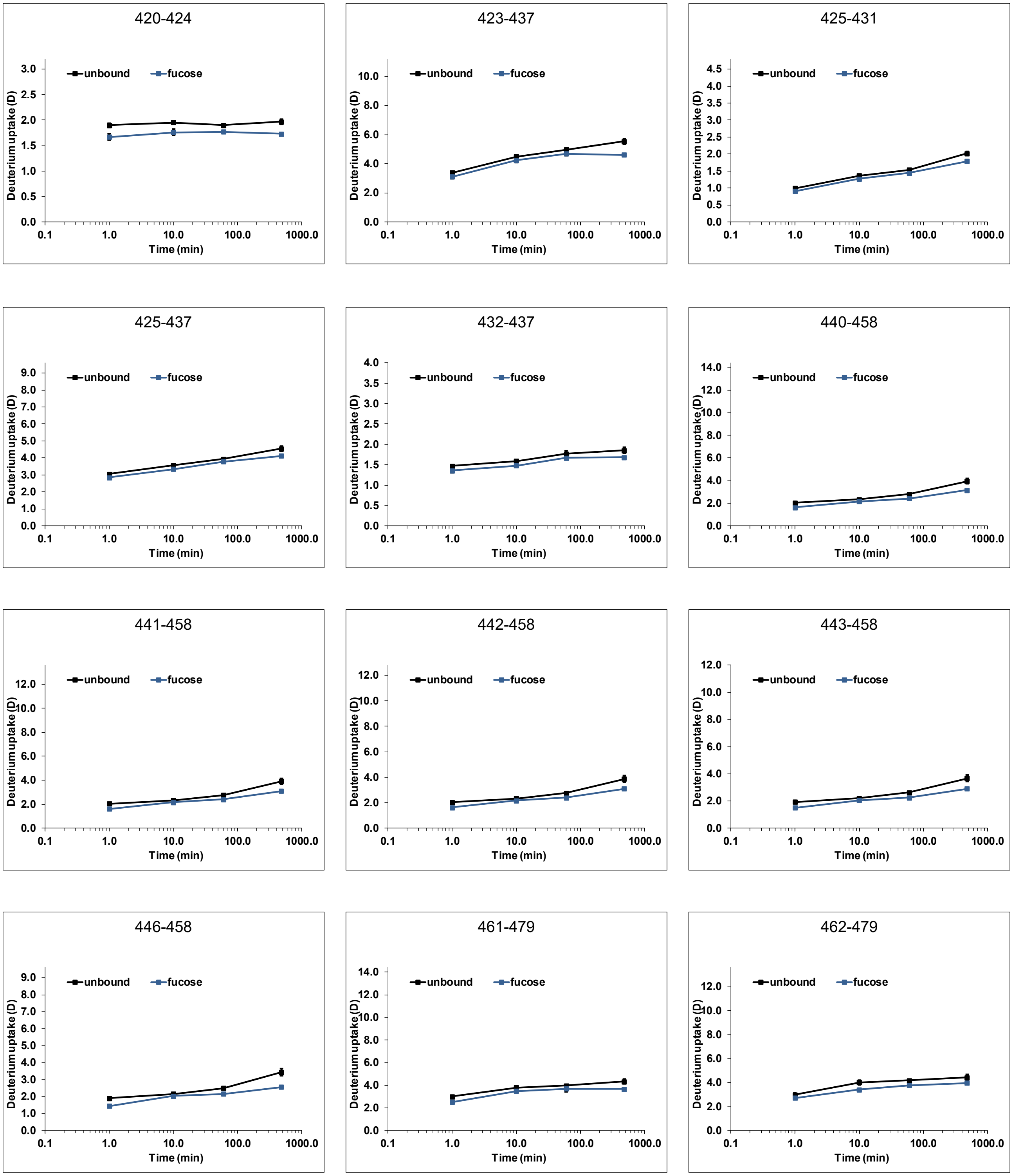

Figure S 24: continued

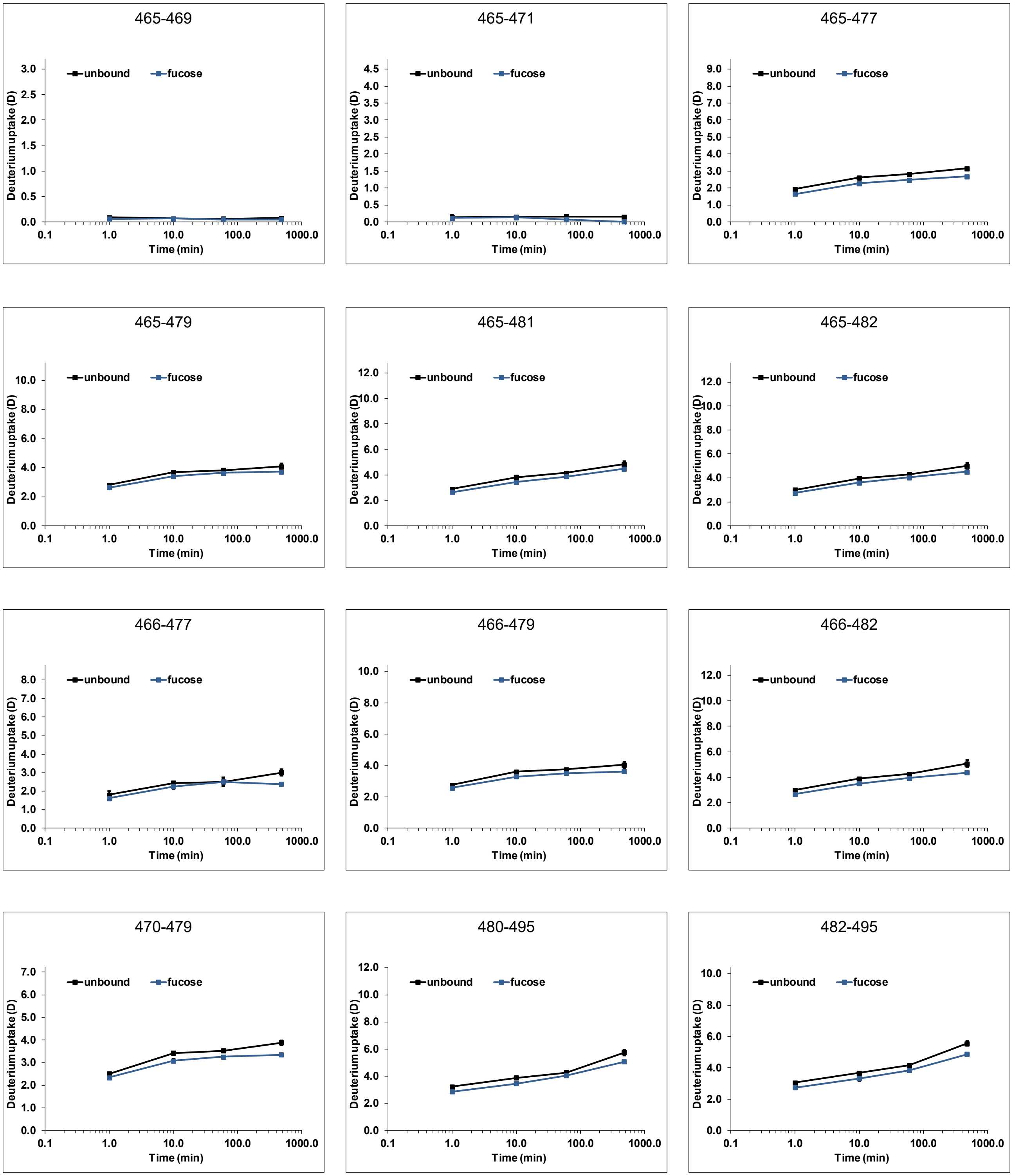


Figure S 24: continued

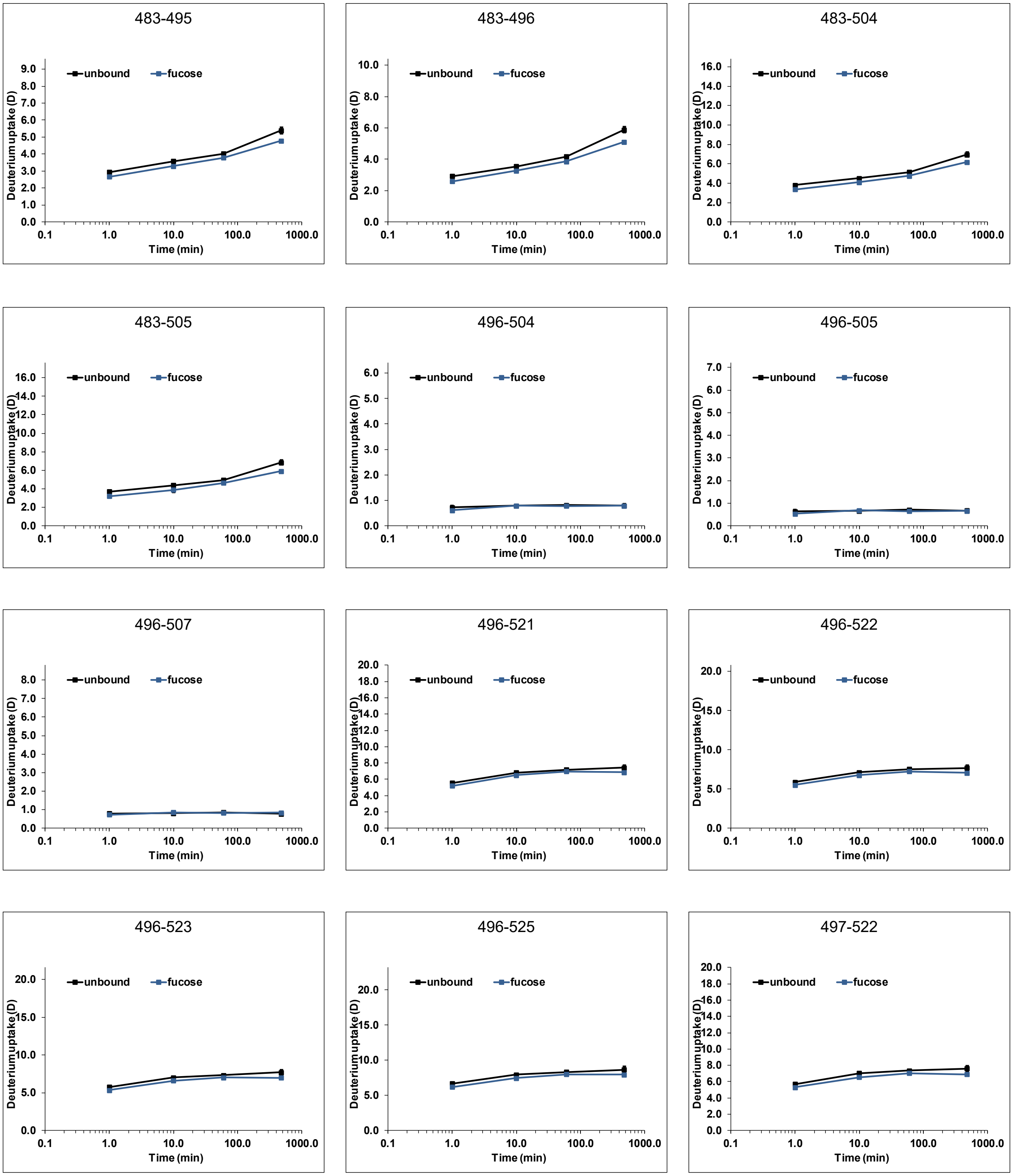

Figure S 24: continued

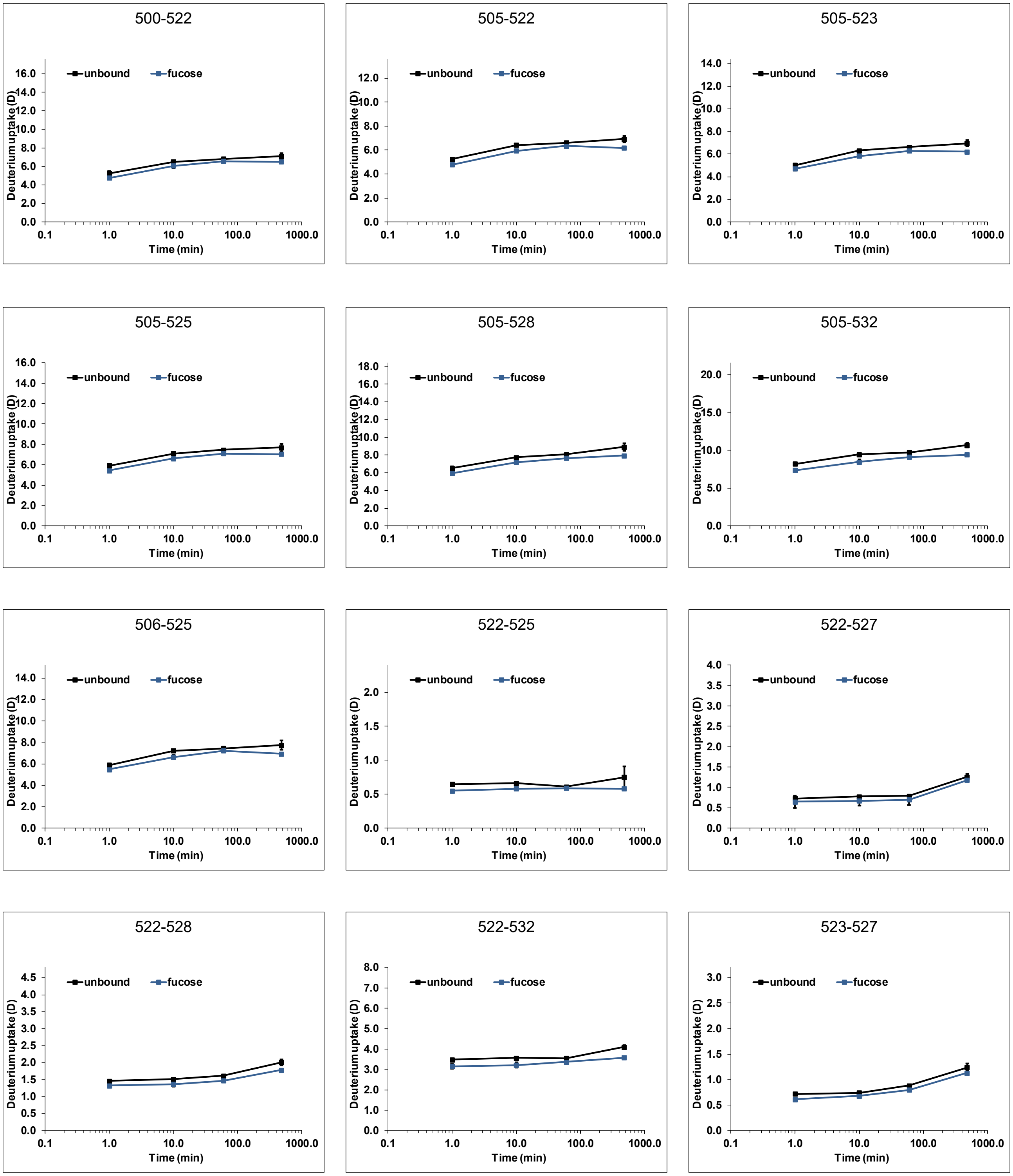

Figure S 24: continued

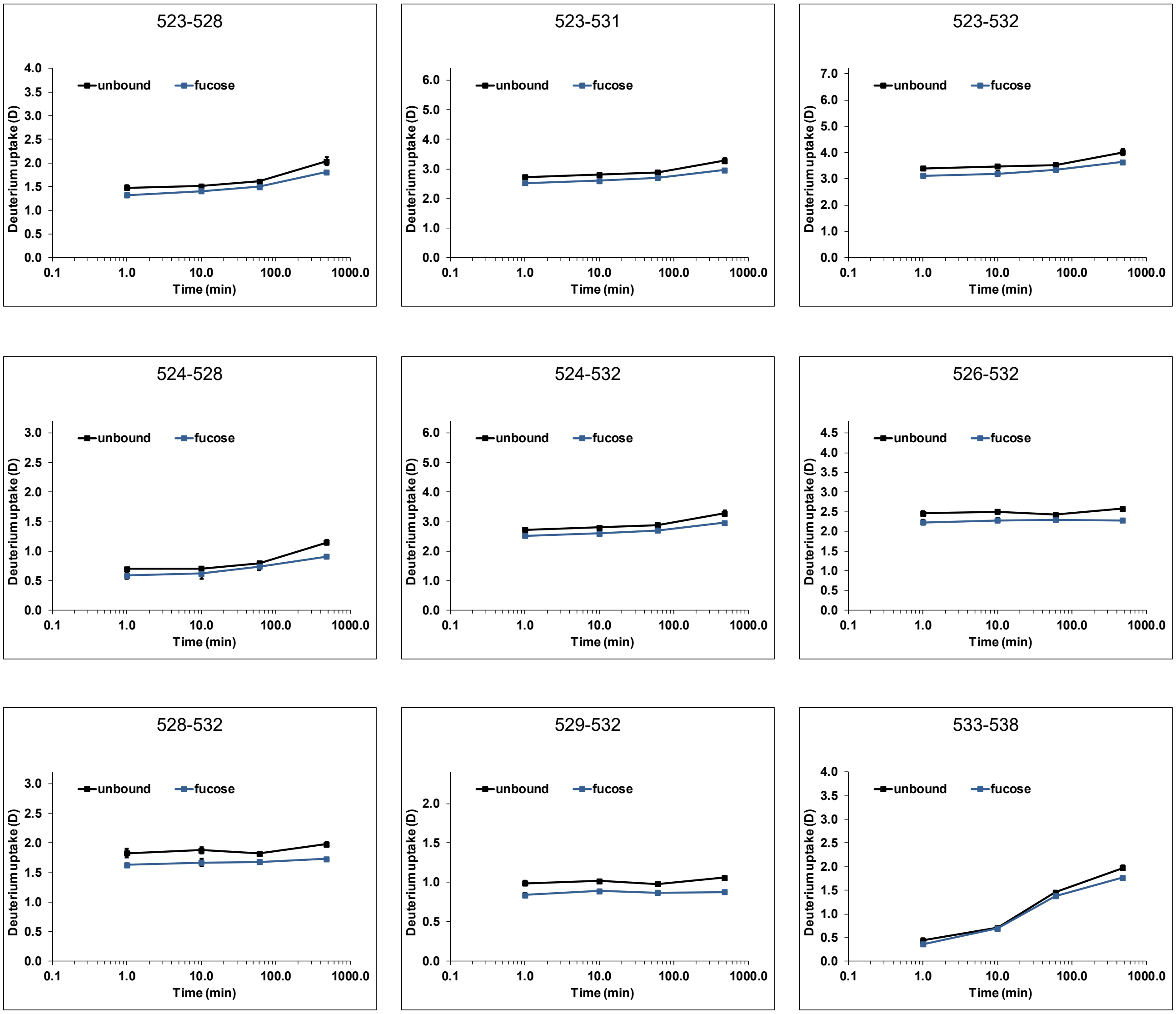


Figure S 24: continued

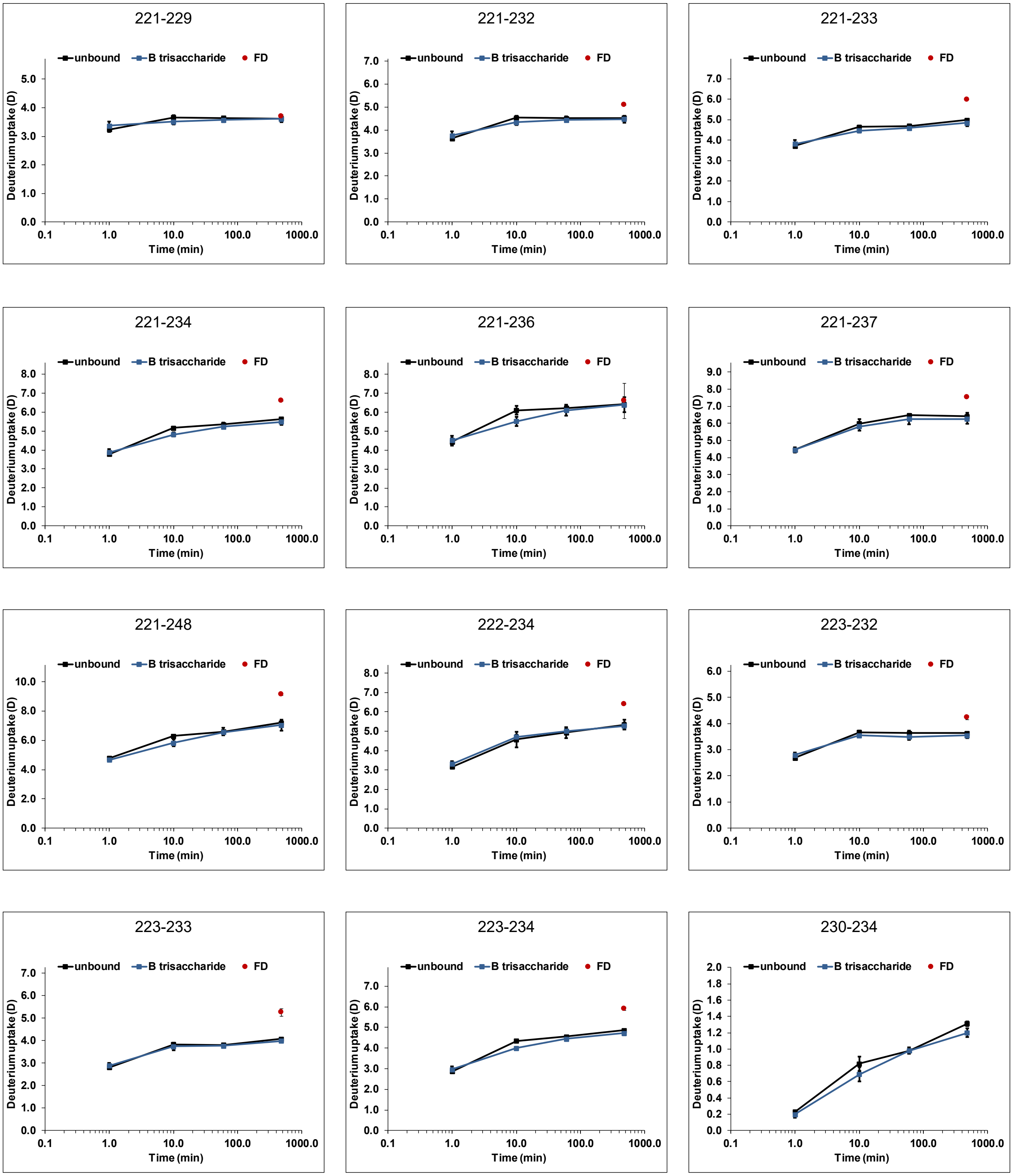


Figure S 25: Deuterium uptake plots for Vietnam P dimers (black) with 10 mM HBGA B trisaccharide (blue) (dataset T2). All time points (1 min, 10 min, 1 h and 8 h) were performed in triplicate, the fully deuterated control (FD, red) represents two measurements. Error bars indicate the standard deviation (SD).

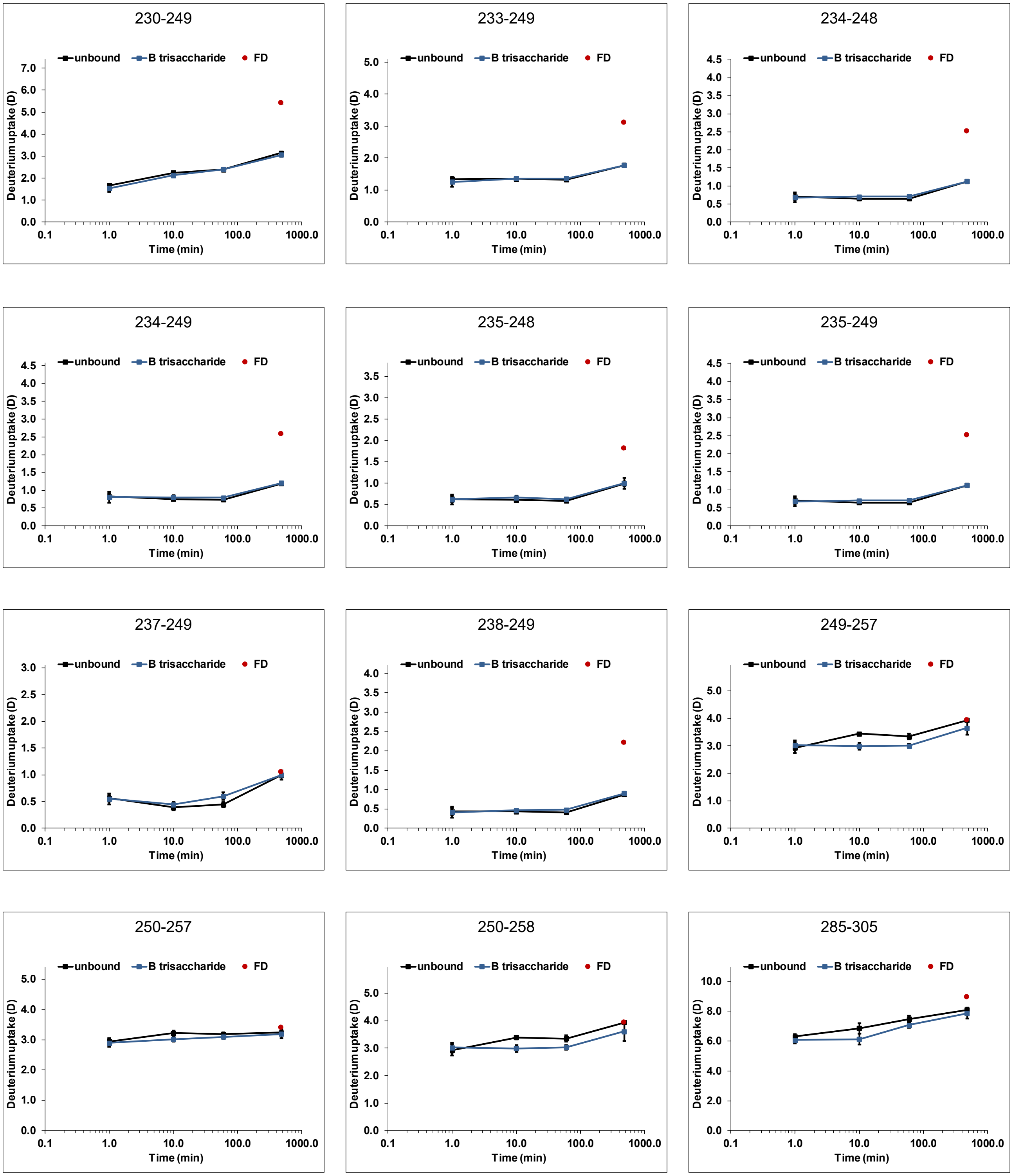


Figure S 25: continued

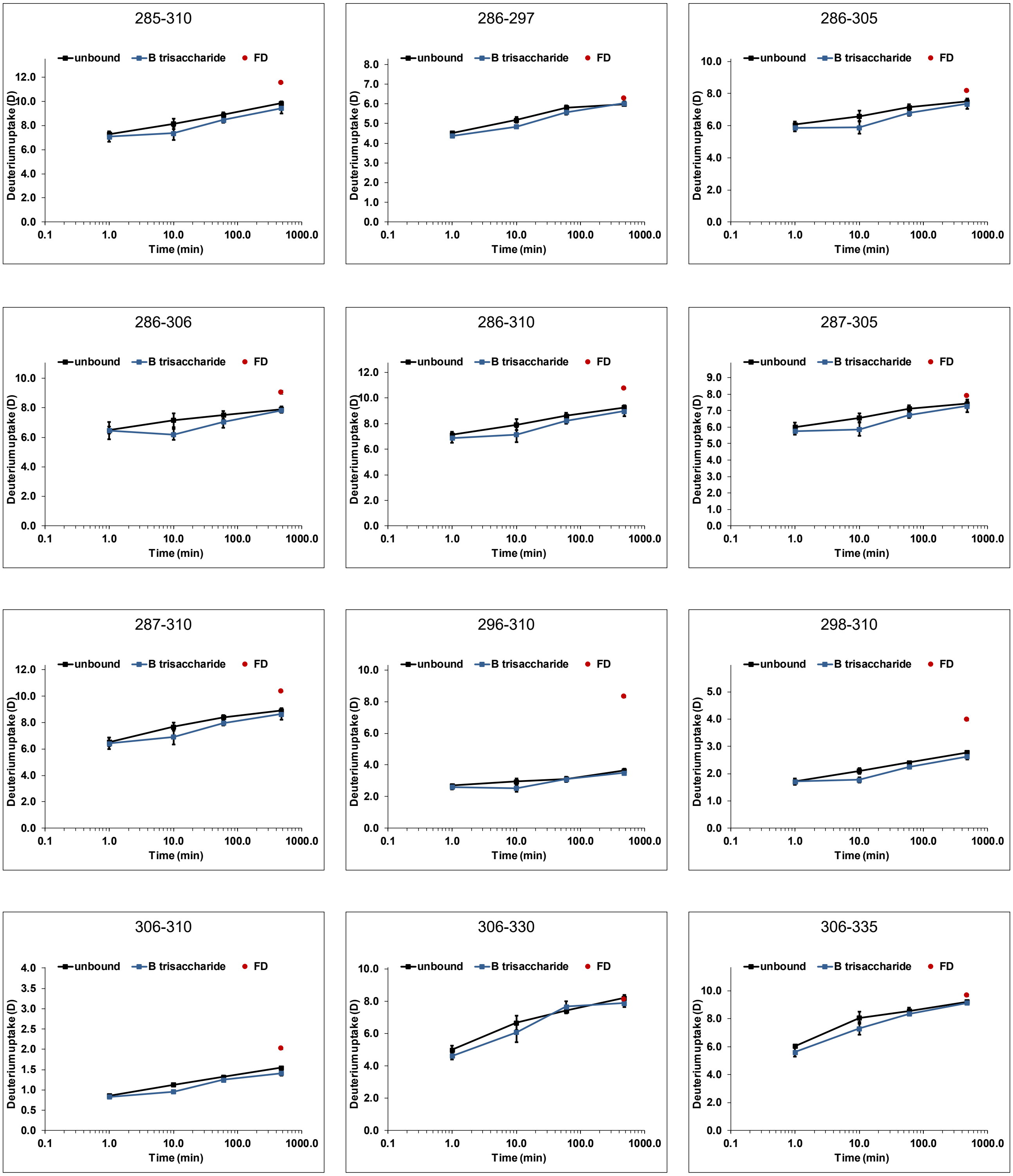

Figure S 25: continued

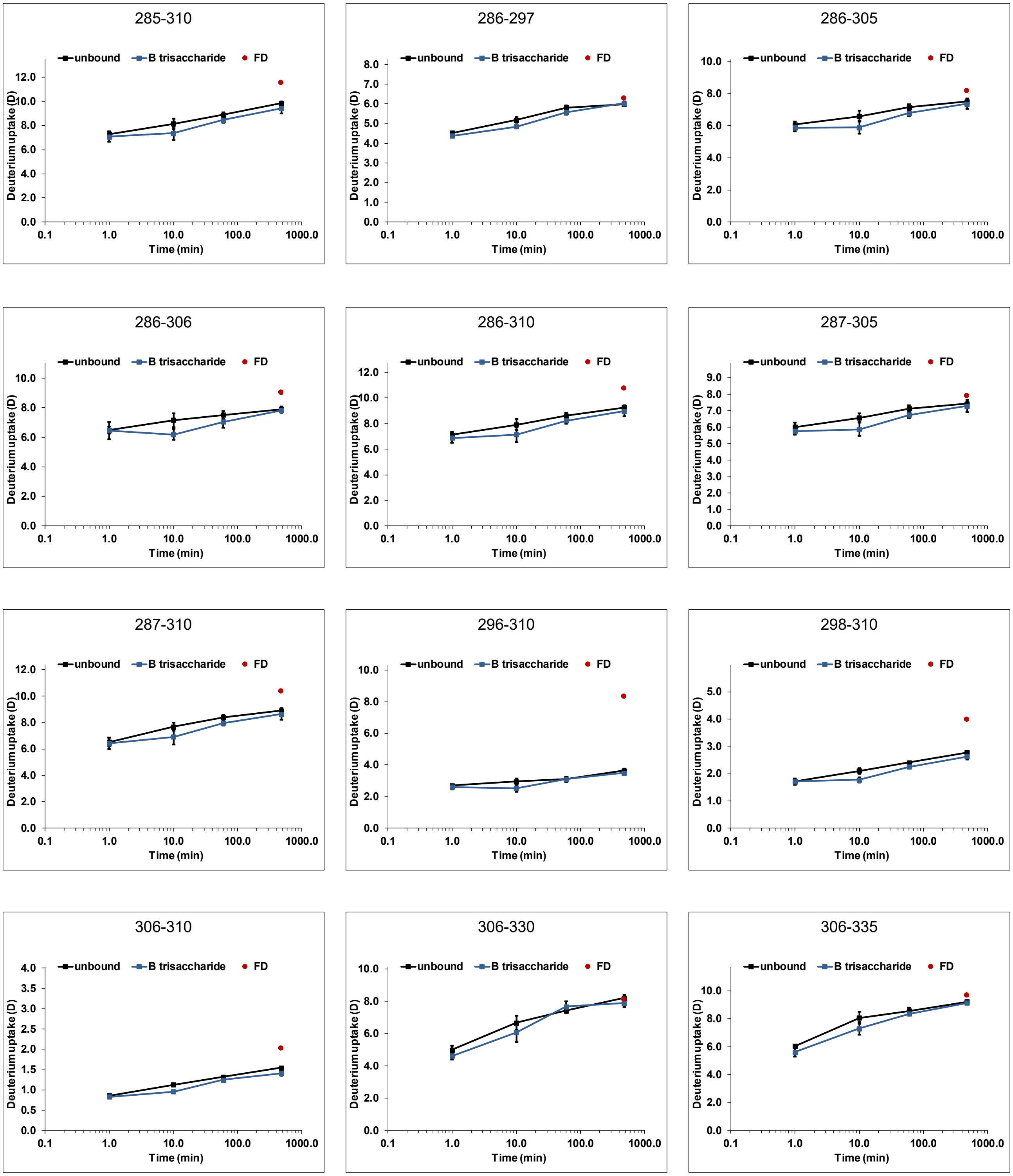

Figure S 25: continued

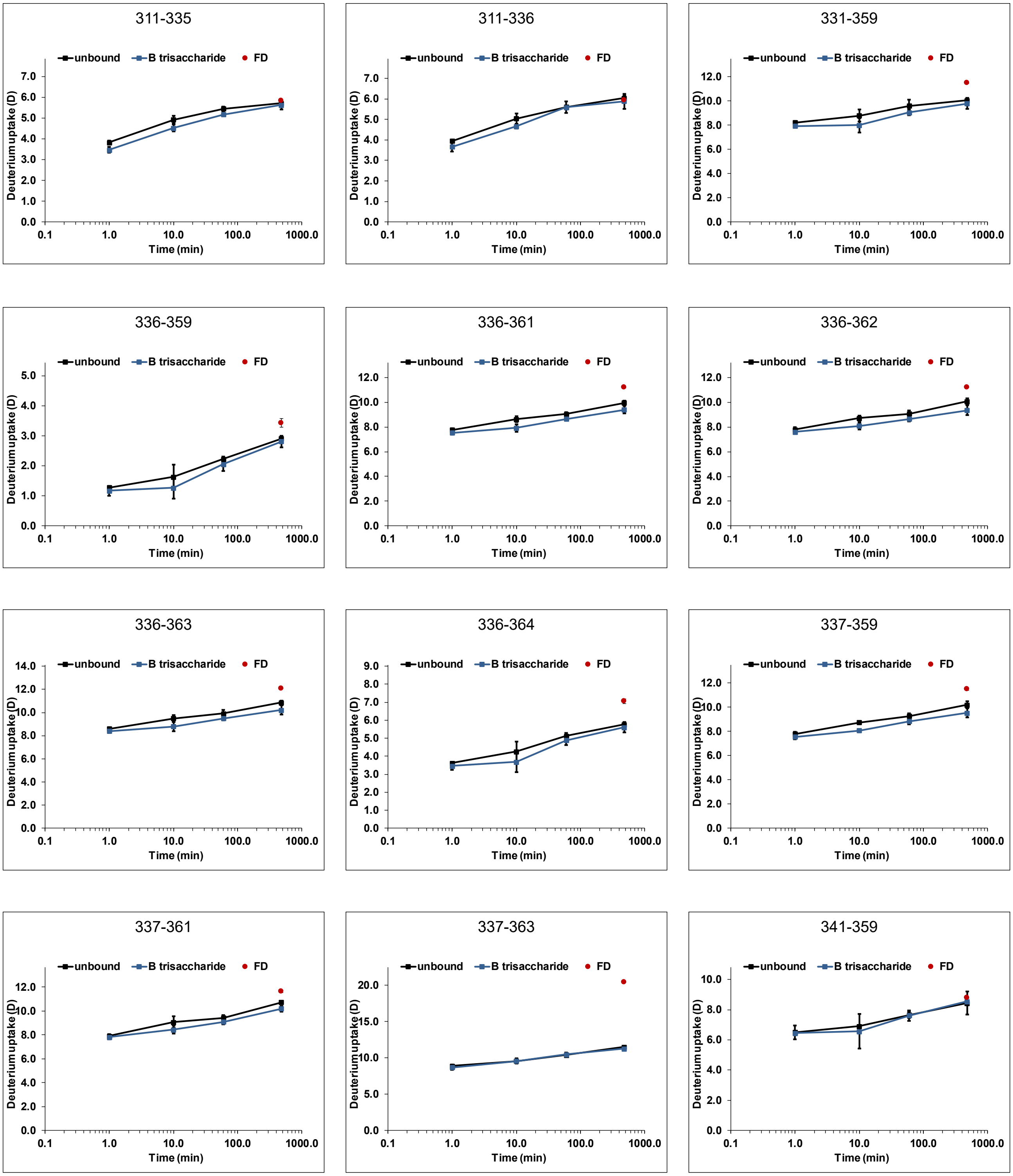

Figure S 25: continued

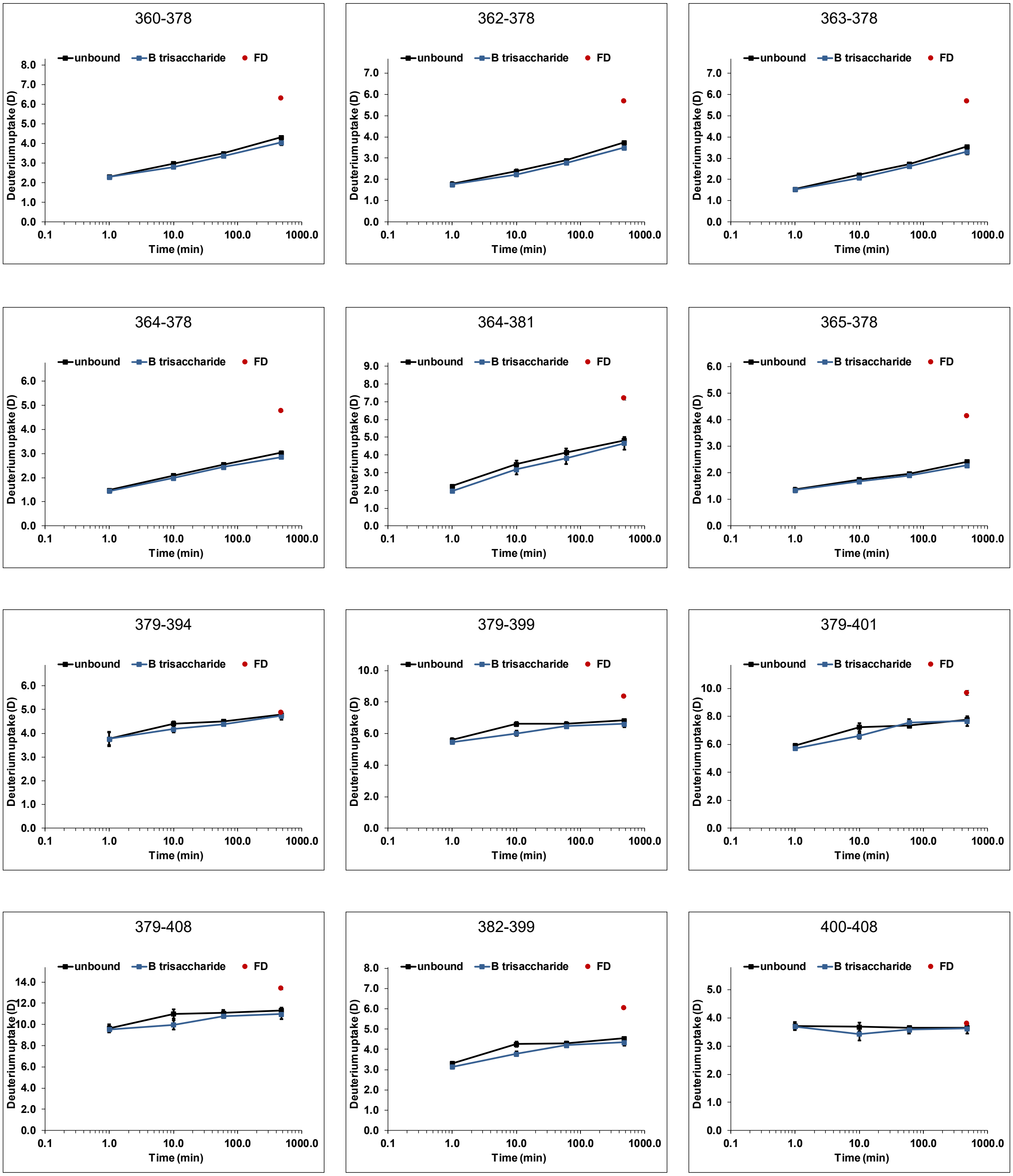


Figure S 25: continued

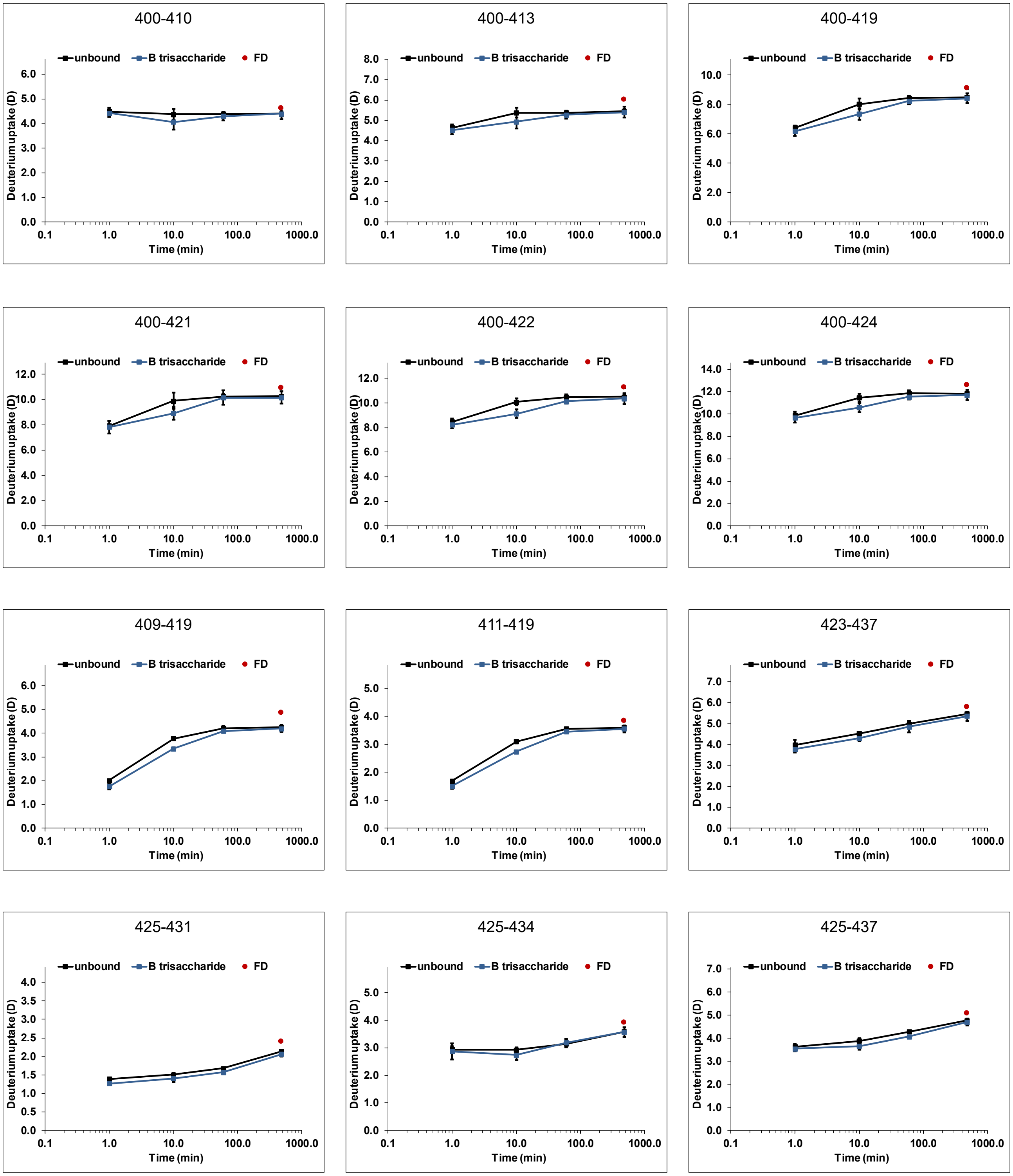


Figure S 25: continued

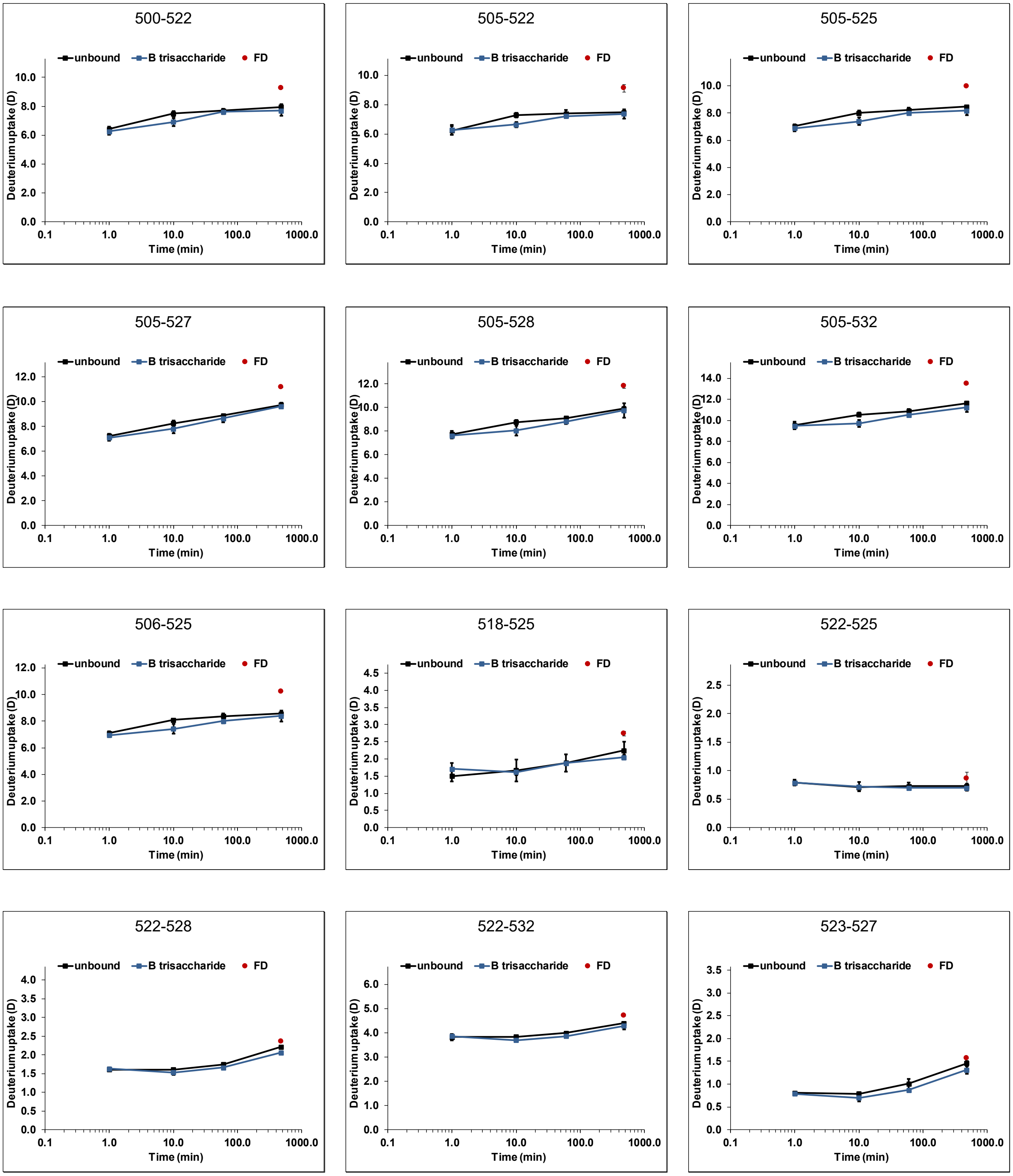


Figure S 25: continued

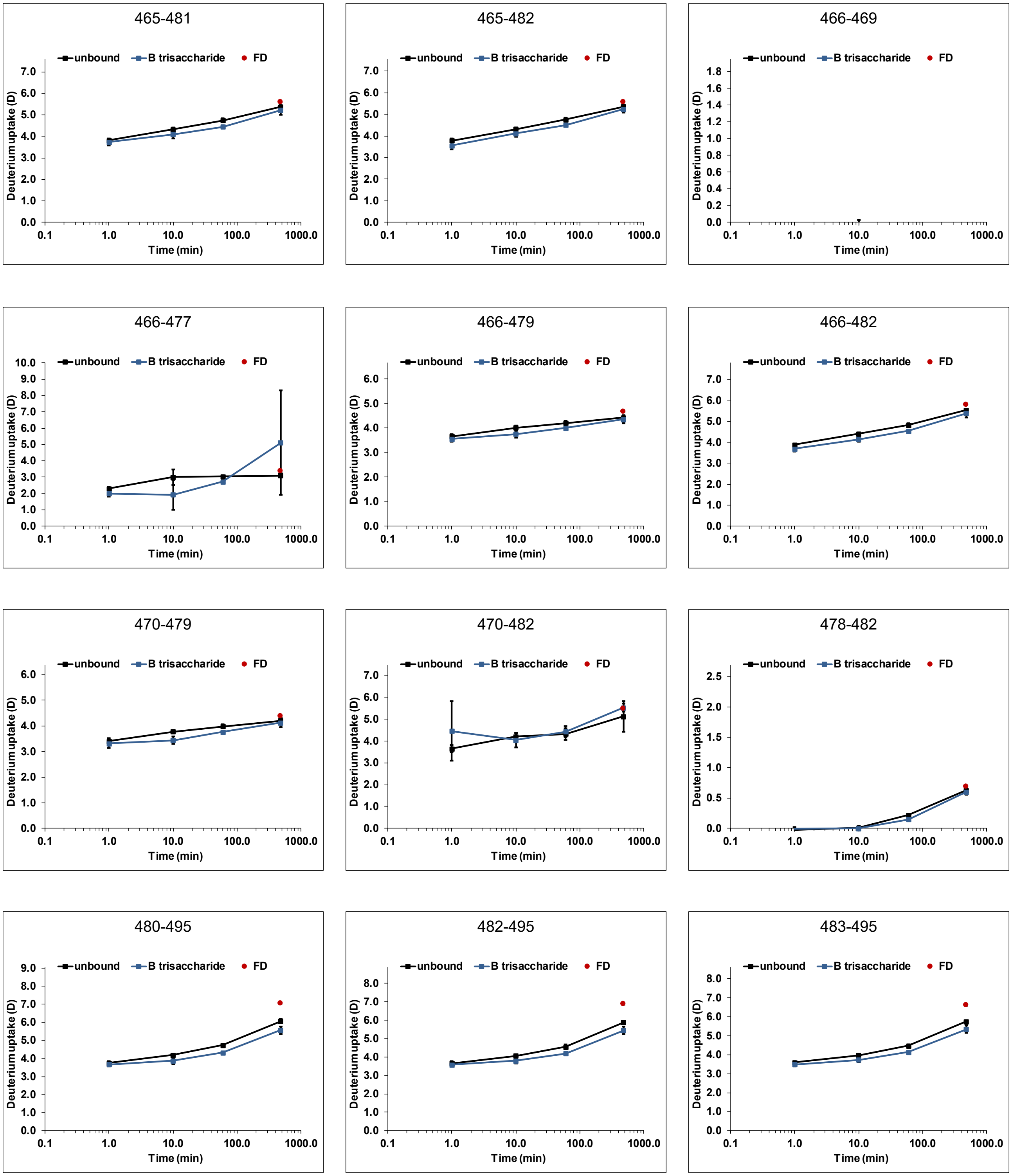

Figure S 25: continued

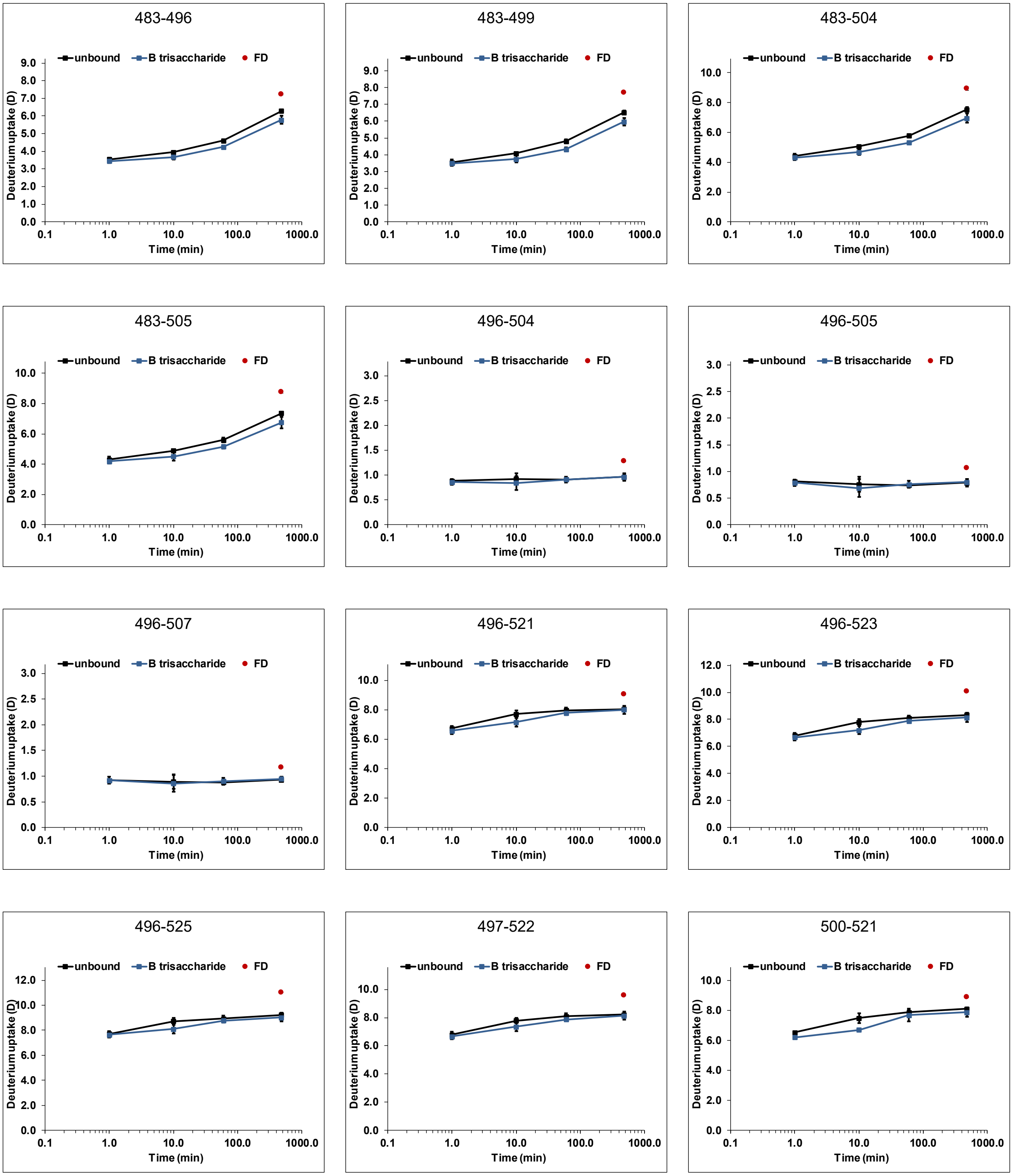


Figure S 25: continued

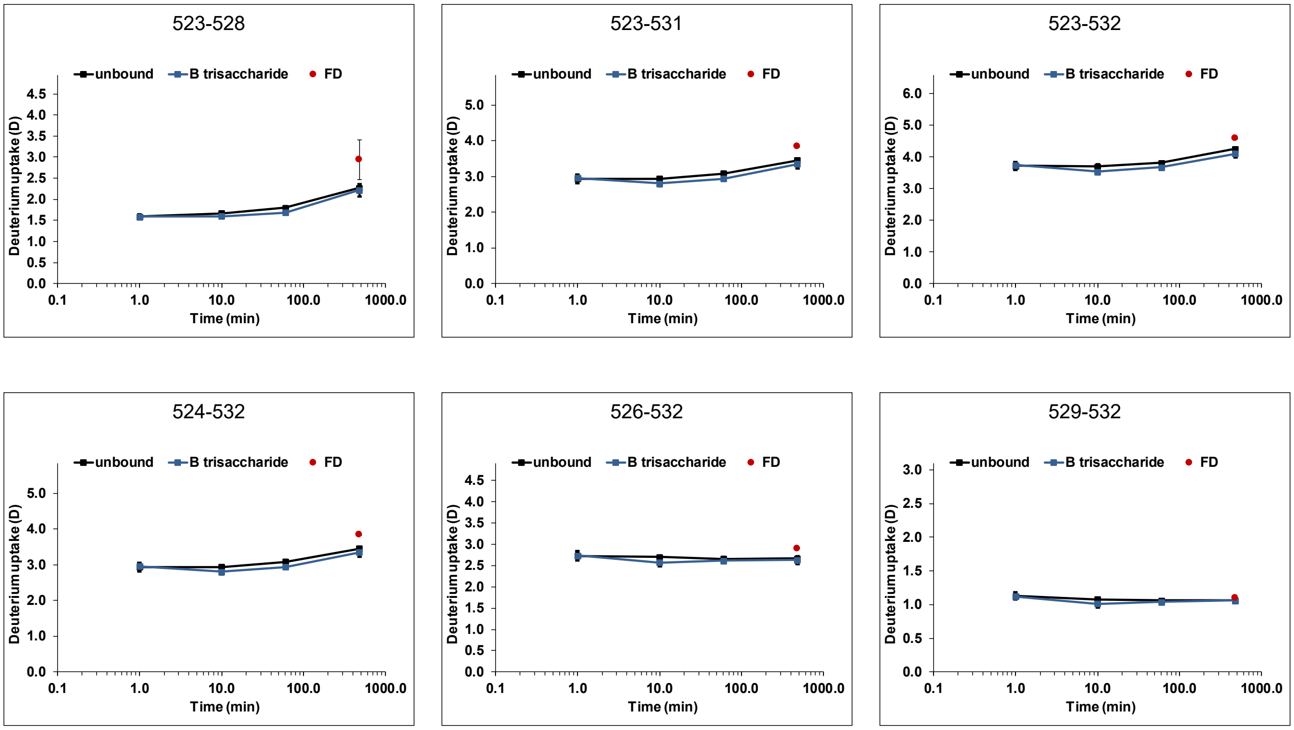


Figure S 25: continued

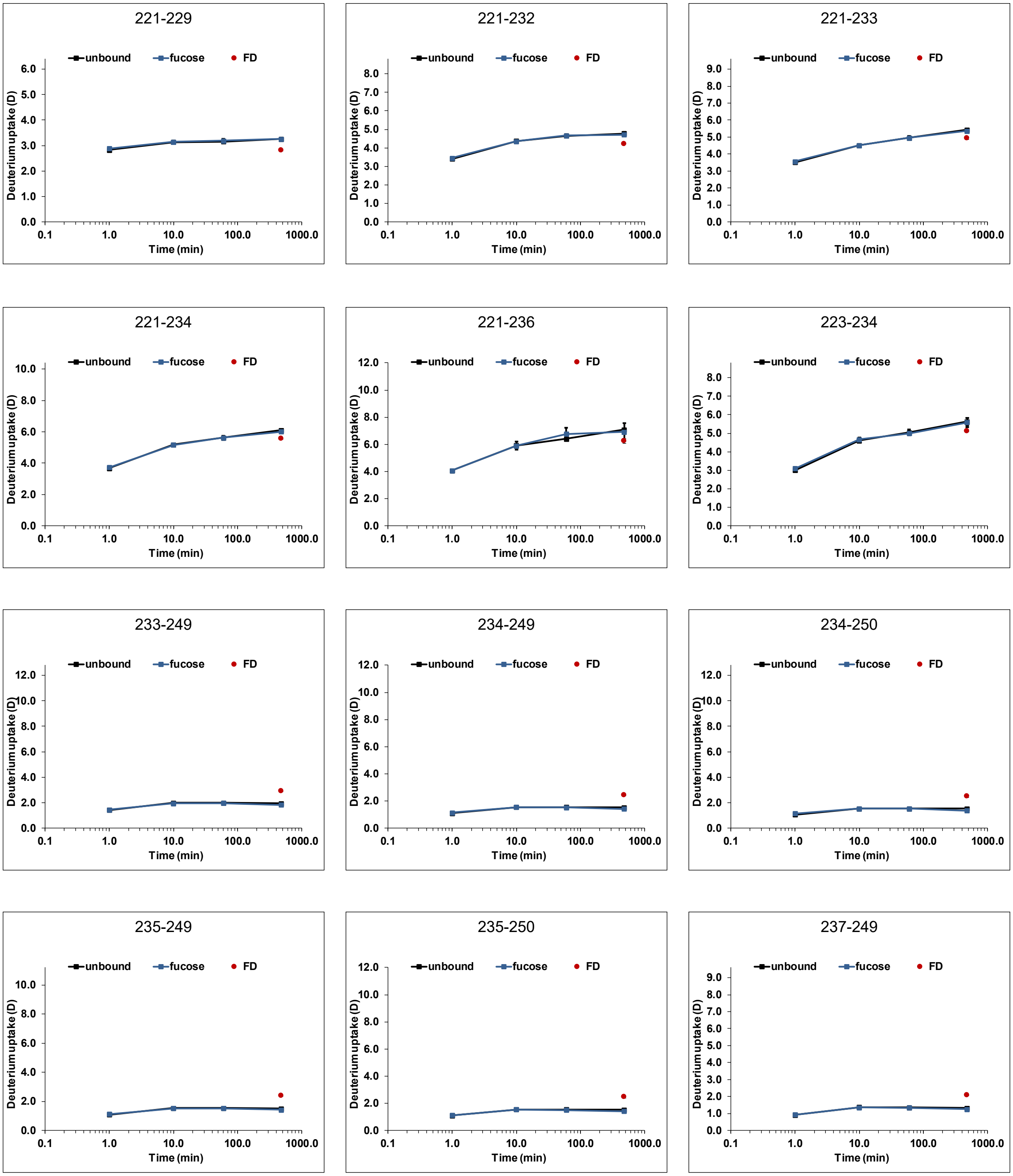


Figure S 26: Deuterium uptake plots for Kawasaki P dimers (black) with 100 mM fucose (blue) (dataset T3). All time points (1 min, 10 min, 1 h and 8 h) were performed in triplicate, the fully deuterated control (FD, red) represents two measurements. Error bars indicate the standard deviation (SD).

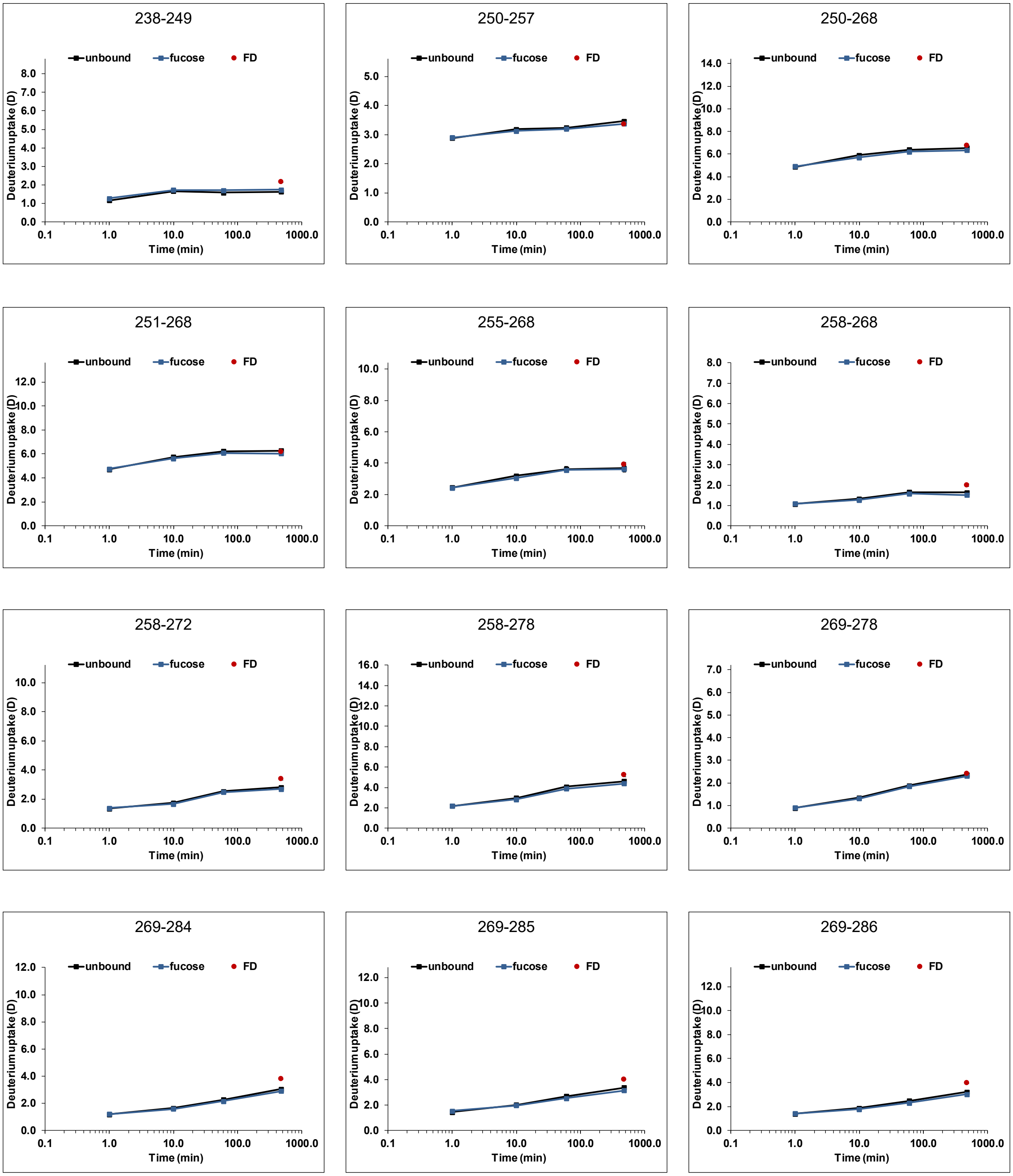


Figure S 26: continued

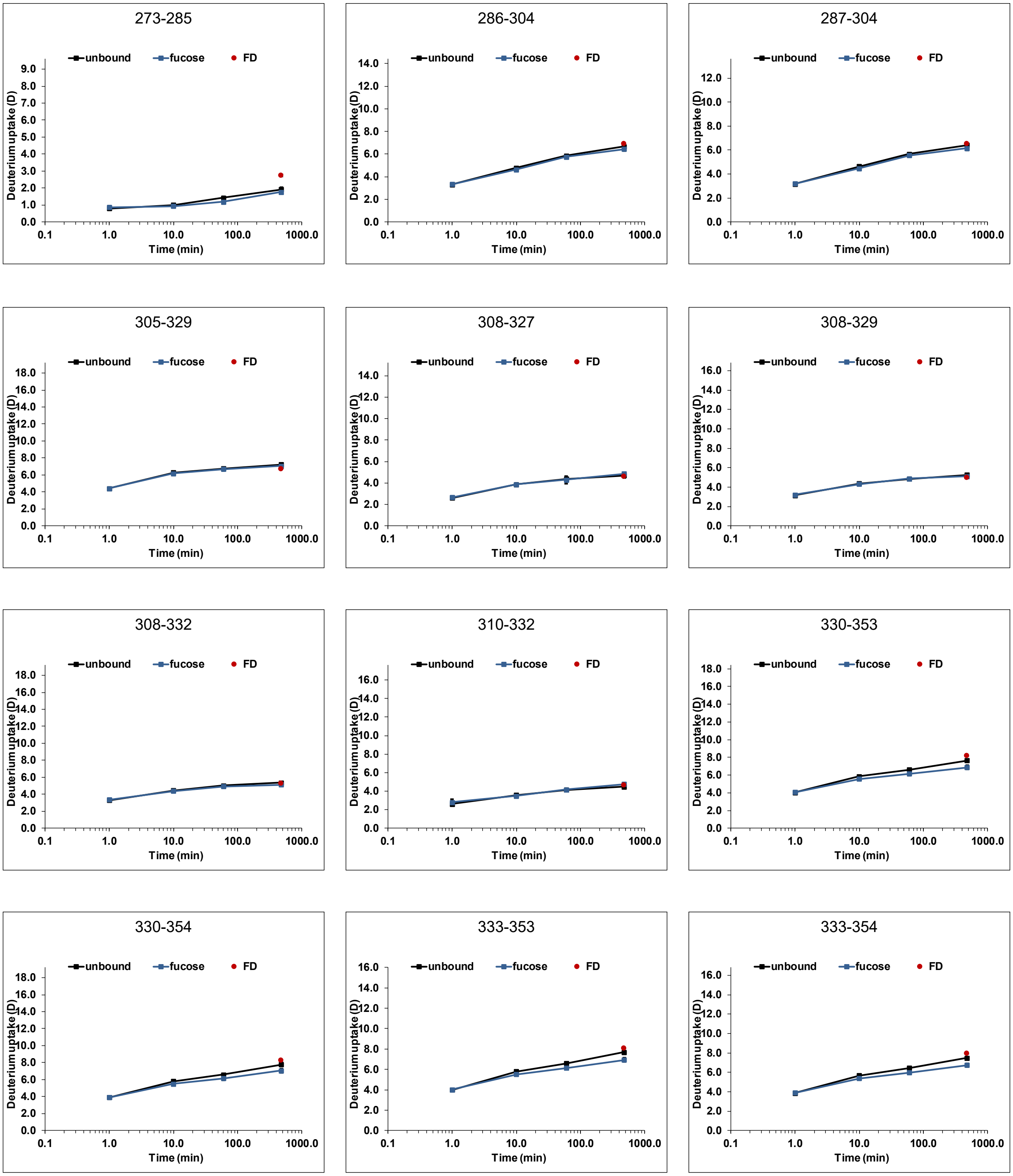


Figure S 26: continued

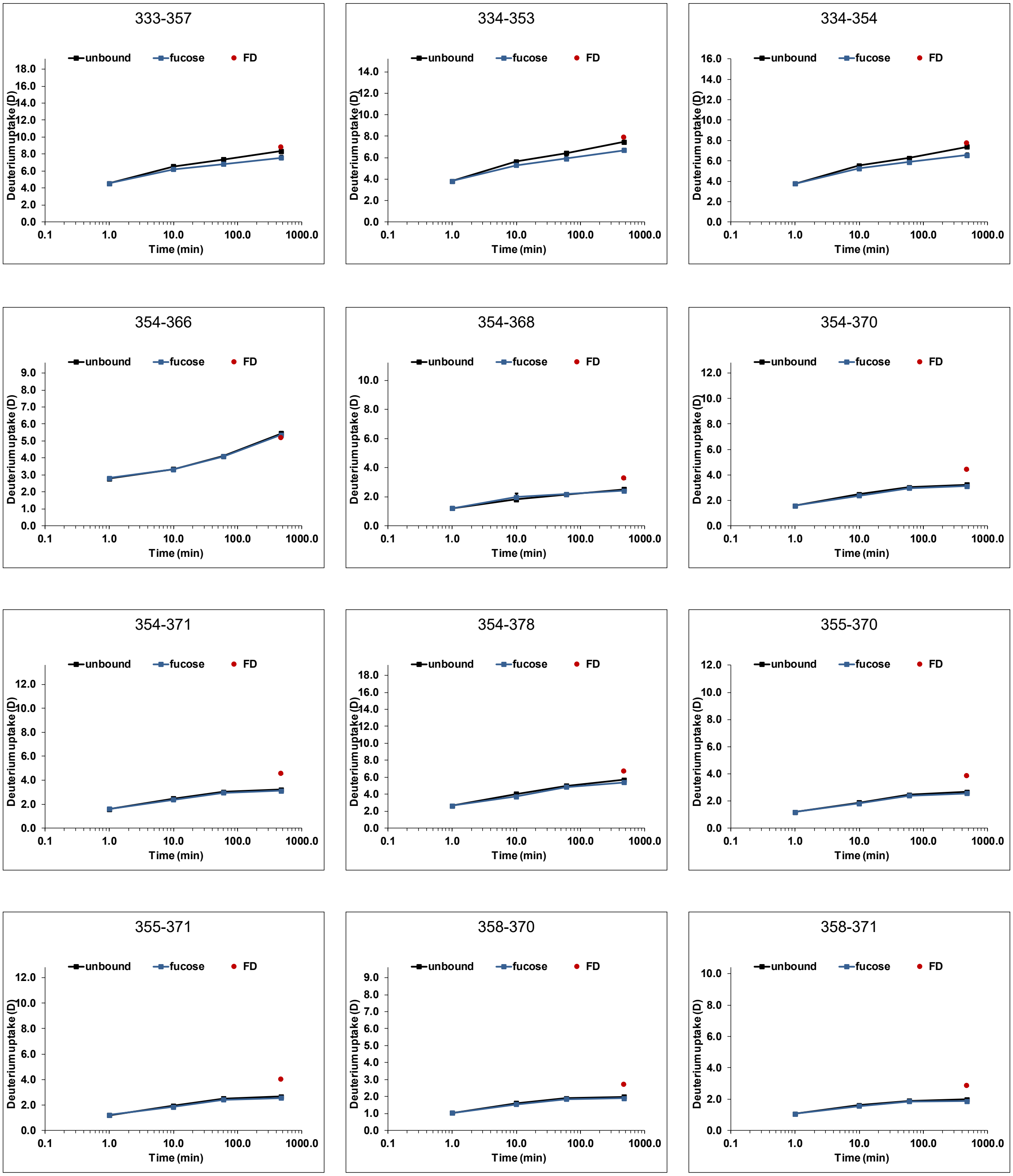


Figure S 26: continued

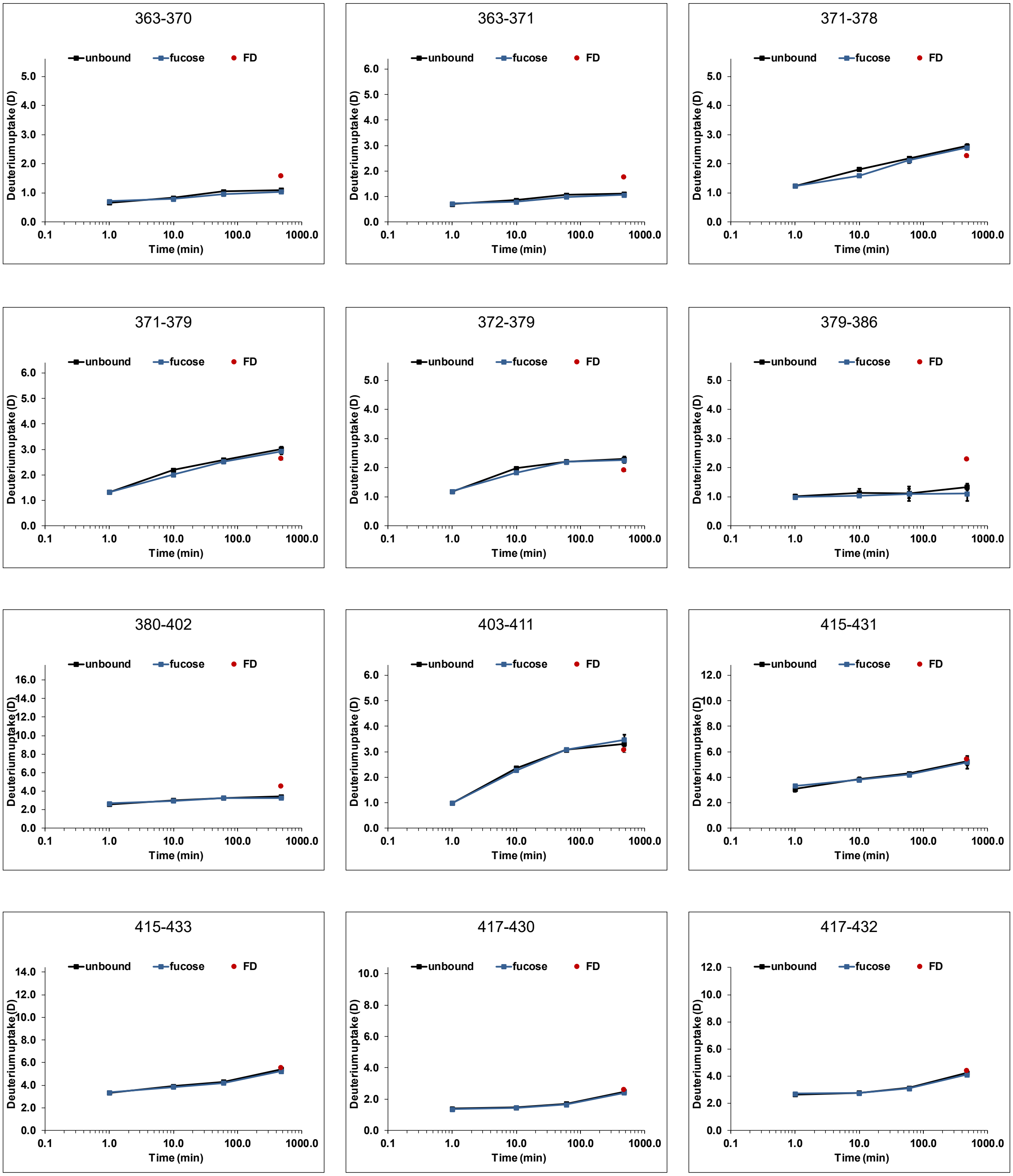


Figure S 26: continued

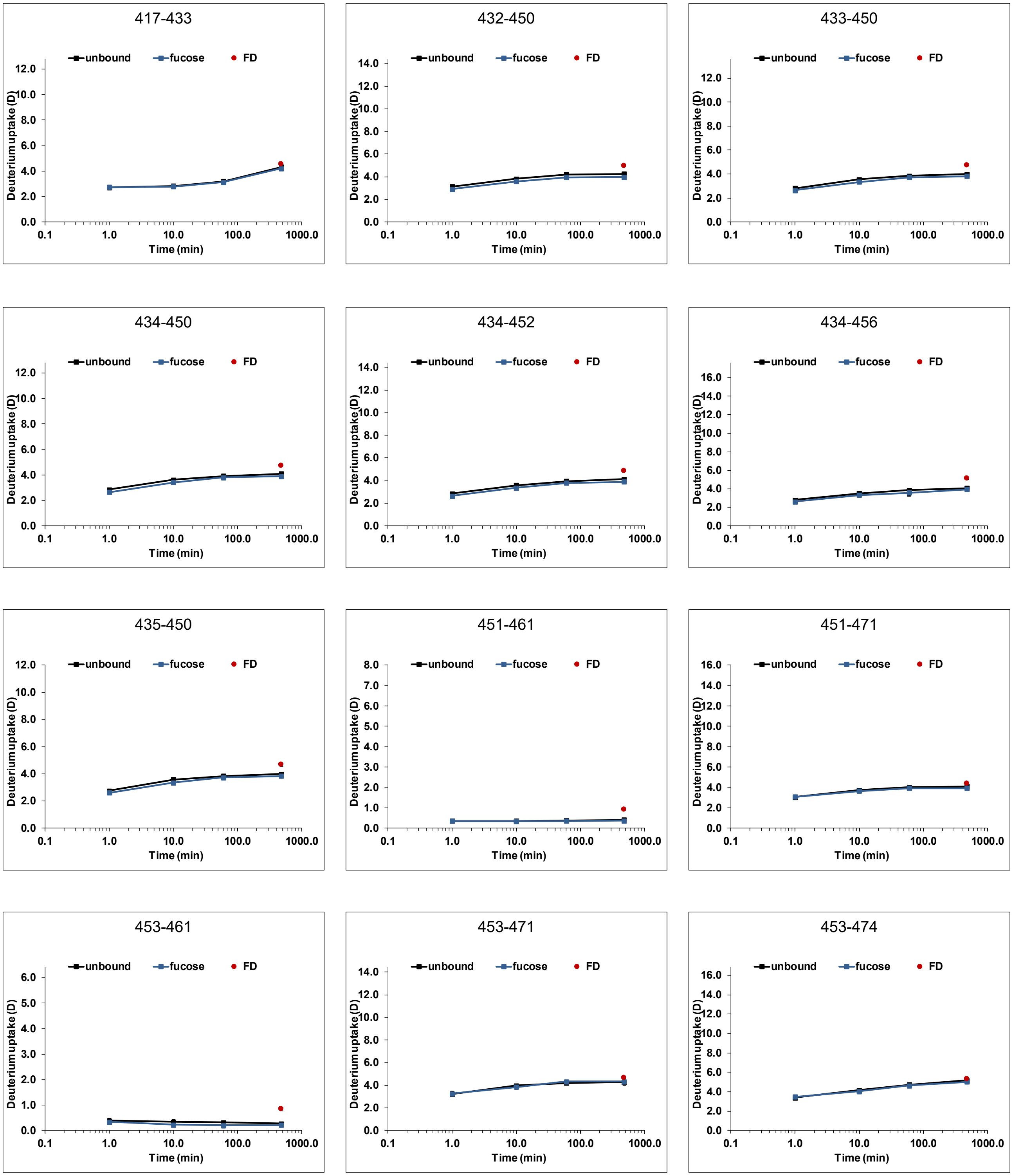


Figure S 26: continued

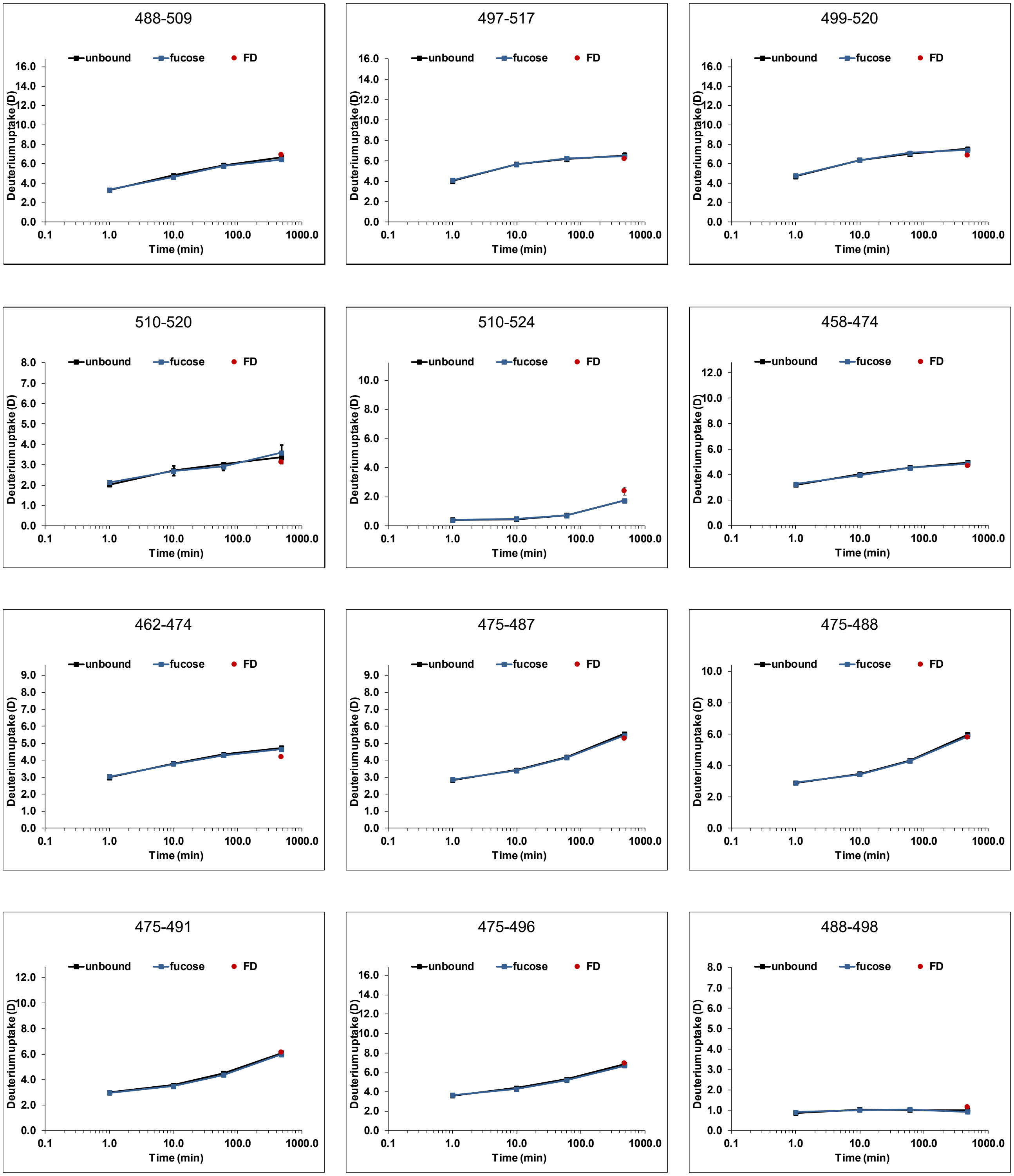

Figure S 26: continued

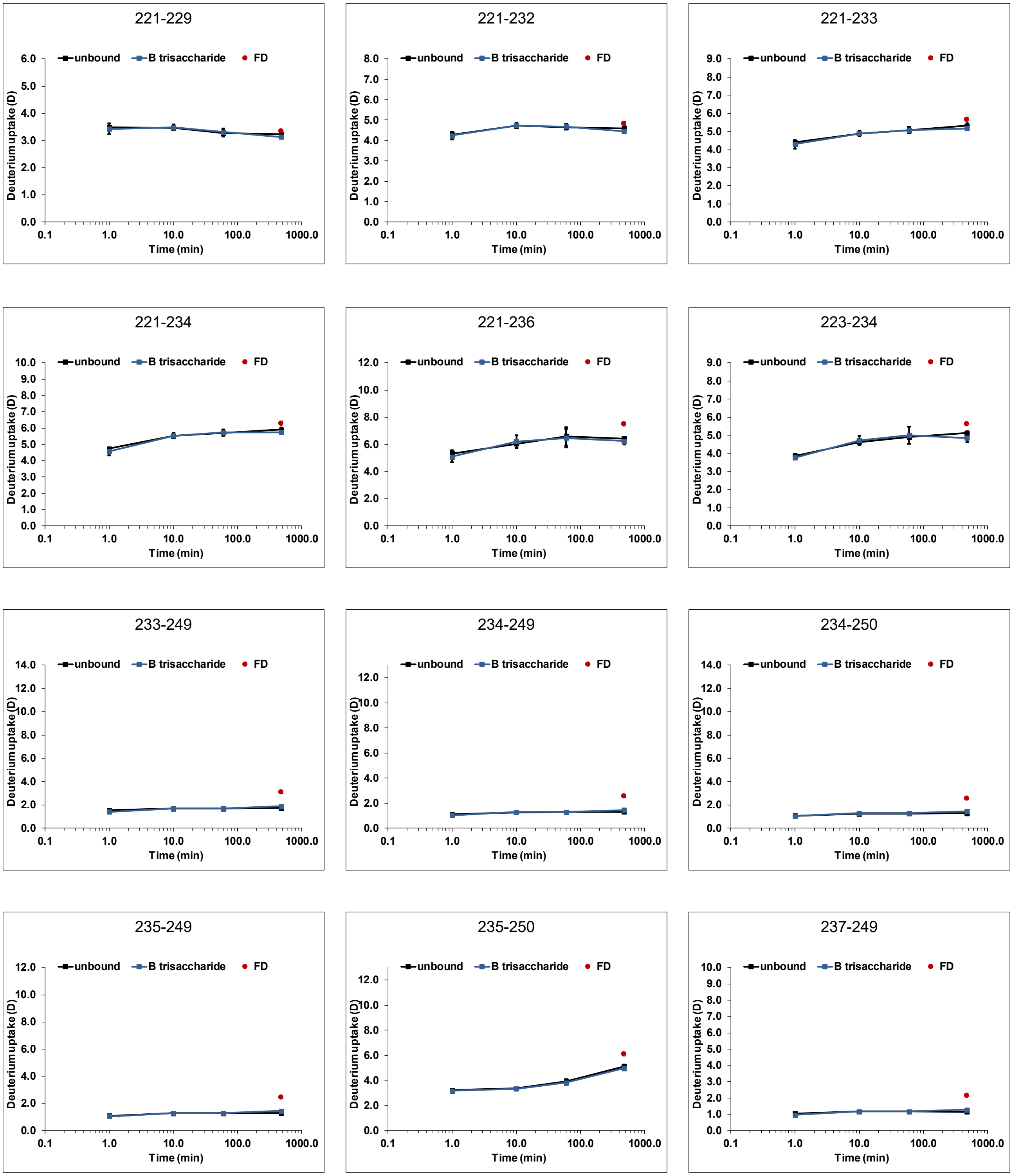


Figure S 27: Deuterium uptake plots for Kawasaki P dimers (black) with 10 mM HBGA B trisaccharide (blue) (dataset T4). All time points (1 min, 10 min, 1 h and 8 h) were performed in triplicate, the fully deuterated control (FD, red) represents two measurements. Error bars indicate the standard deviation (SD).

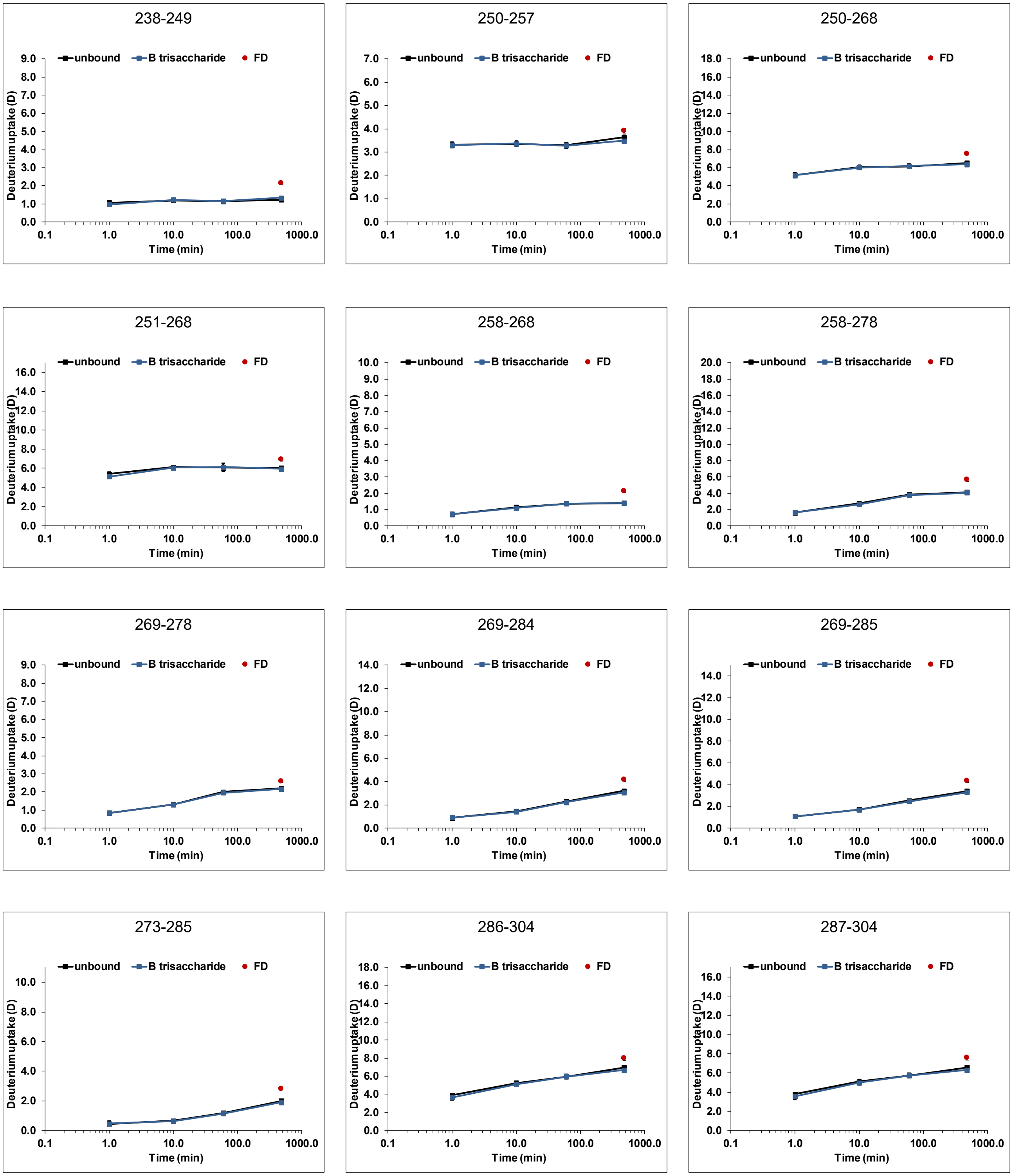


Figure S 27: continued

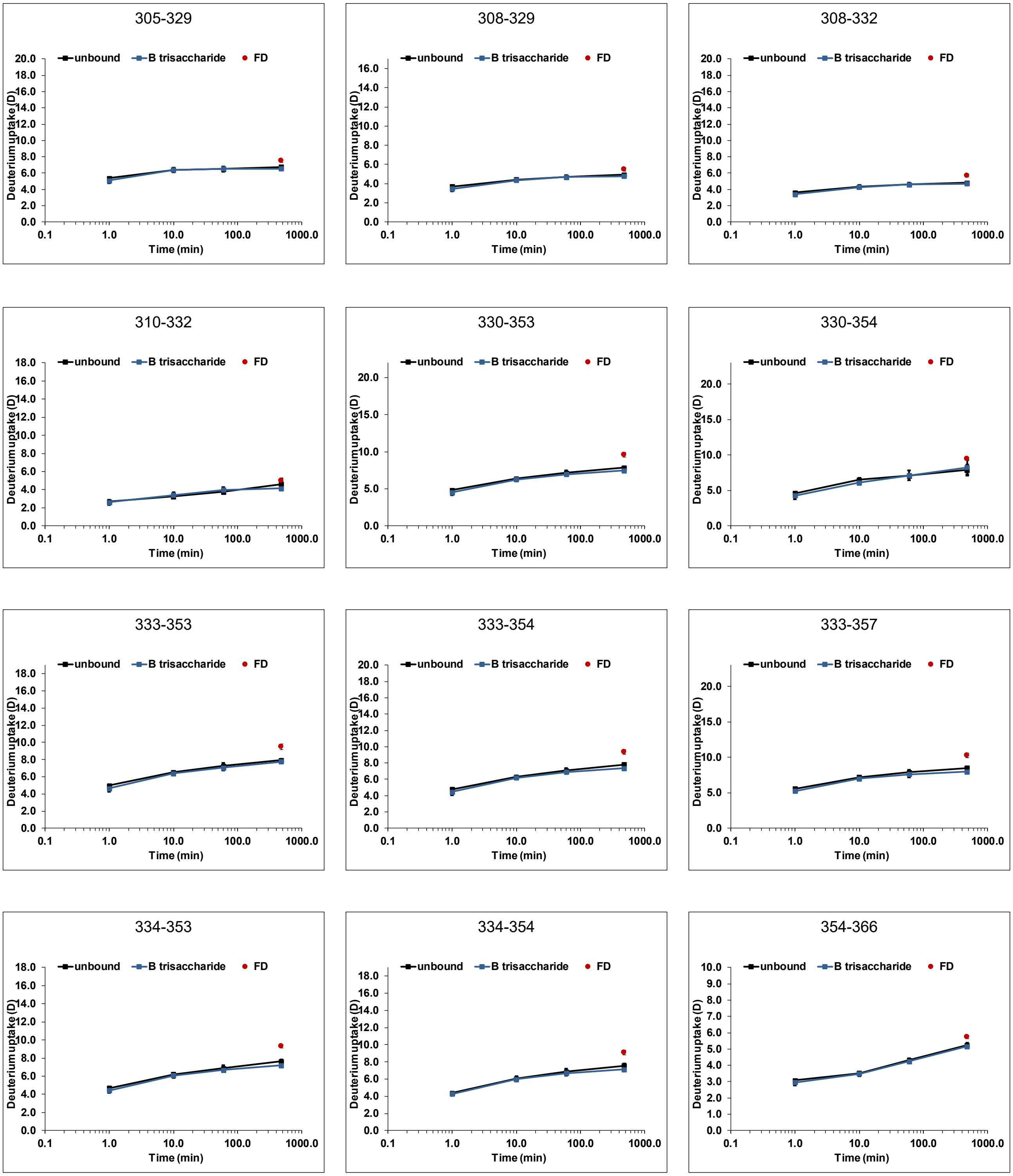


Figure S 27: continued

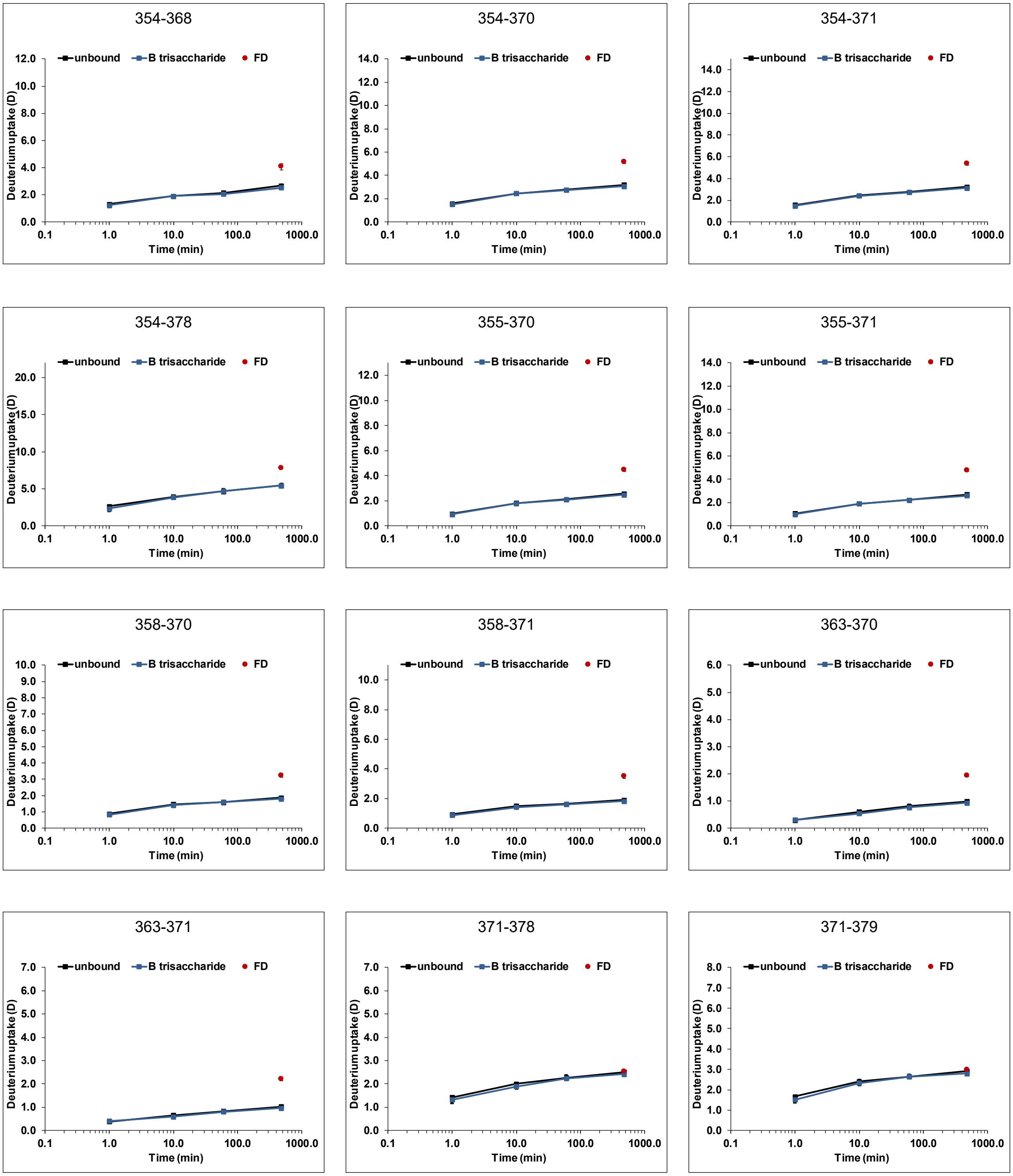


Figure S 27: continued

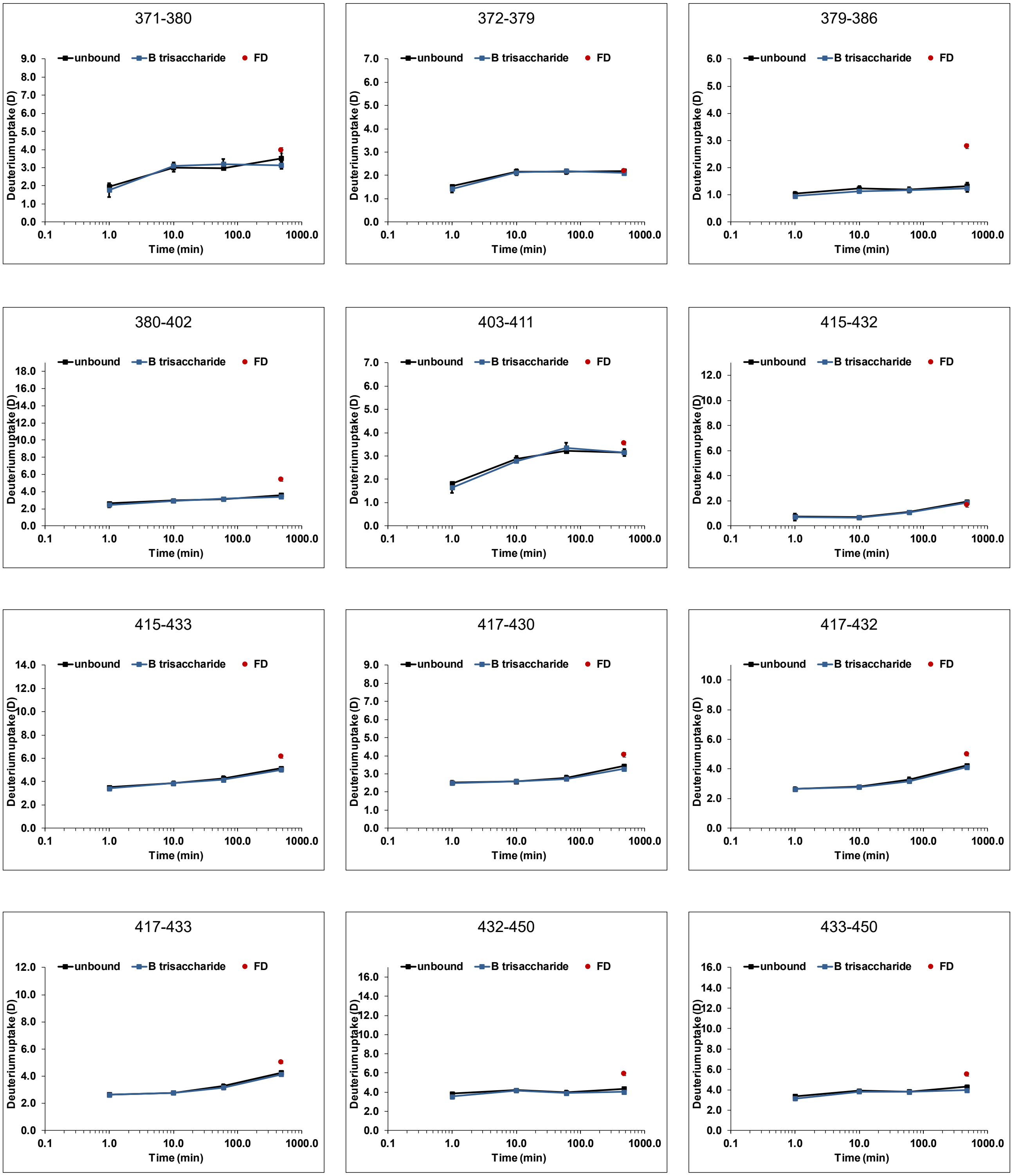


Figure S 27: continued

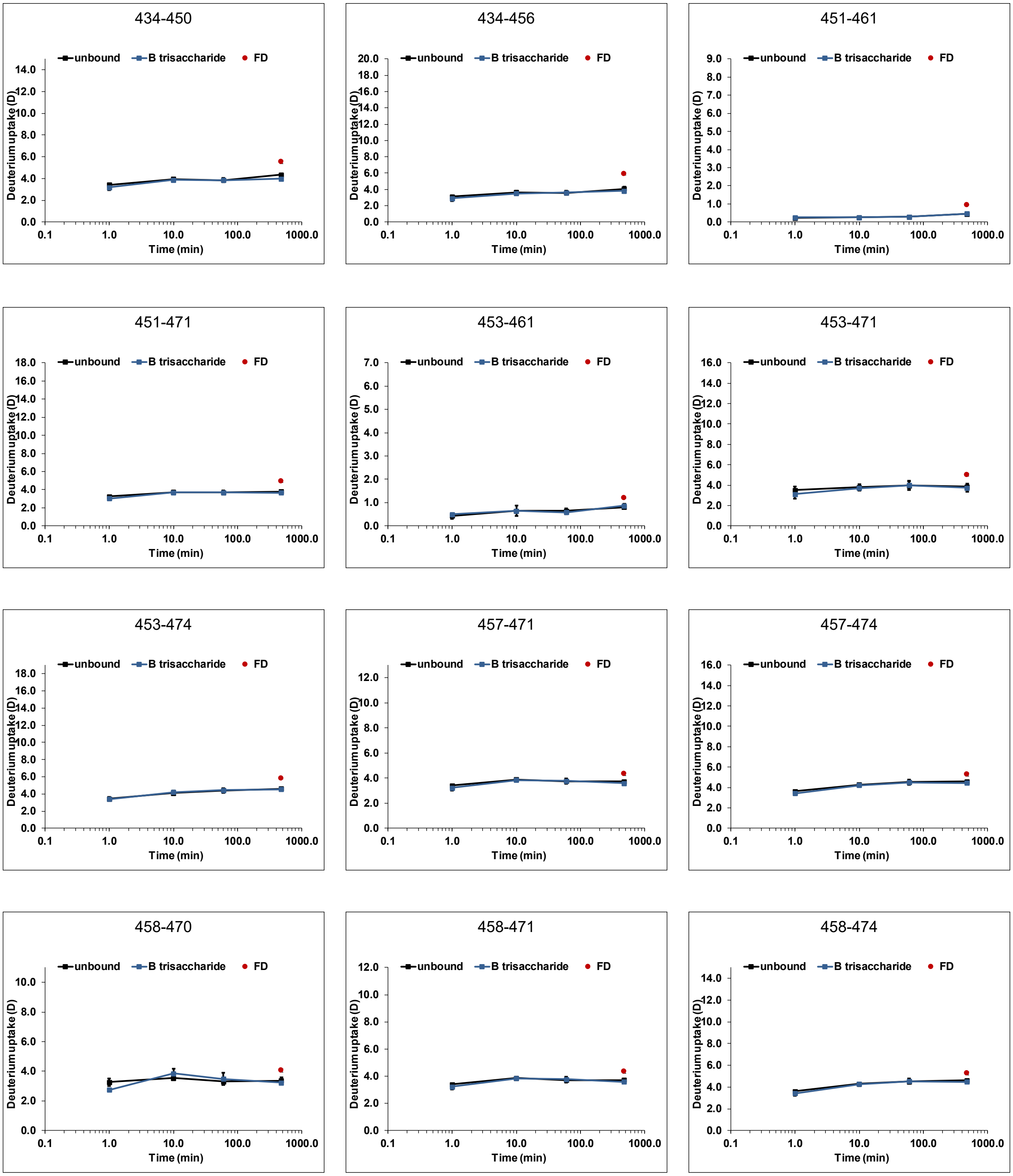


Figure S 27: continued

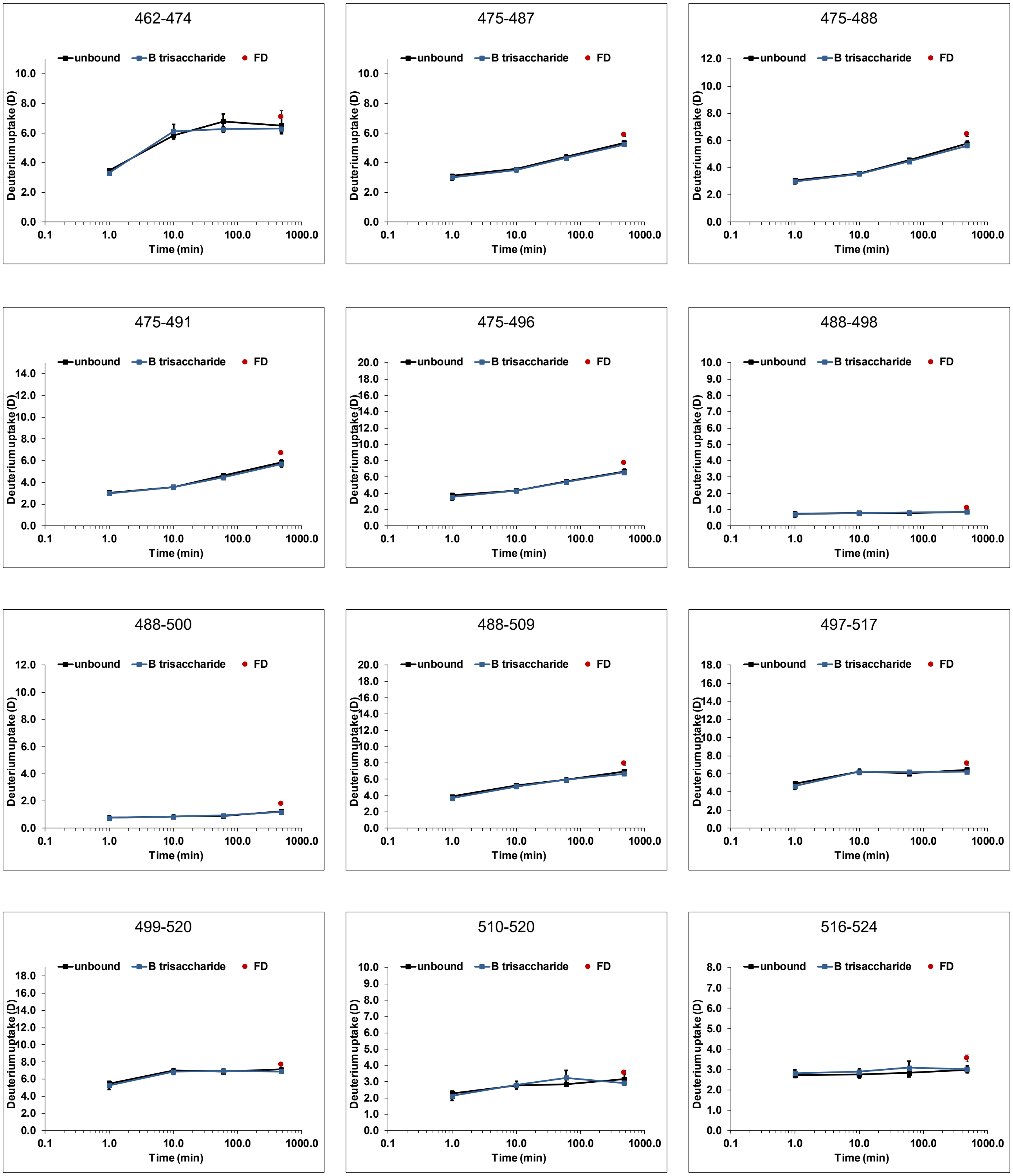


Figure S 27: continued

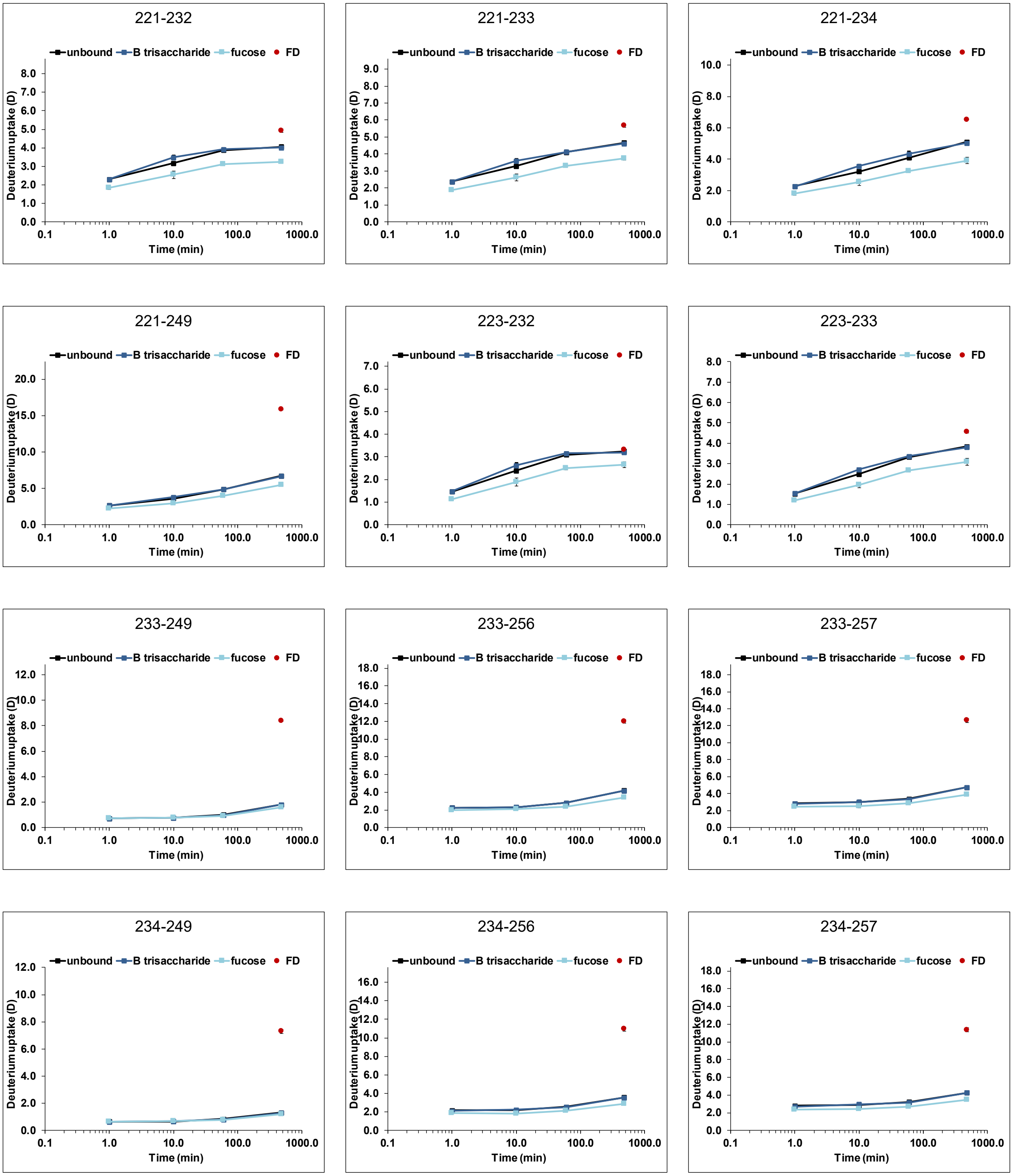


Figure S 28: Deuterium uptake plots for wildtype MI001 P dimers (black) with 10 mM HBGA B trisaccharide (blue) and 100 mM fucose (light blue) (dataset T5). All time points (1 min, 10 min, 1 h and 8 h) were performed in triplicate, the fully deuterated control (FD, red) represents two measurements. Error bars indicate the standard deviation (SD).

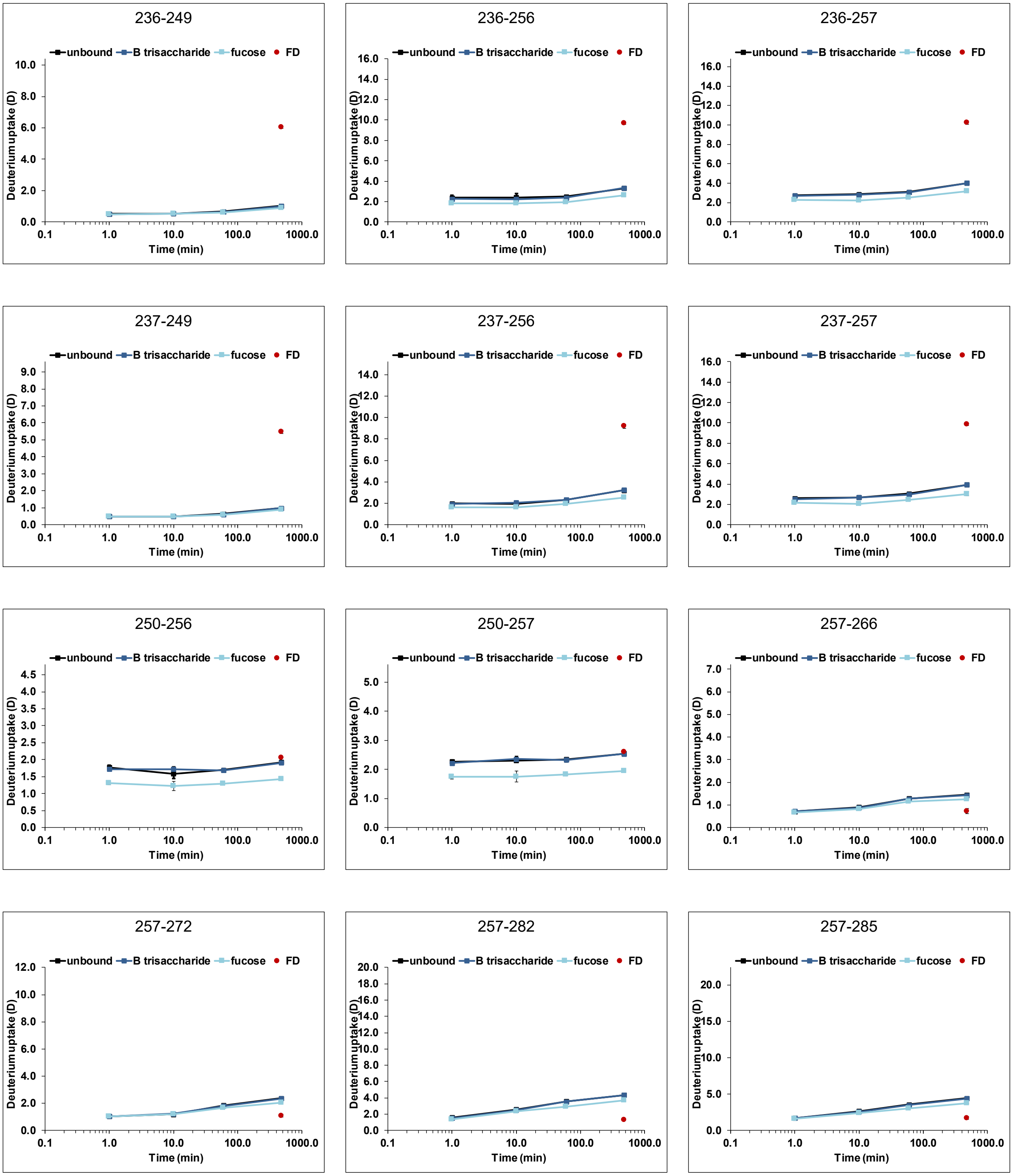


Figure S 28: continued

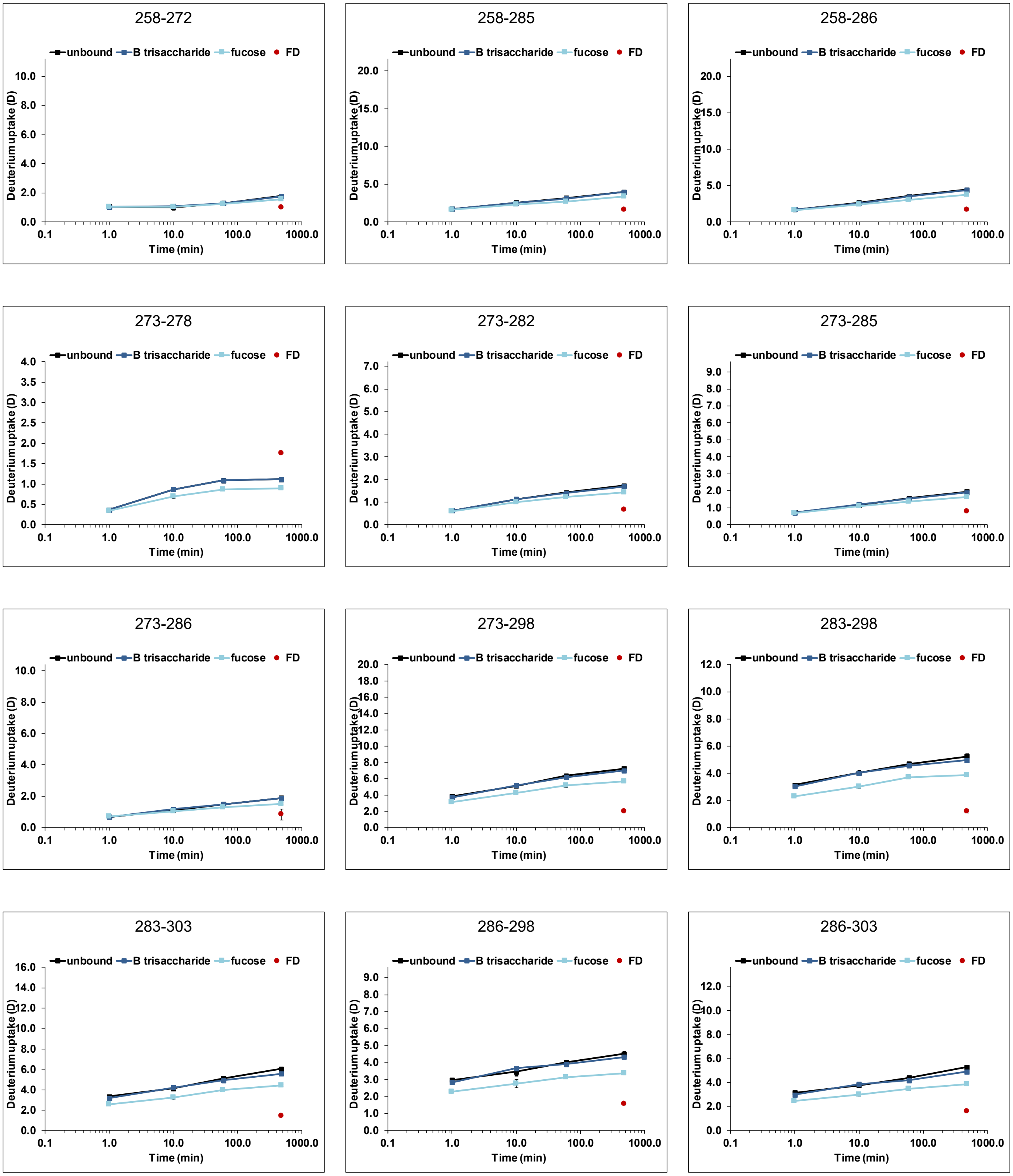


Figure S 28: continued

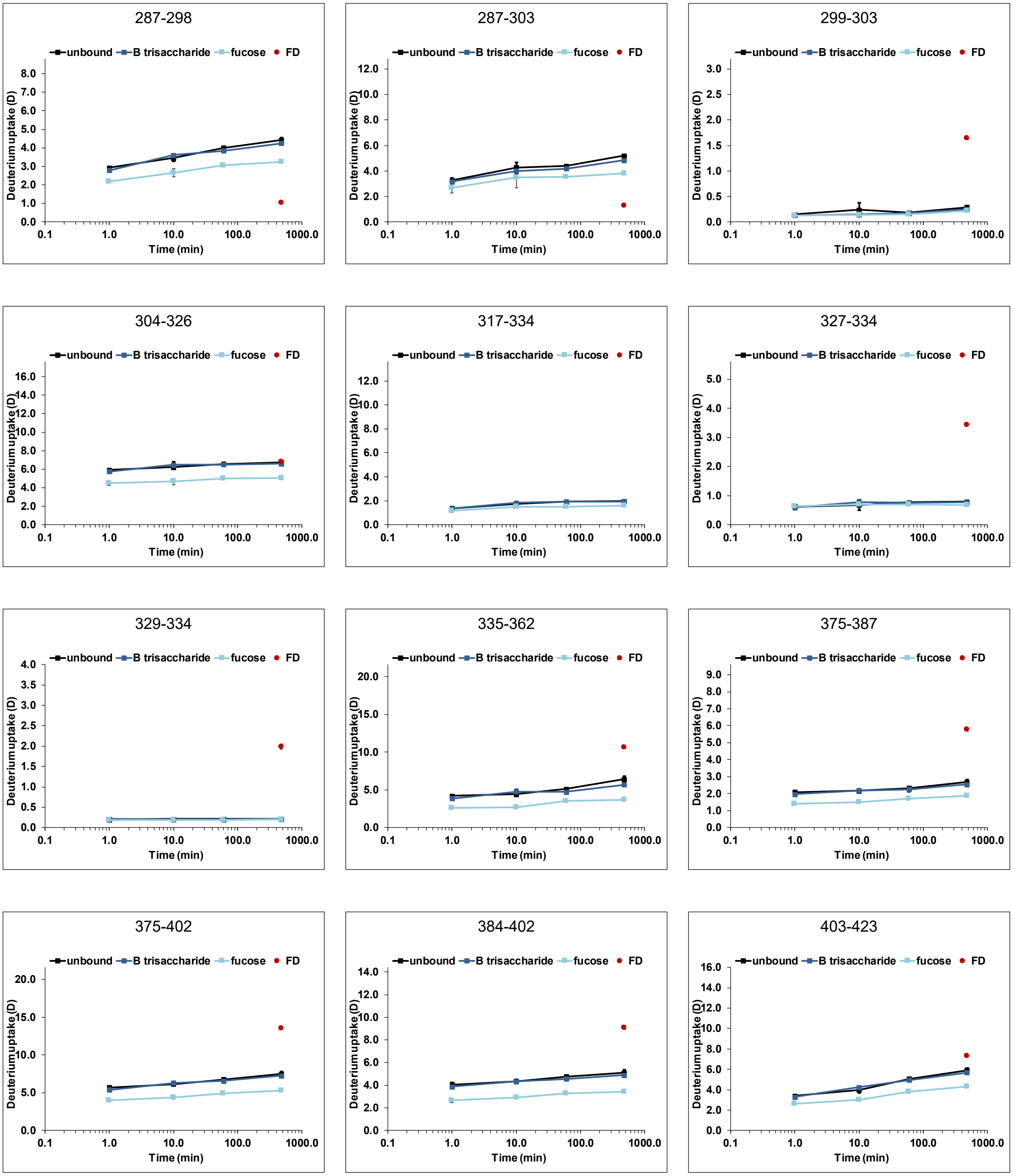


Figure S 28: continued

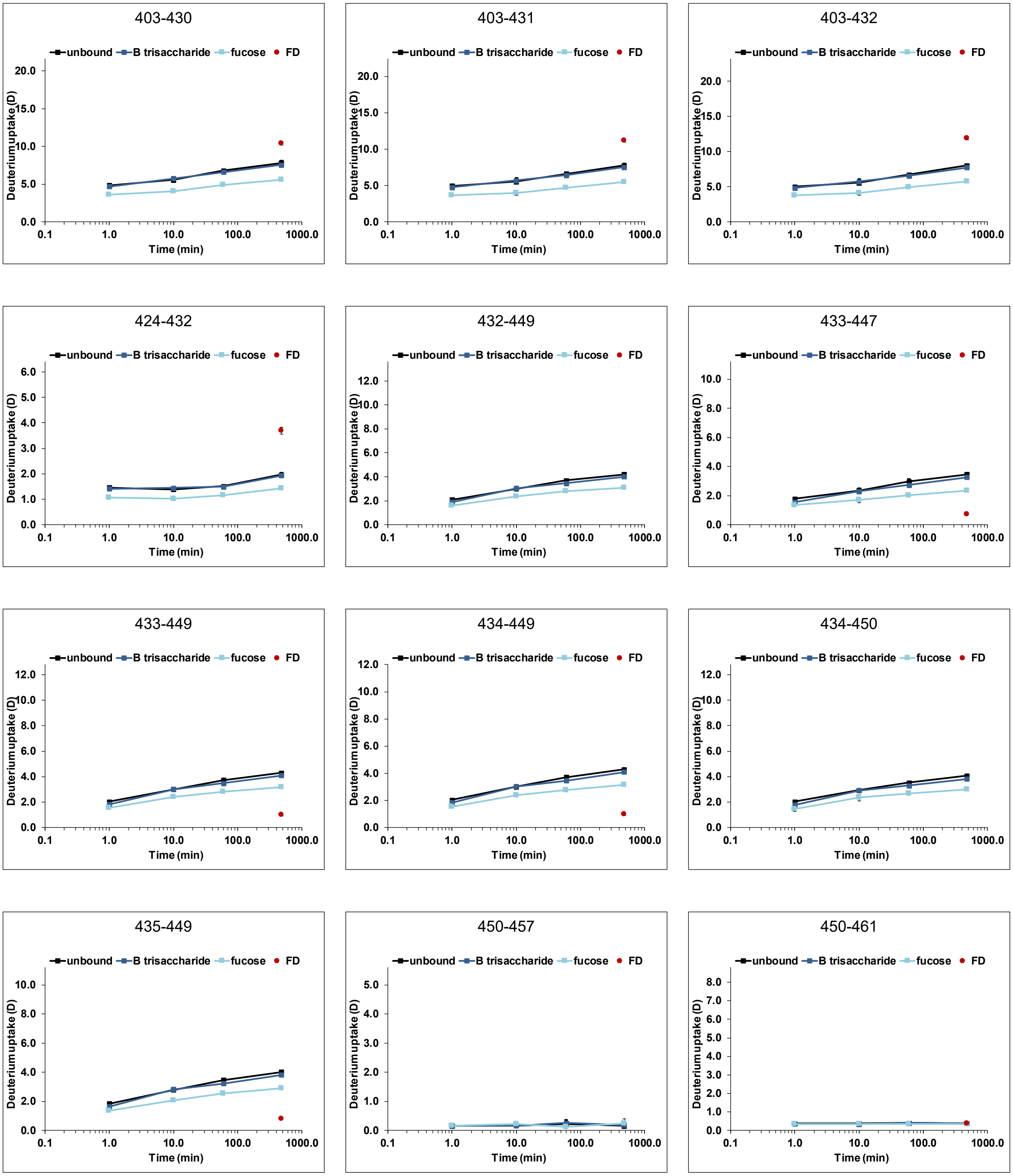


Figure S 28: continued

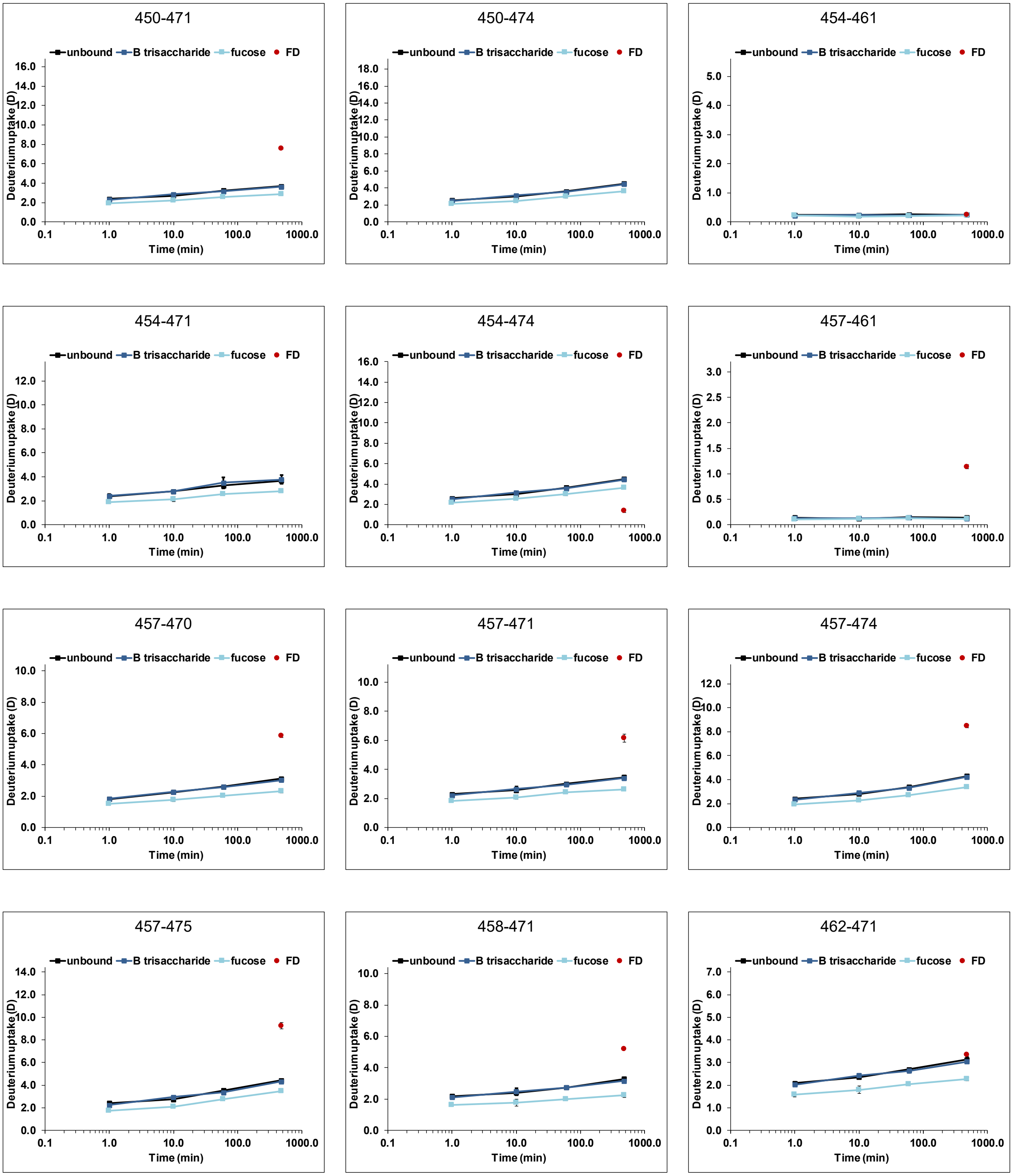


Figure S 28: continued

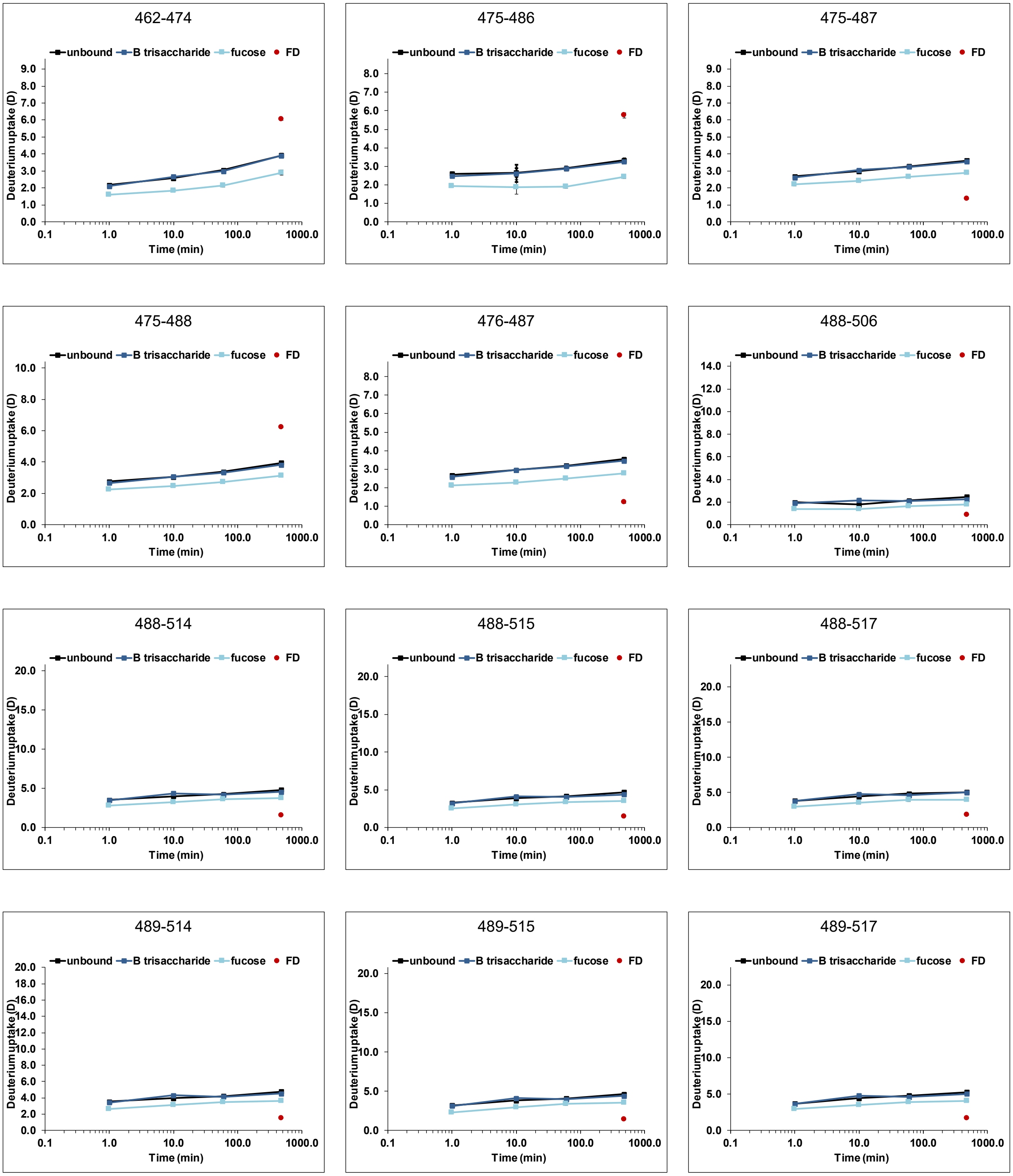


Figure S 28: continued

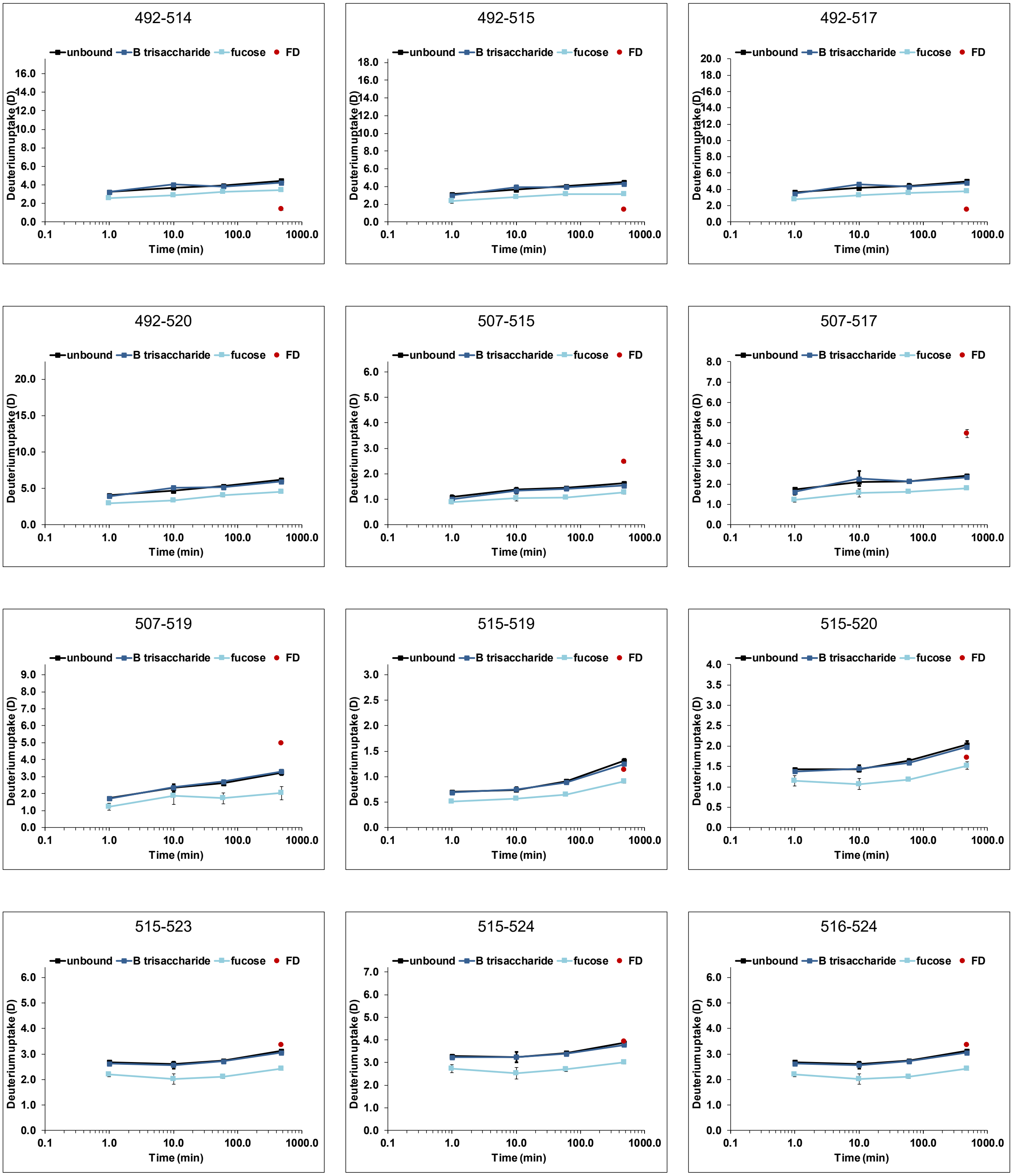


Figure S 28: continued

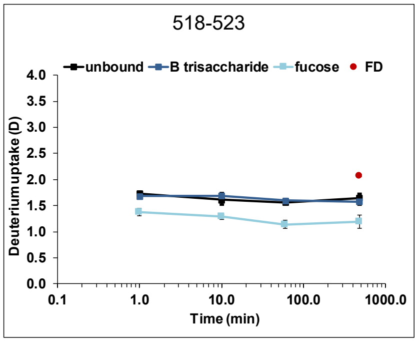


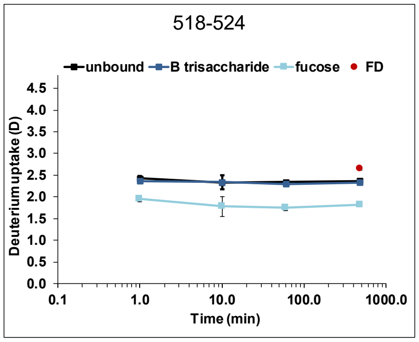


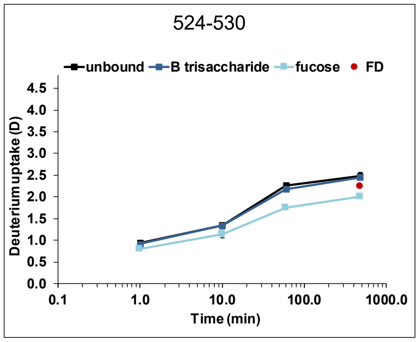


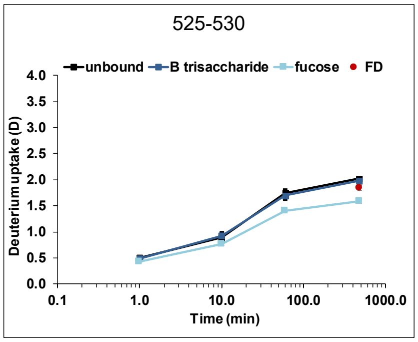


Figure S 28: continued

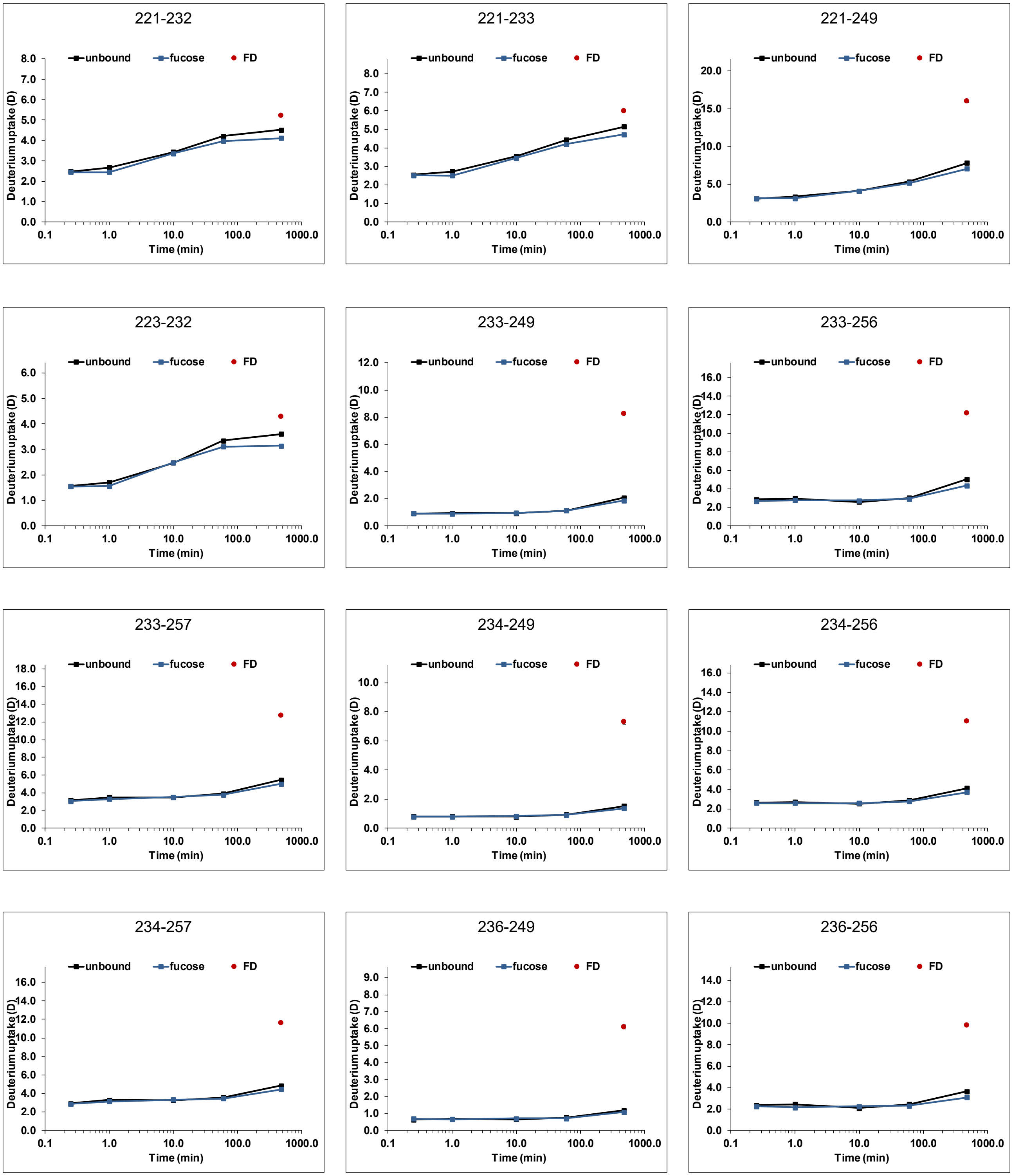


Figure S 29: Deuterium uptake plots for wildtype MI001 P dimers (black) with 100 mM fucose (blue) (dataset T6). All time points (1 min, 10 min, 1 h and 8 h) were performed in single measurements, the fully deuterated control (FD, red) represents two measurements. Error bars indicate the standard deviation (SD).

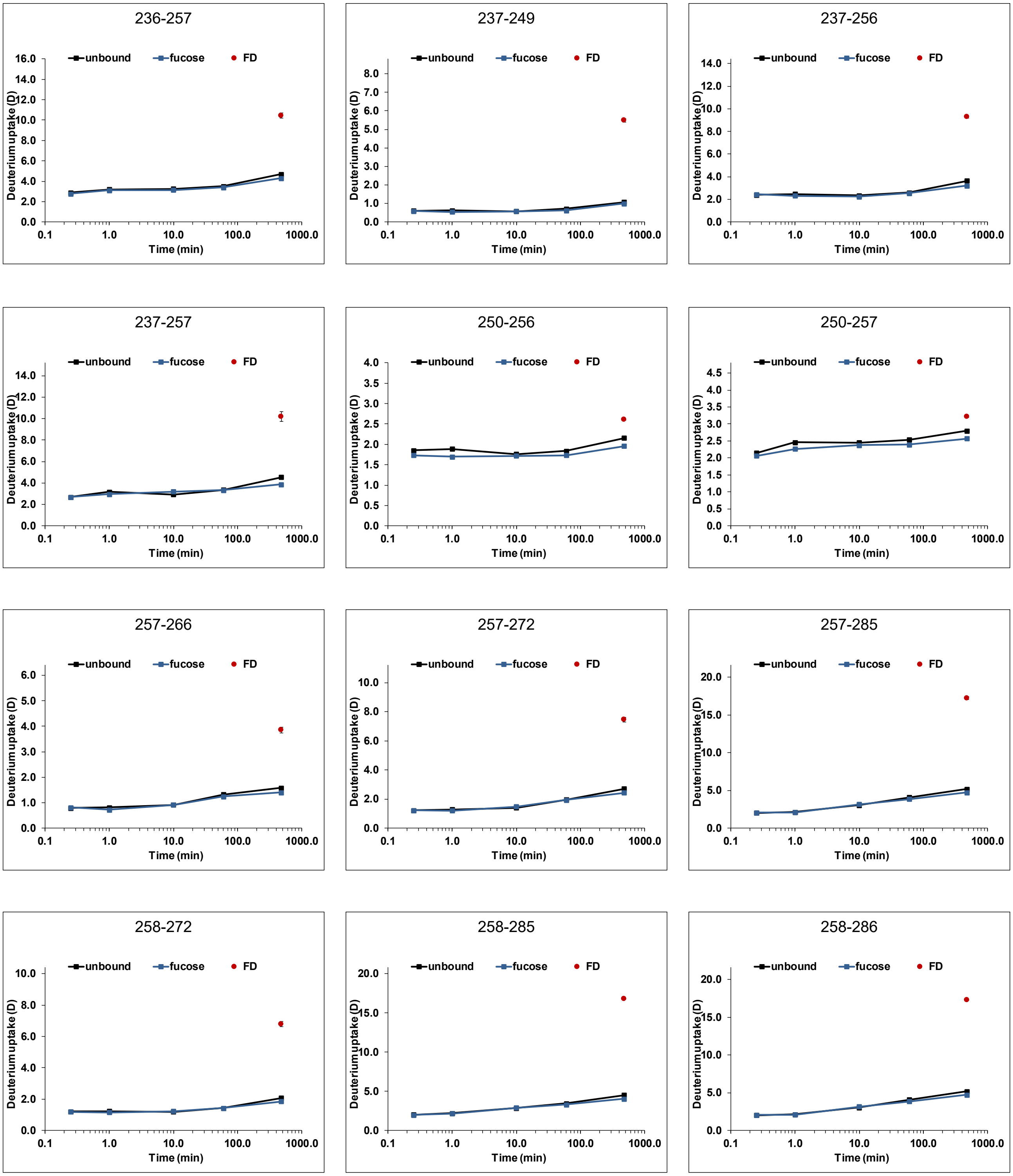


Figure S 29: continued

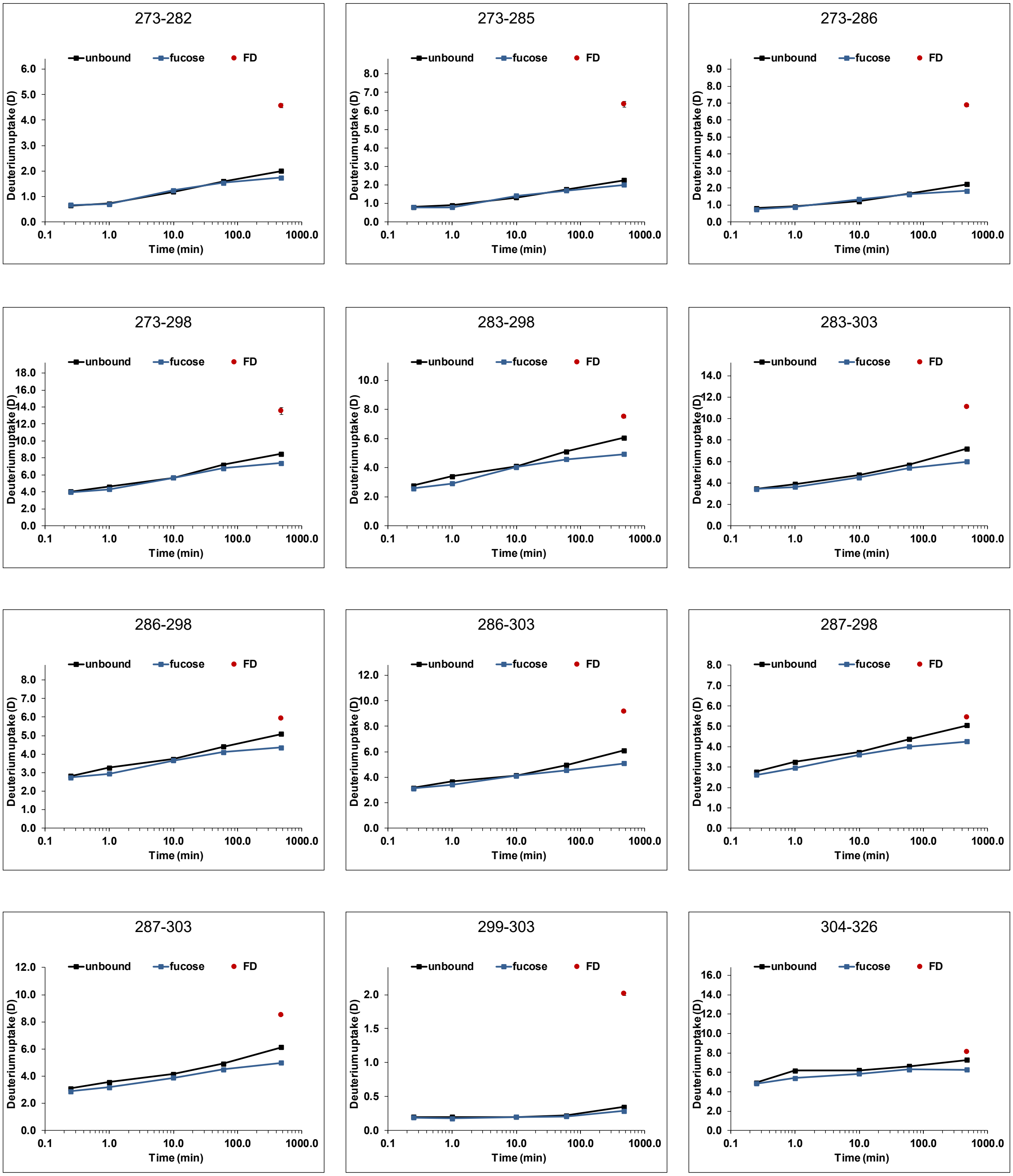


Figure S 29: continued

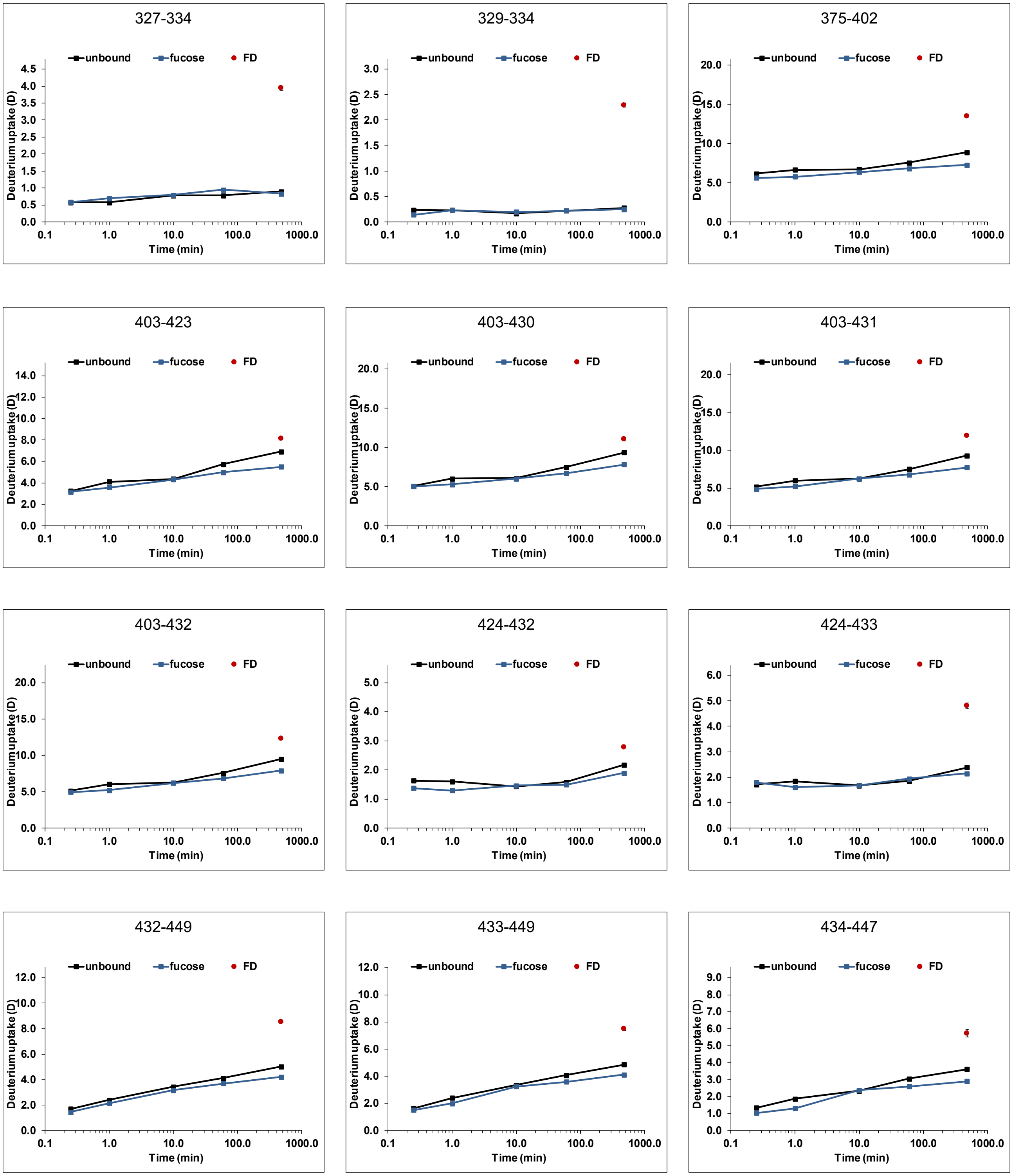


Figure S 29: continued

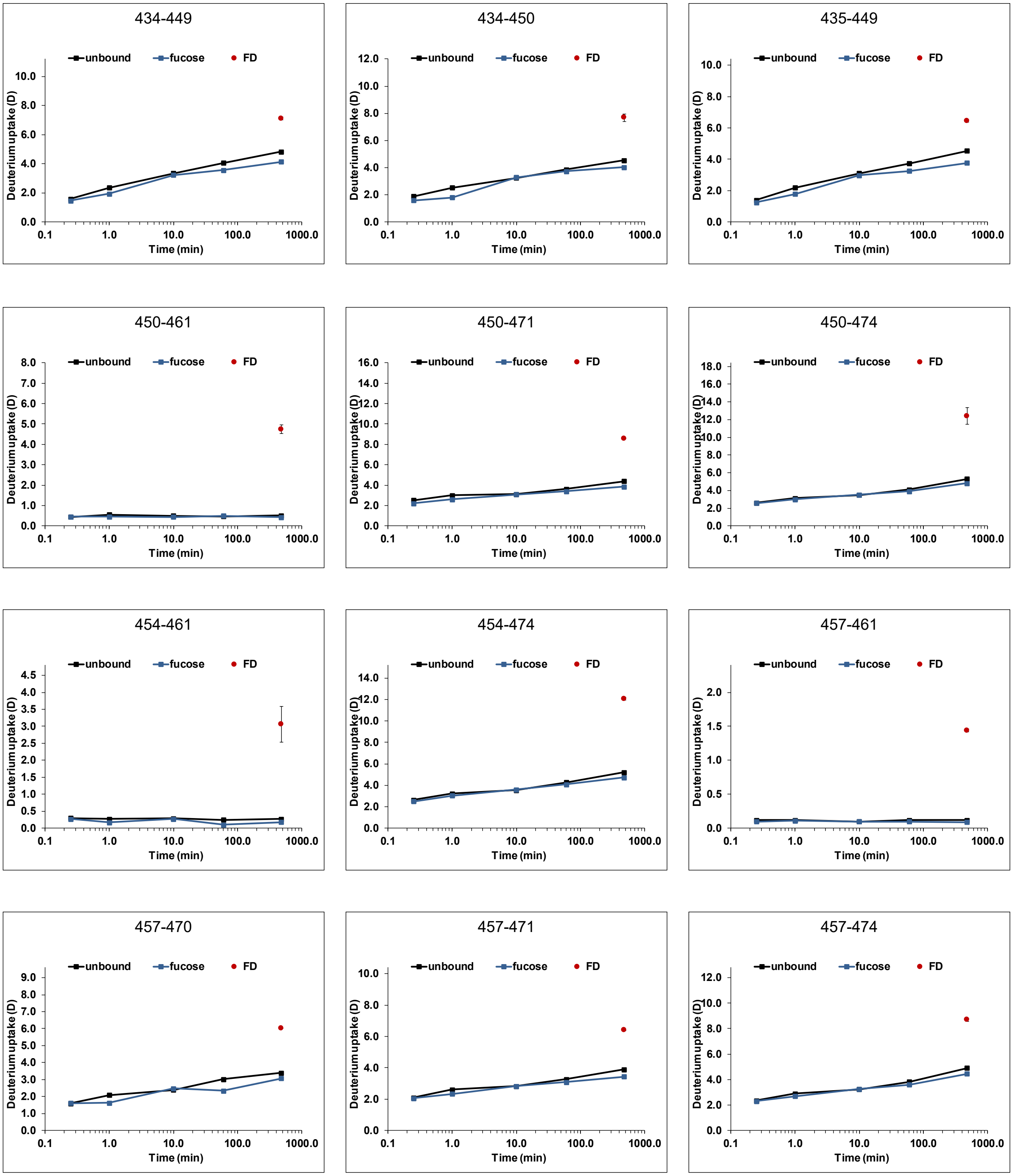


Figure S 29: continued

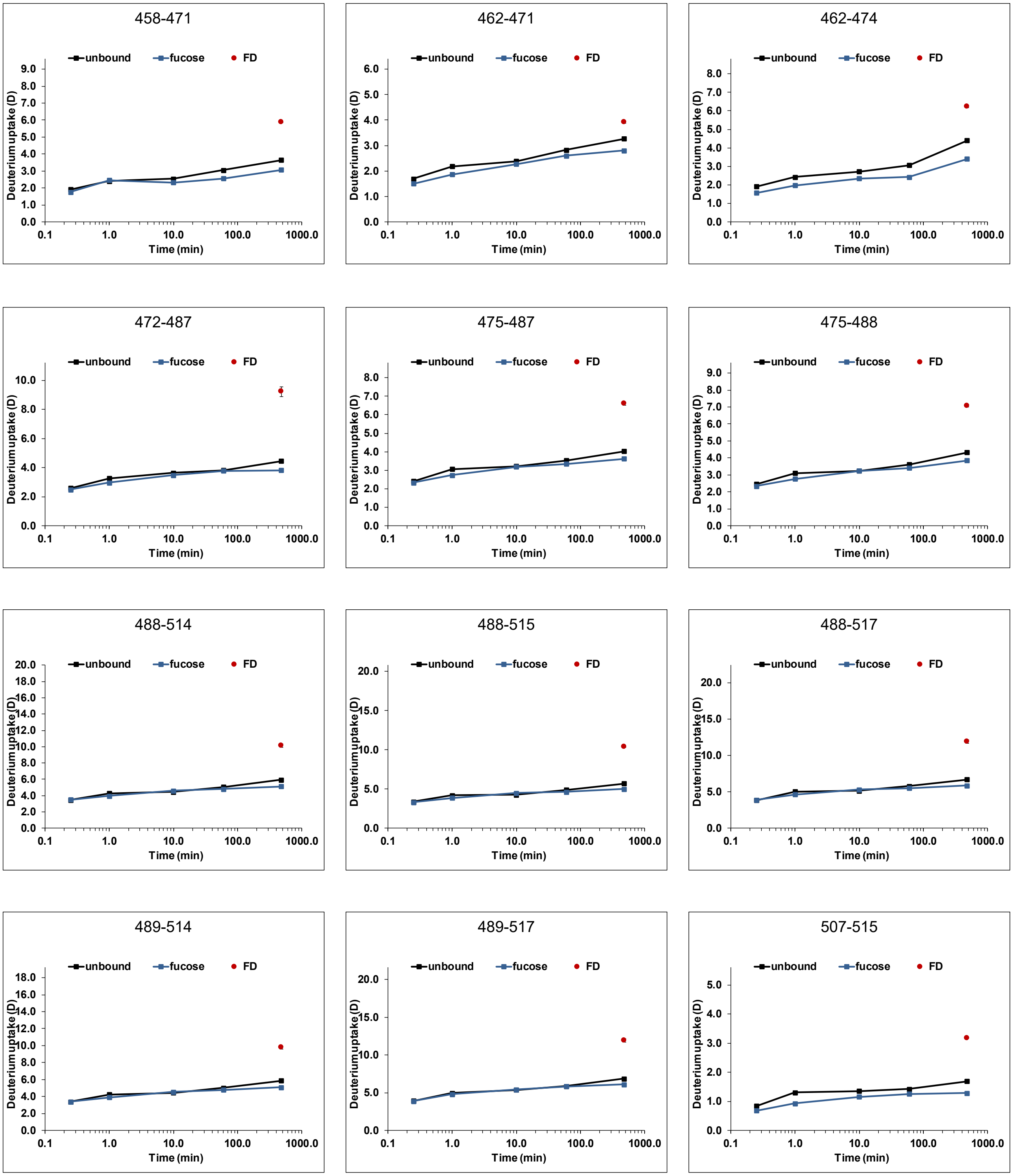


Figure S 29: continued

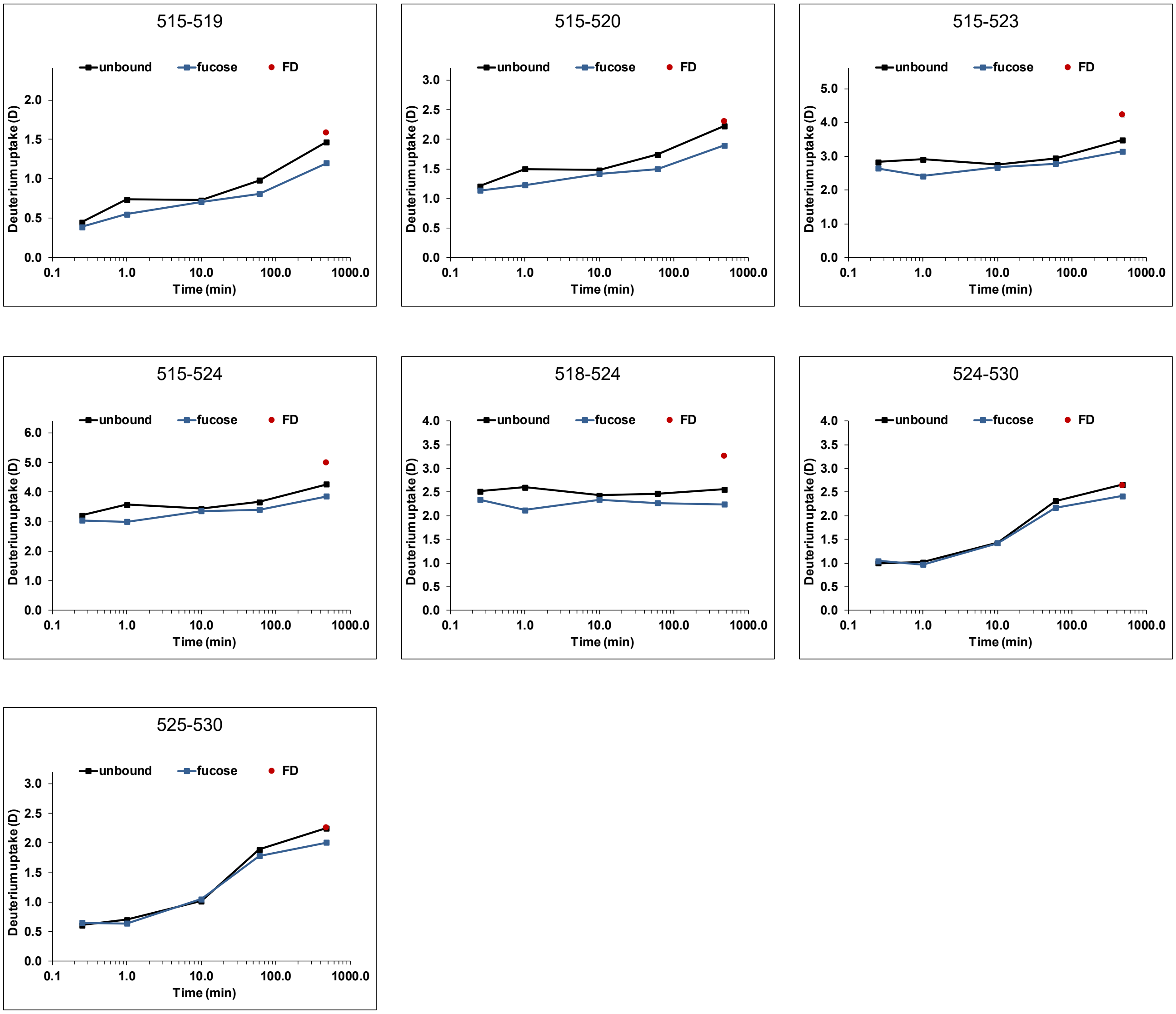


Figure S 29: continued

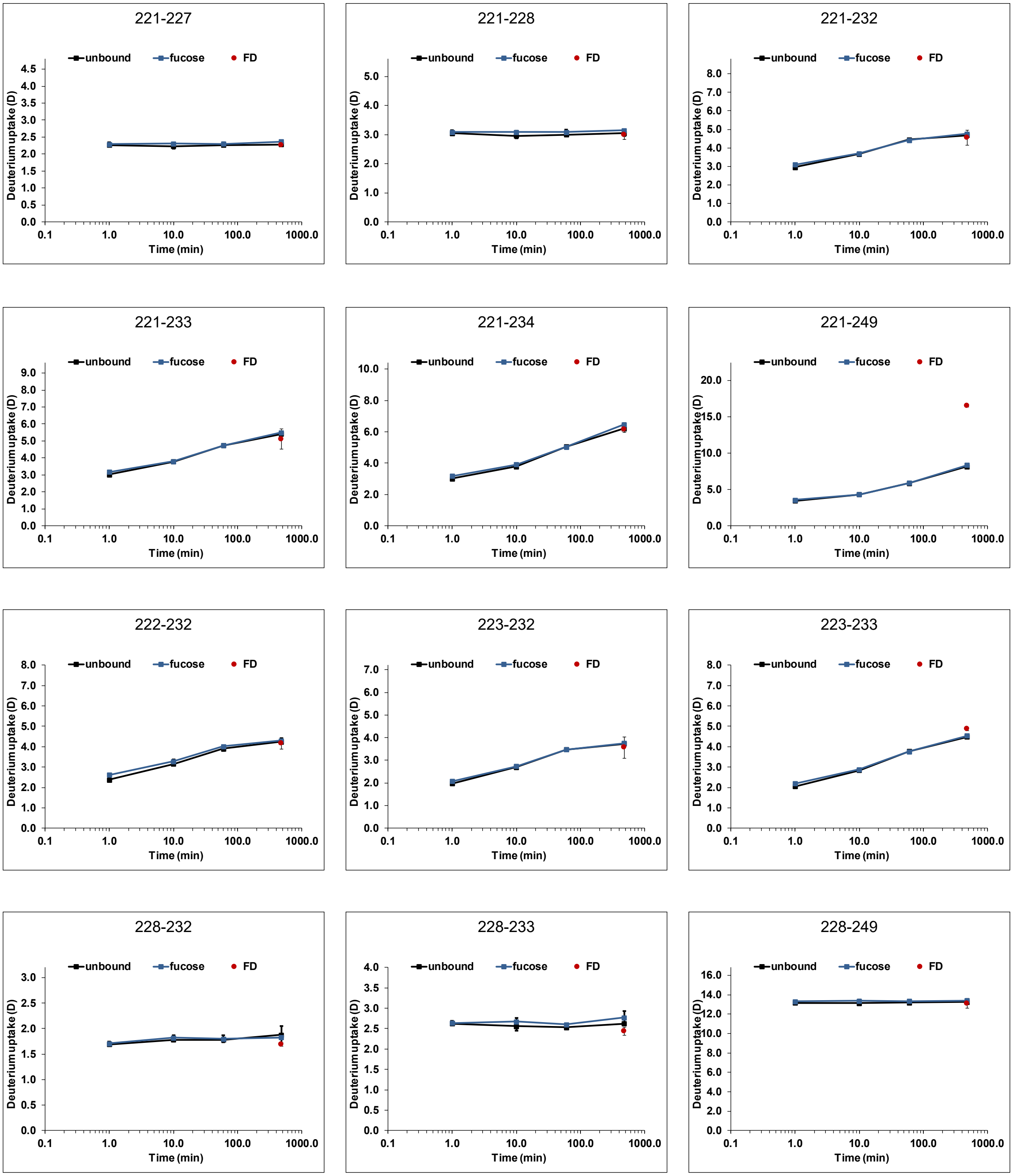


Figure S 30: Deuterium uptake plots for partially deamidated MI001 P dimers (black) with 100 mM fucose (blue) (dataset T7). All time points (1 min, 10 min, 1 h and 8 h) were performed in triplicate, the fully deuterated control (FD, red) represents two measurements. Error bars indicate the standard deviation (SD).

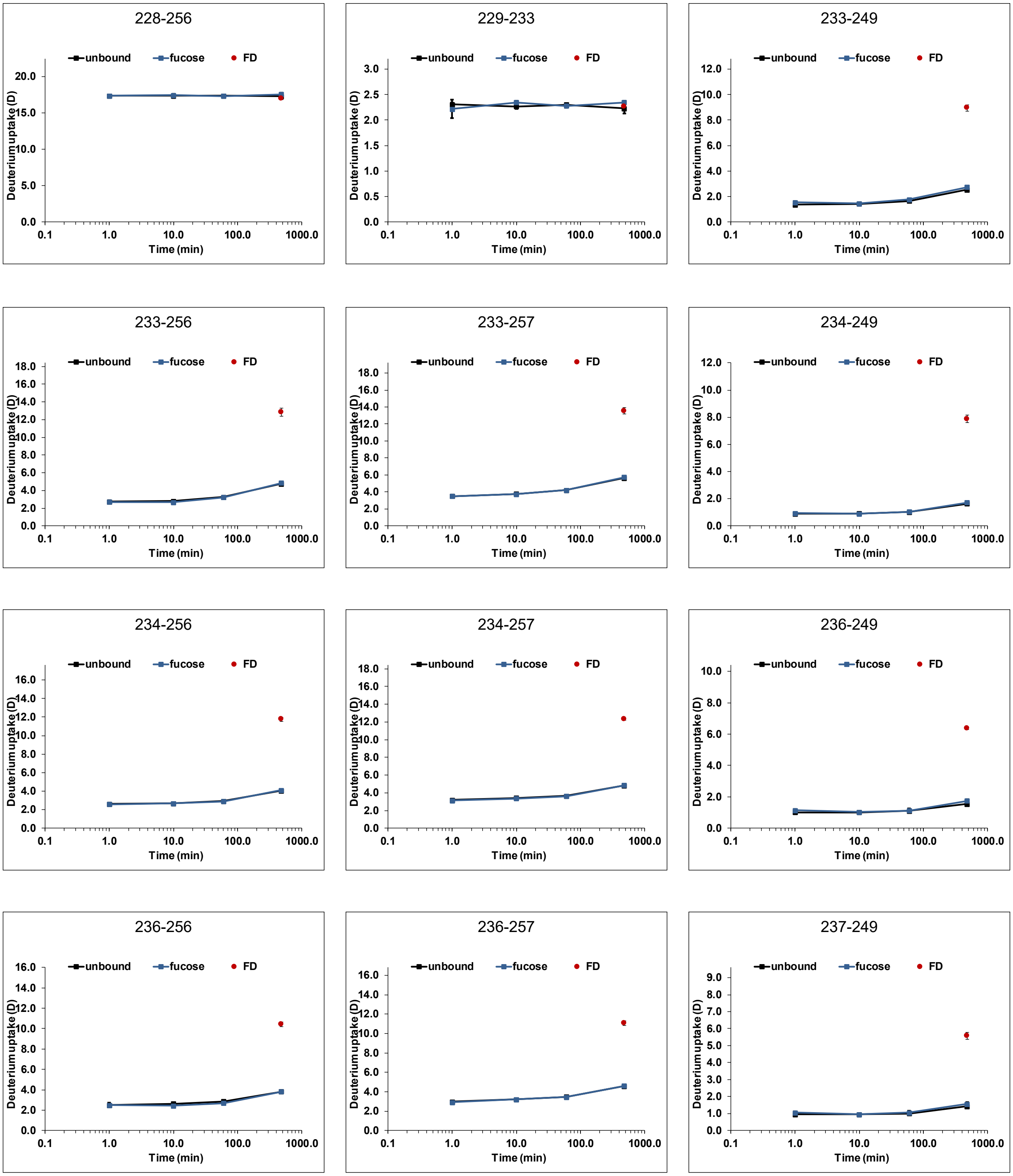


Figure S 30: continued

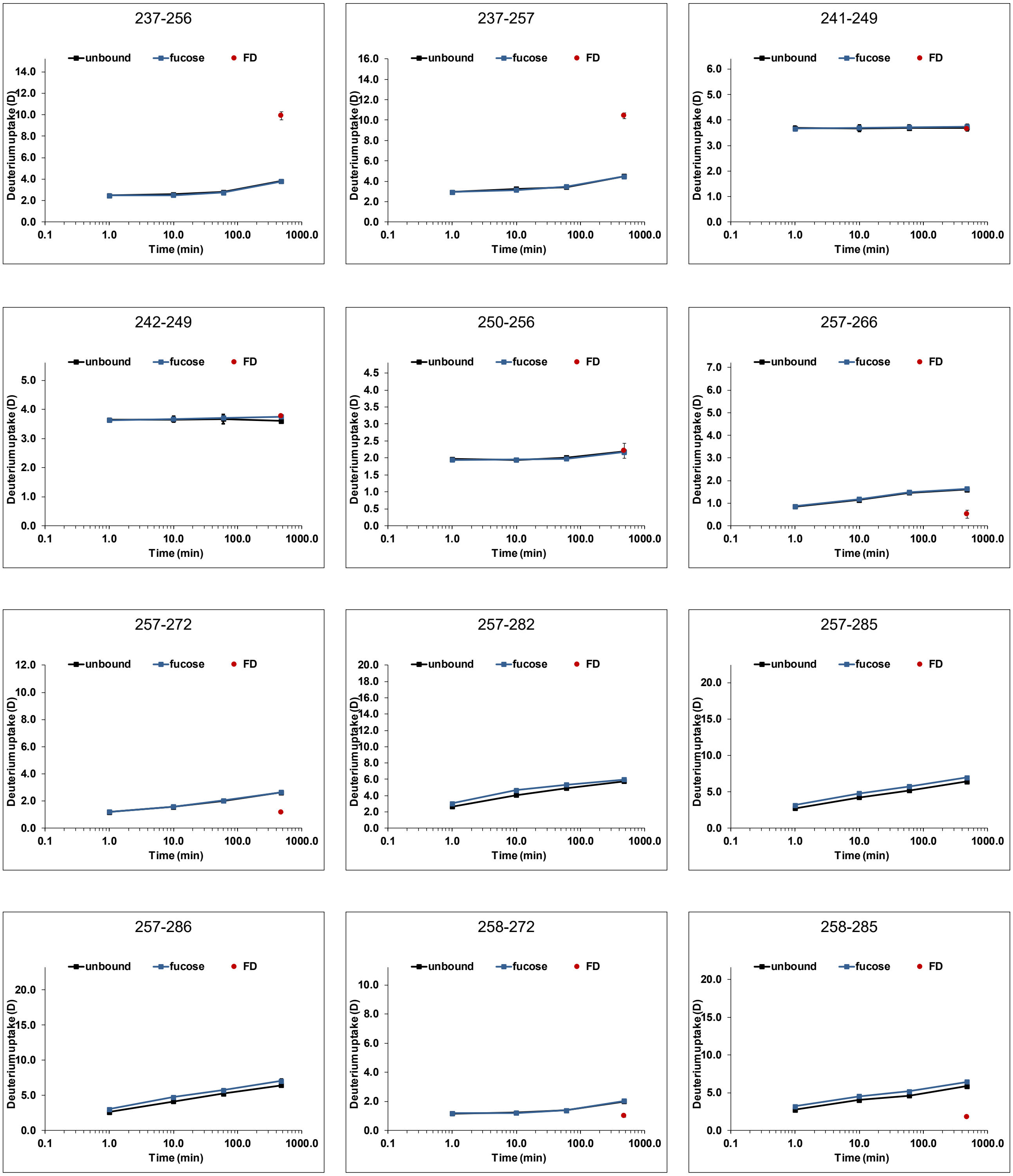


Figure S 30: continued

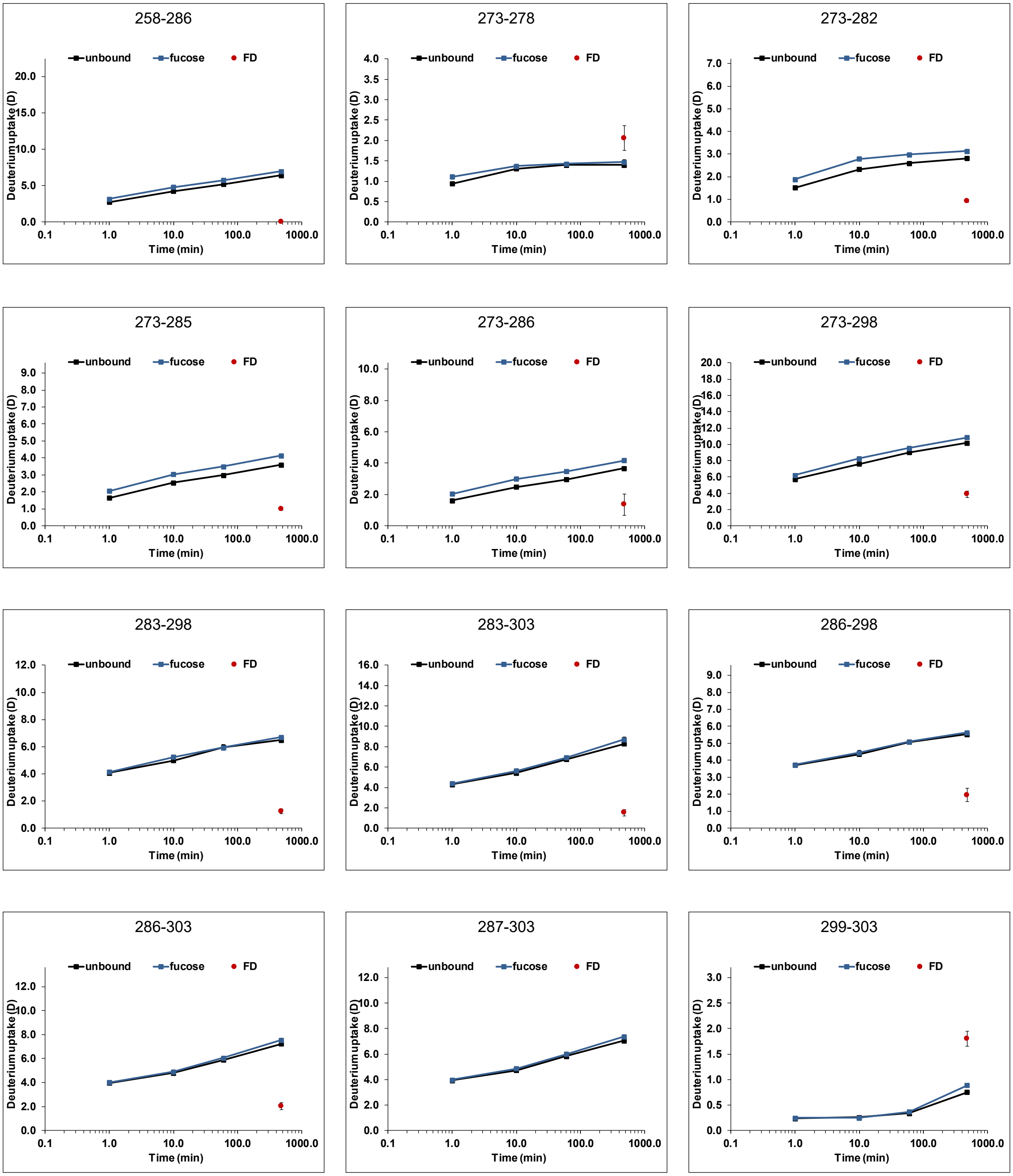

Figure S 30: continued

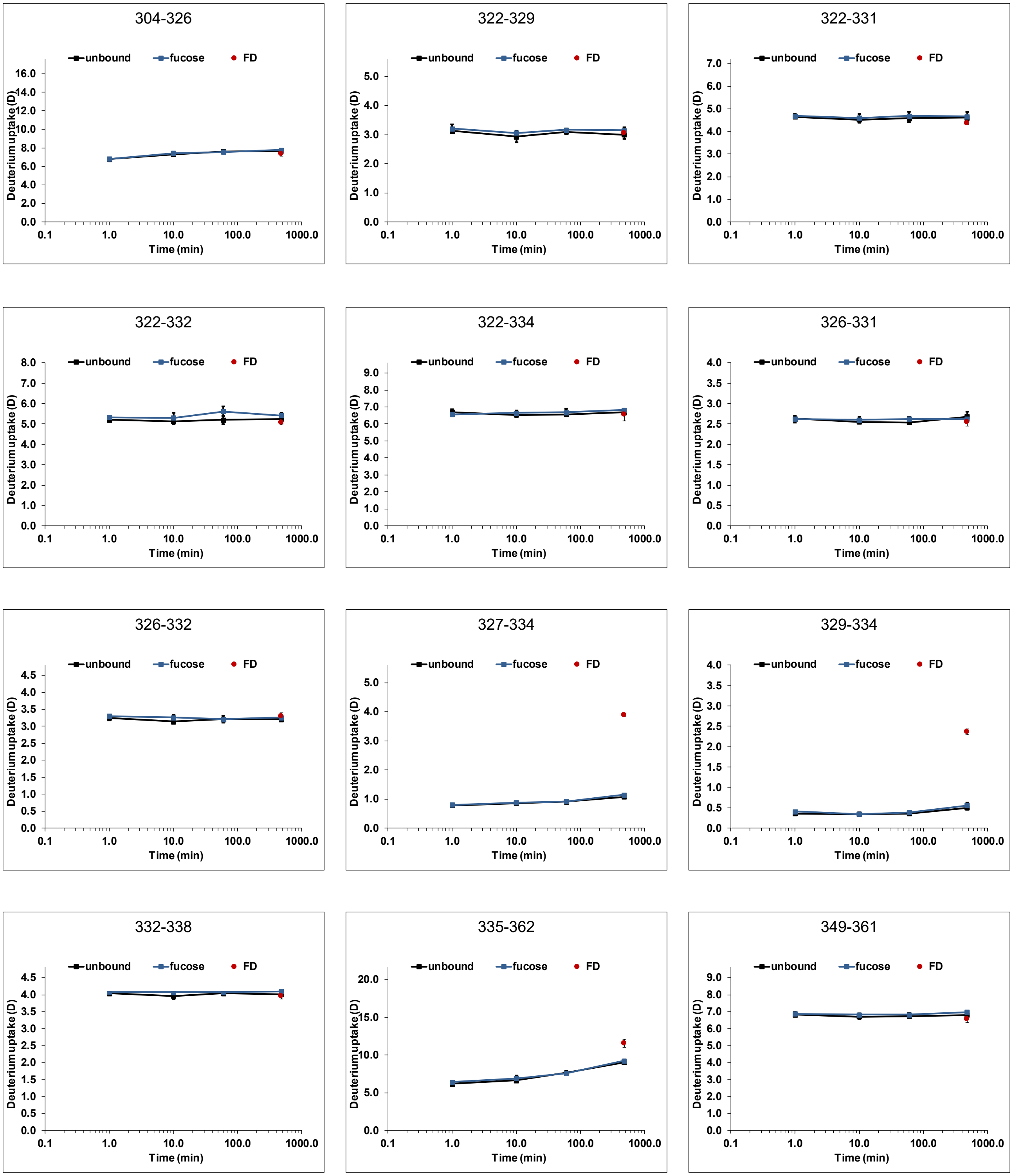


Figure S 30: continued

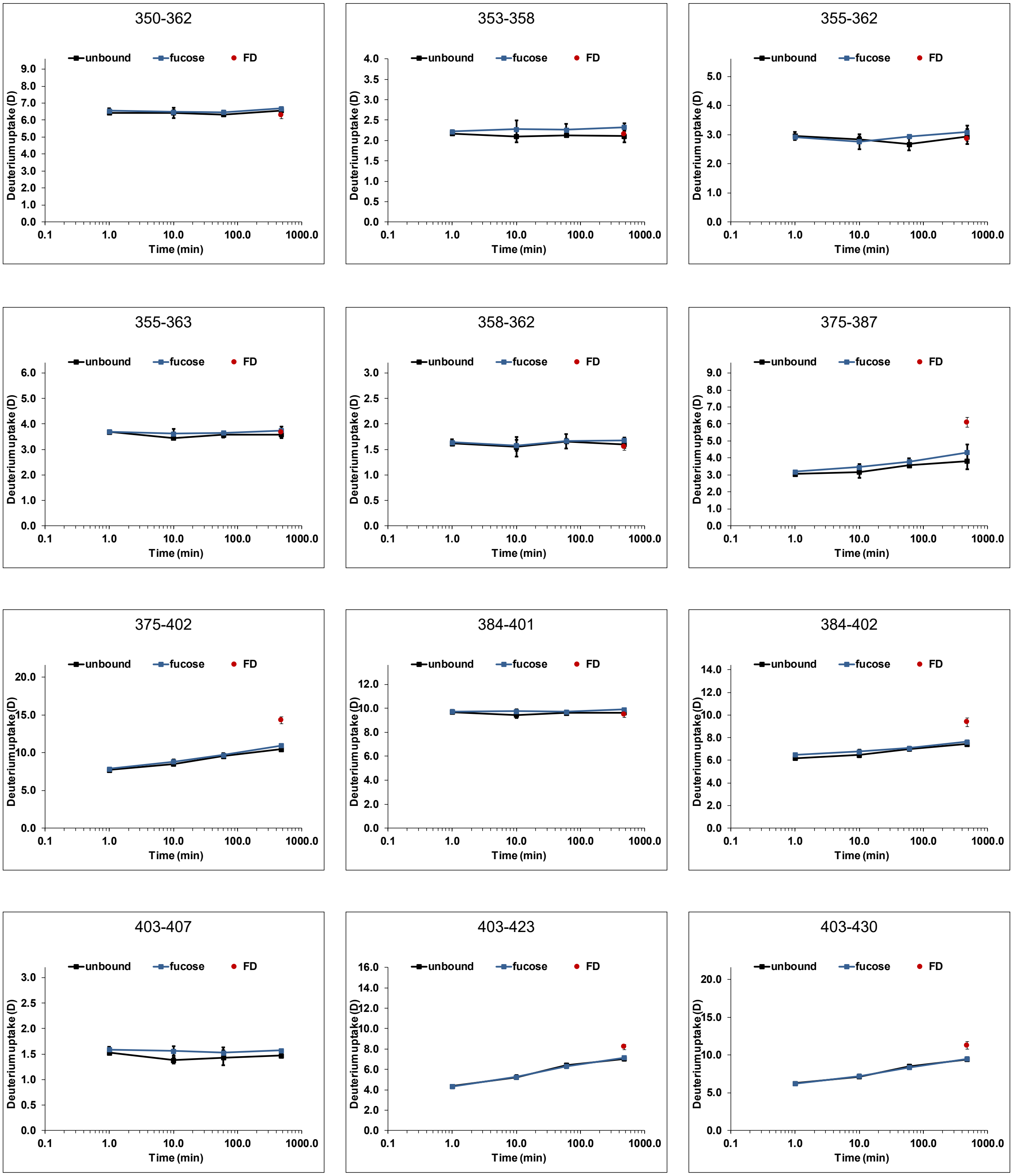


Figure S 30: continued

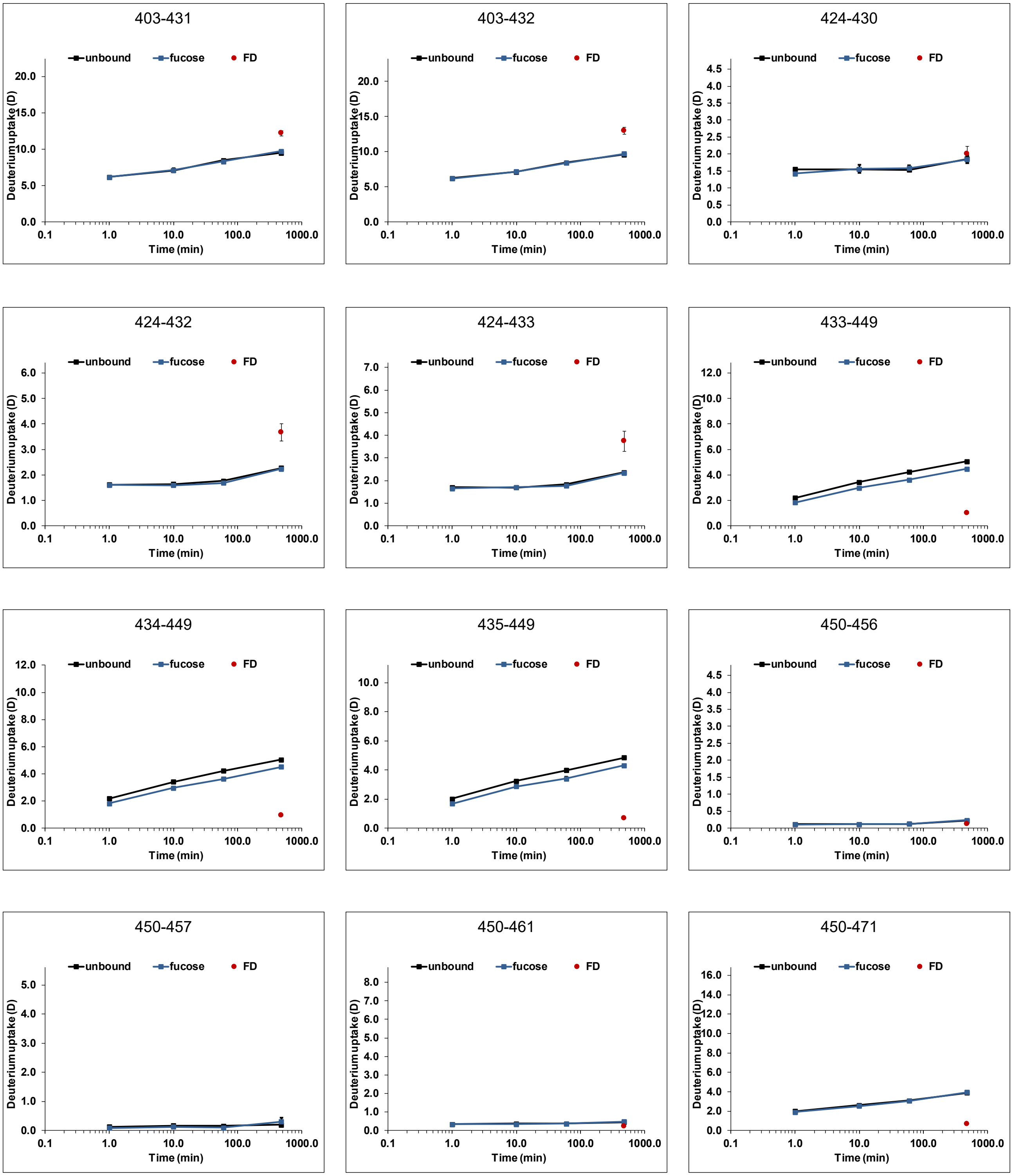


Figure S 30: continued

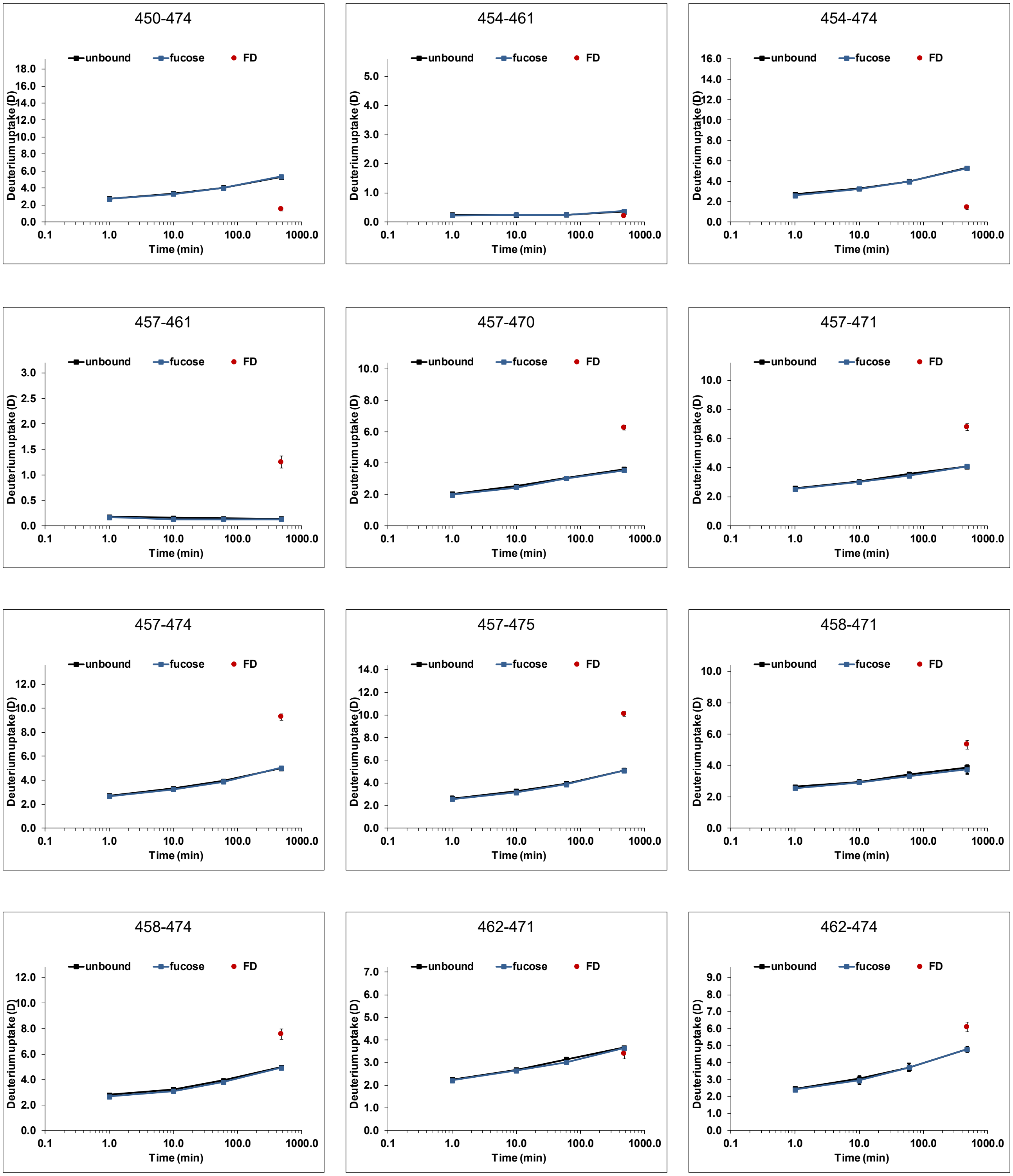


Figure S 30: continued

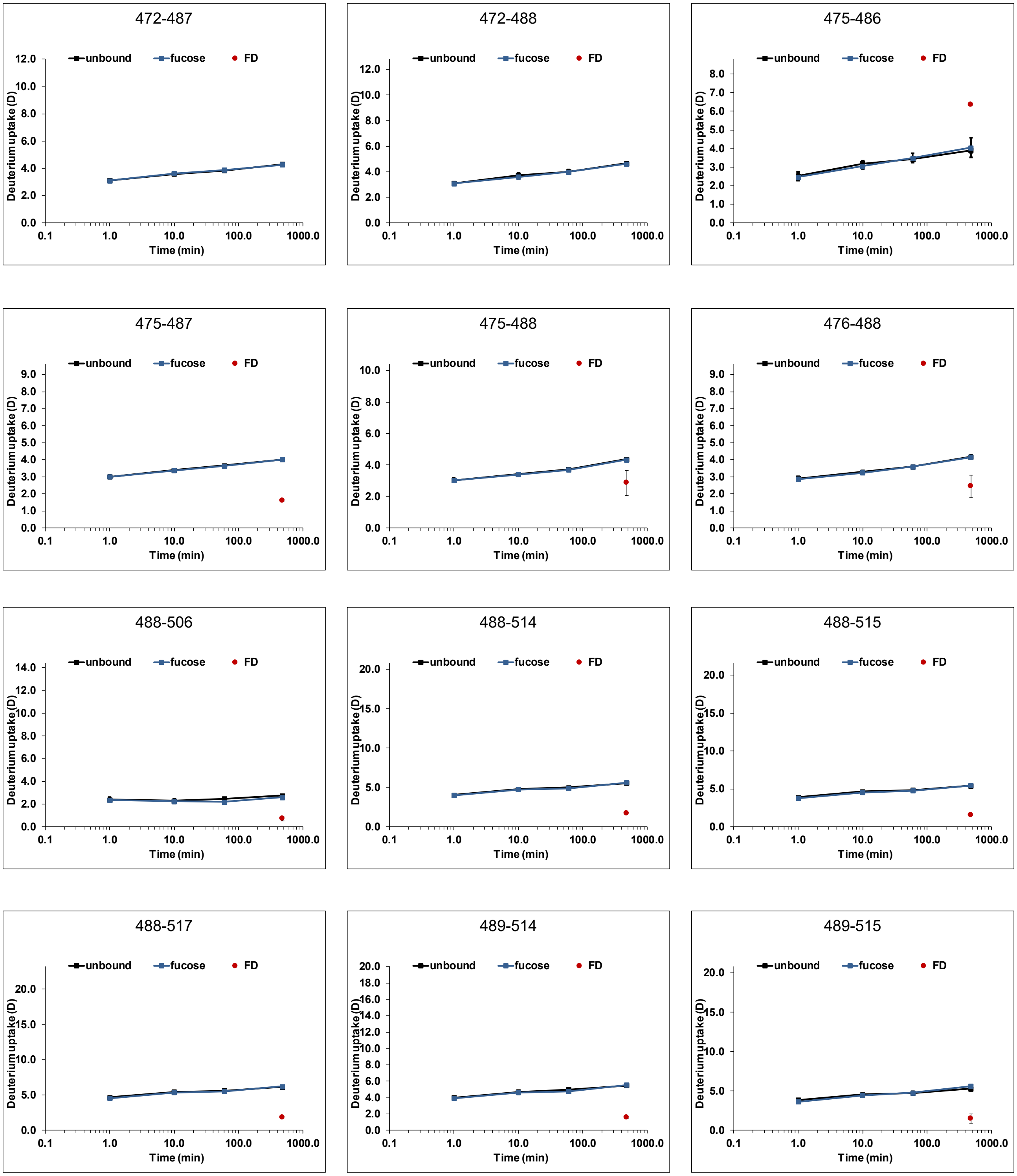


Figure S 30: continued

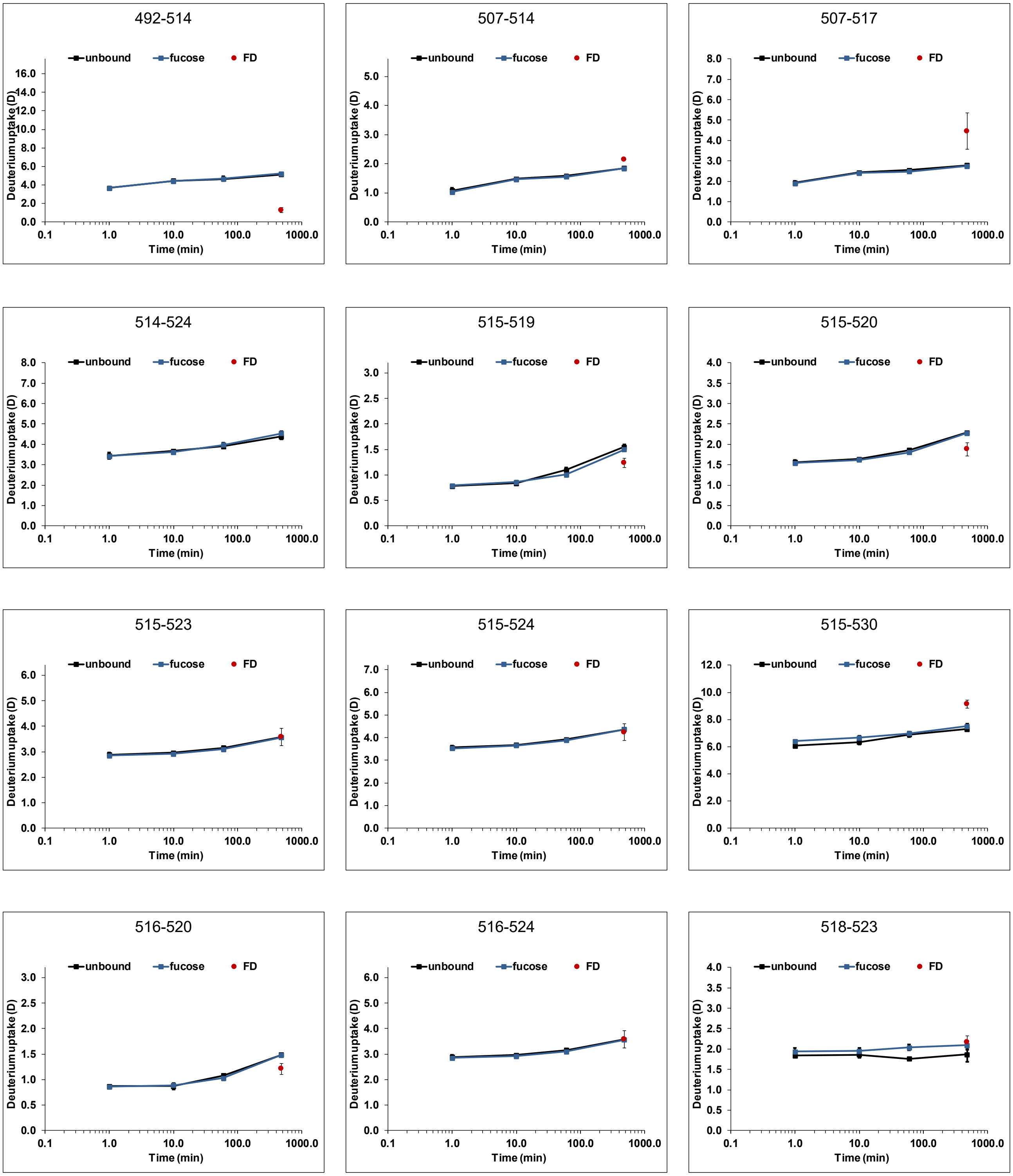

Figure S 30: continued

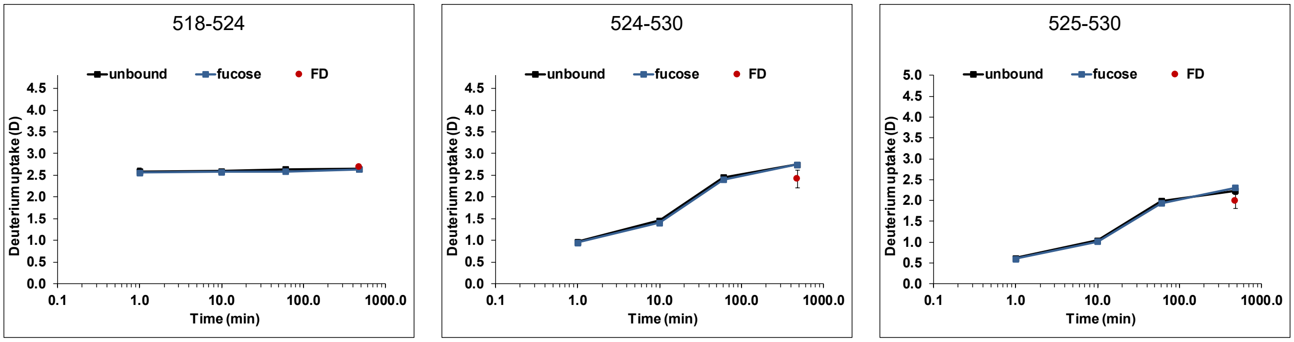


Figure S 30: continued

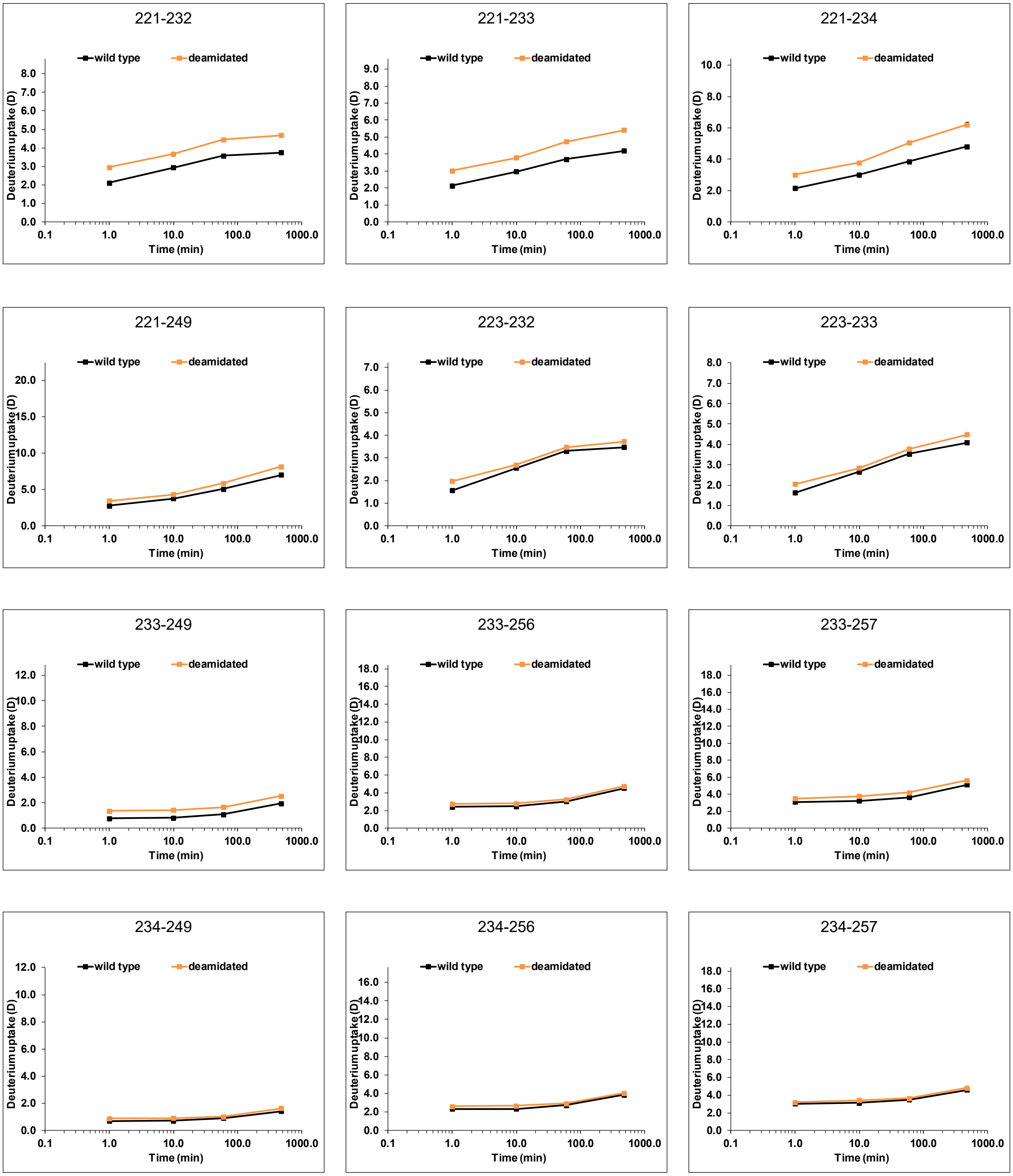


Figure S 31: Deuterium uptake plot comparison for wildtype (black) and partially deamidated MI001 P dimers (orange) without any ligand (dataset T8). All time points (1 min, 10 min, 1 h, 8 h) were performed in triplicate. The ratio of the FD controls from wildtype and deamidated measurements was used for normalization to account for different back exchange levels between the two datasets. Error bars indicate the standard deviation (SD).

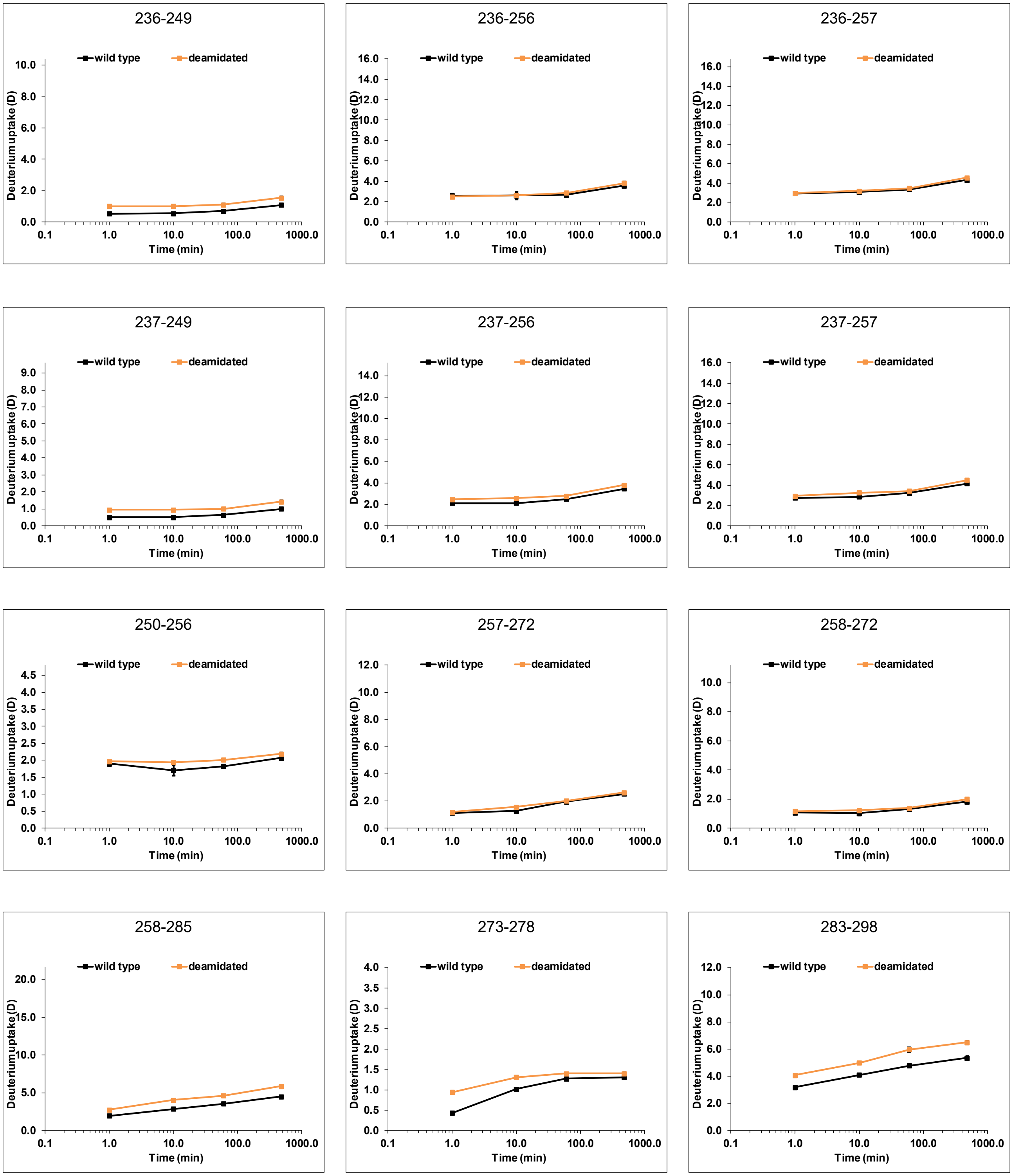

Figure S 31: continued

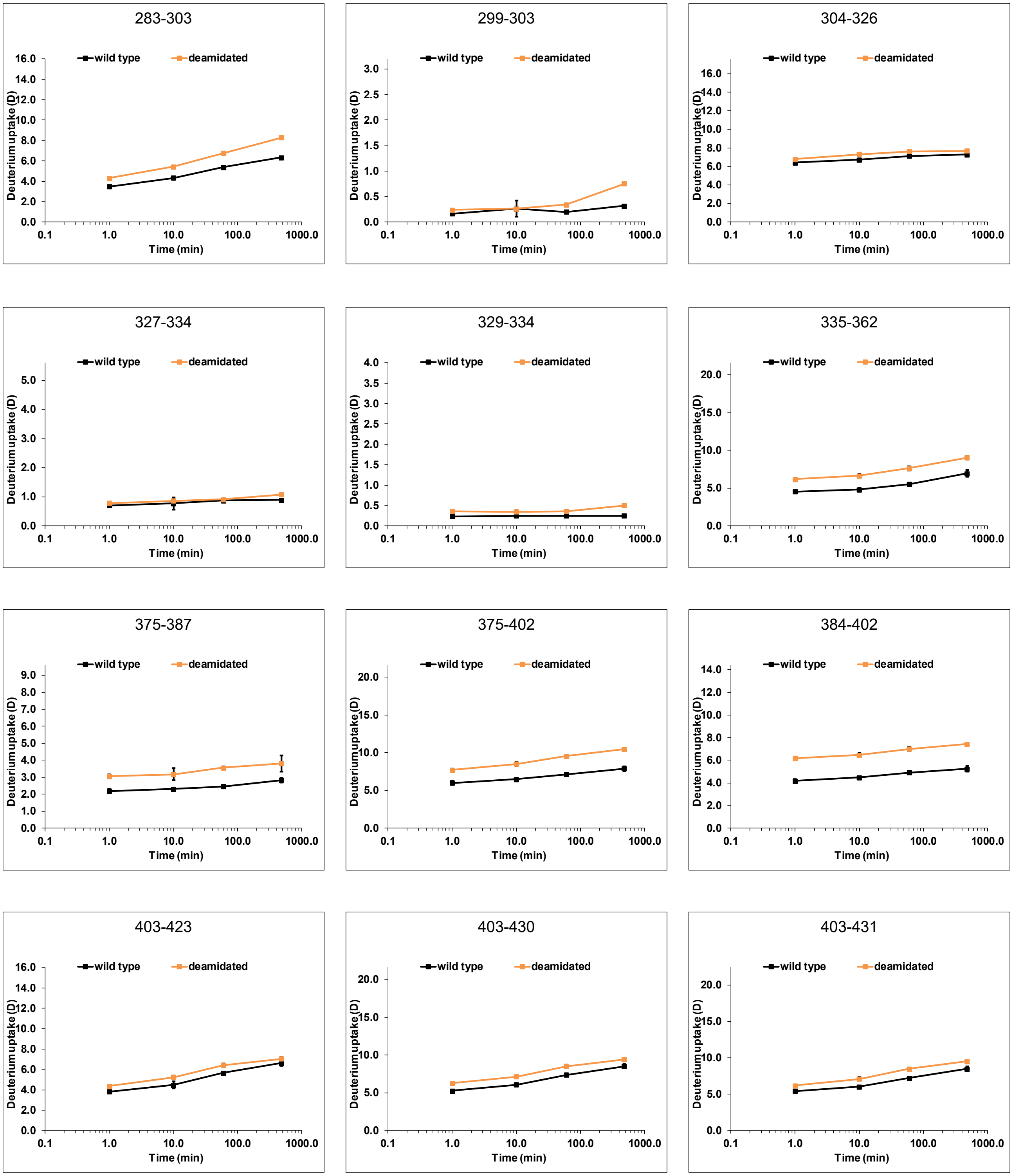


Figure S 31: continued

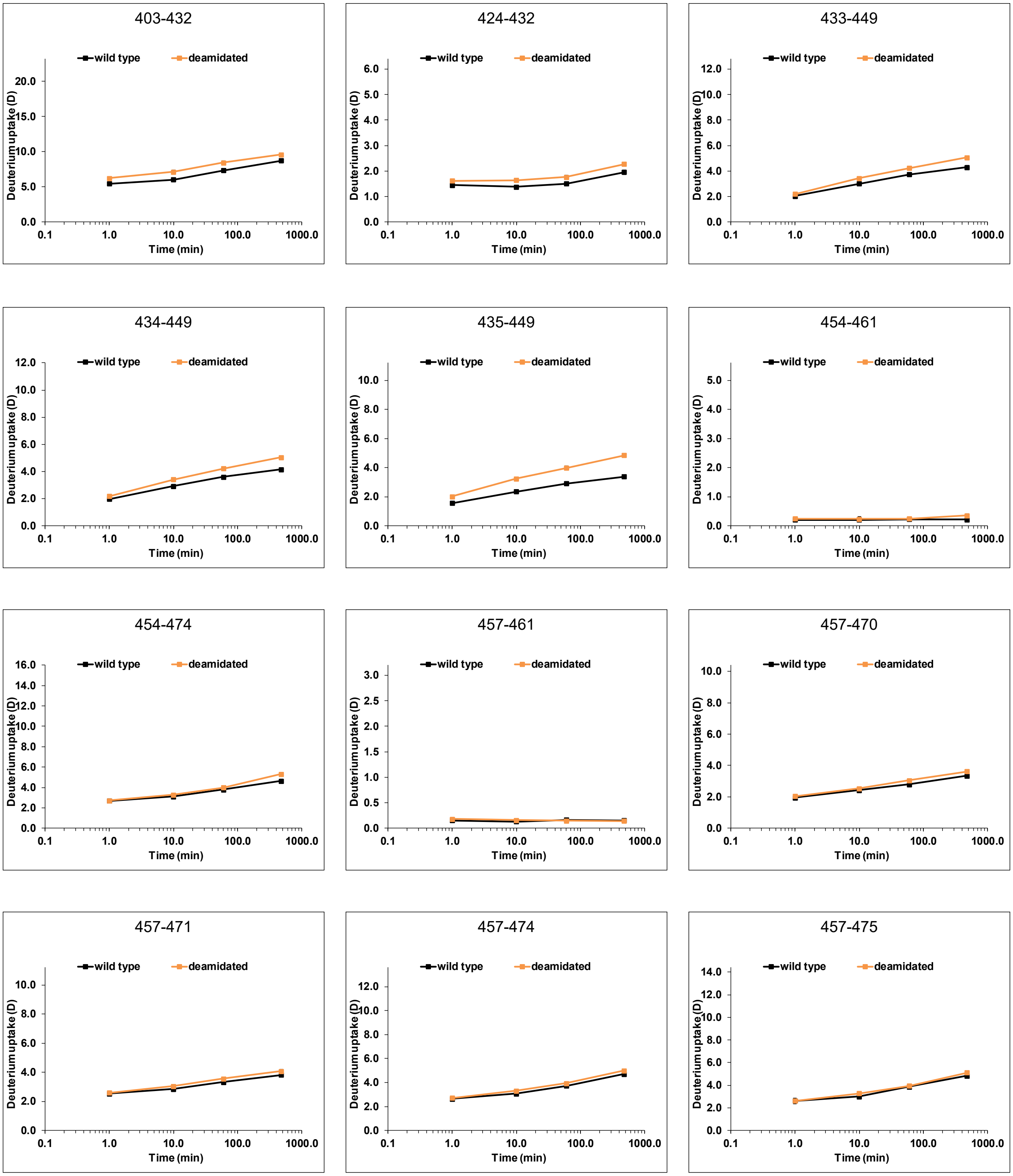


Figure S 31: continued

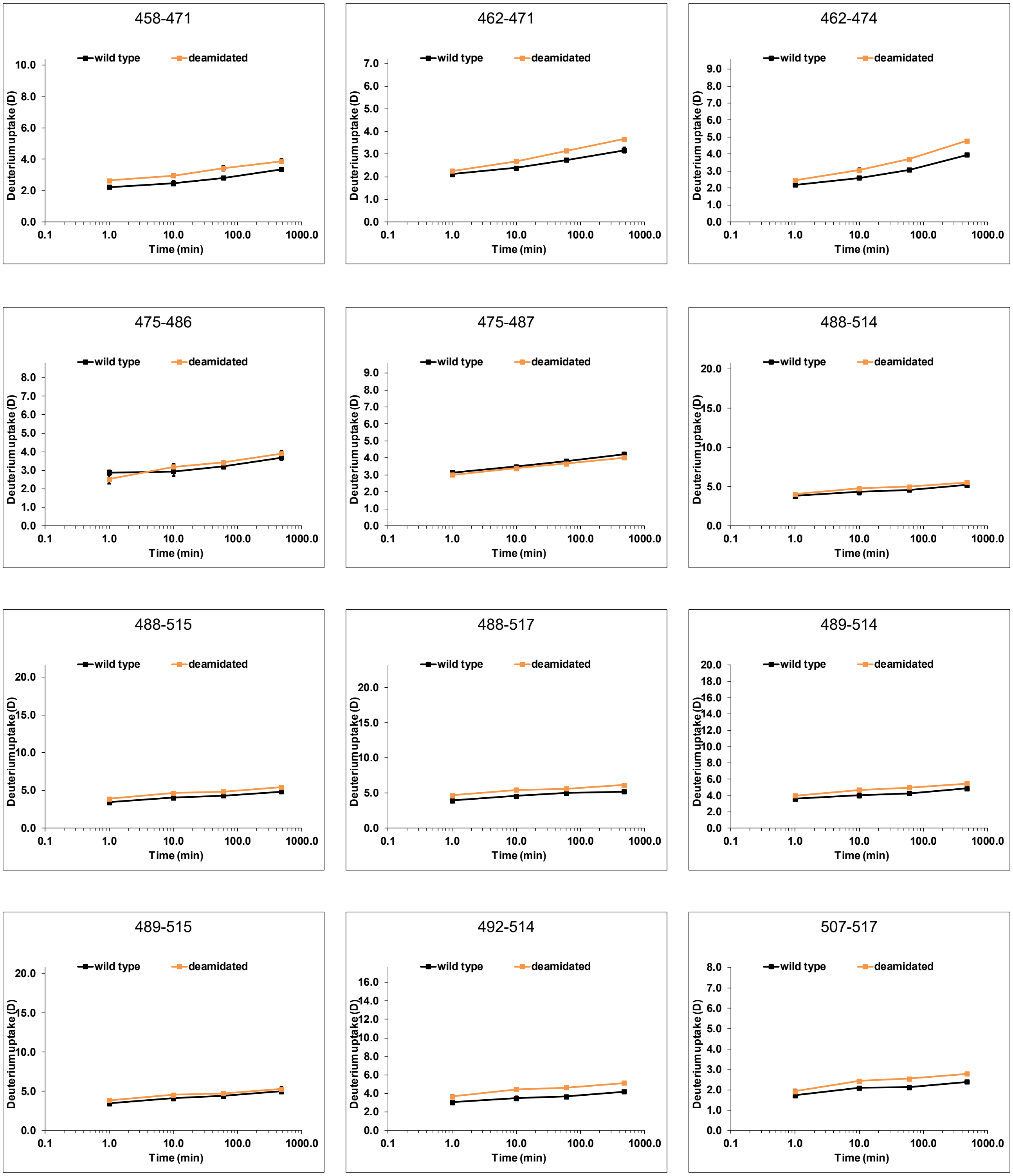


Figure S 31: continued

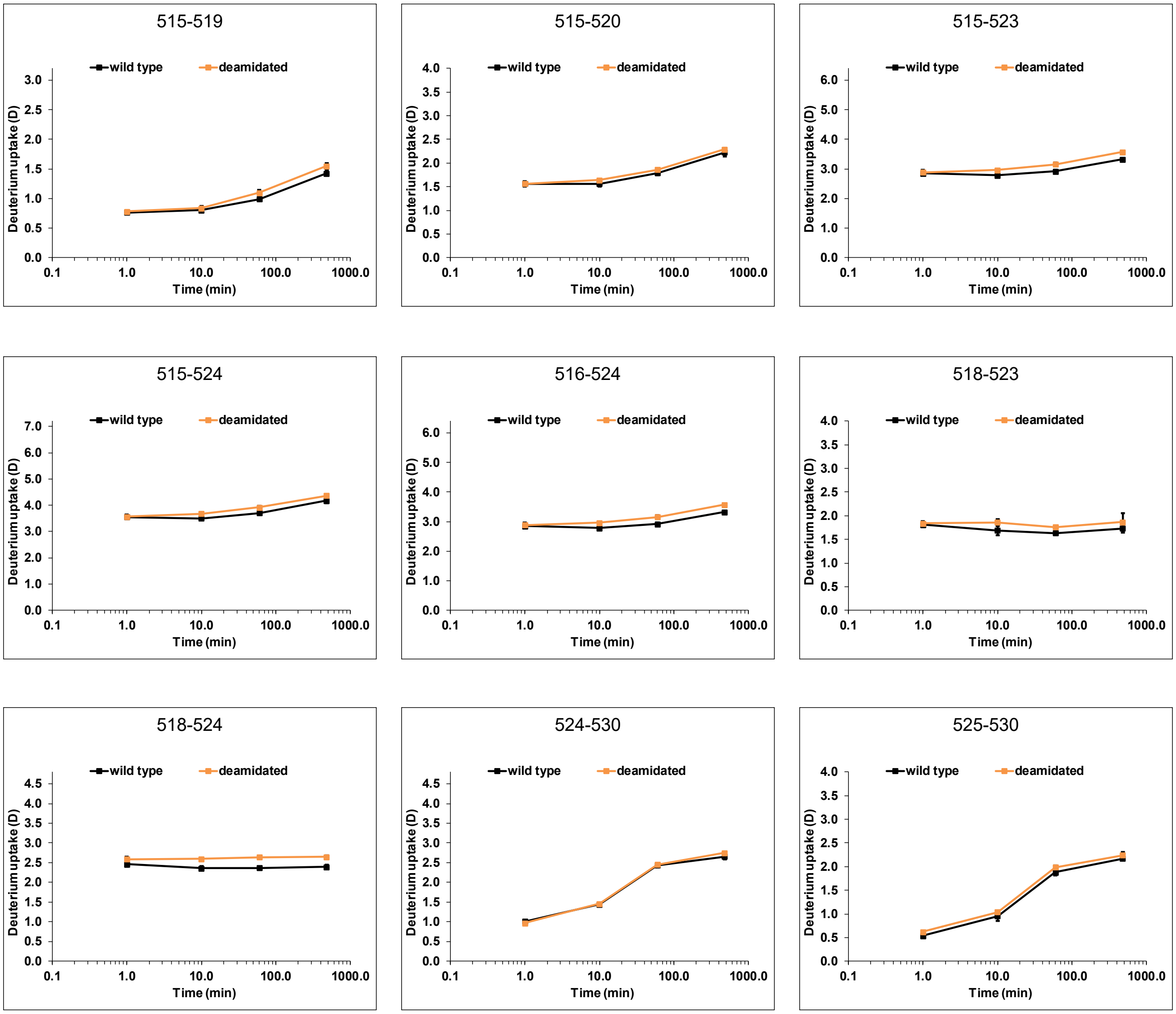


Figure S 31: continued

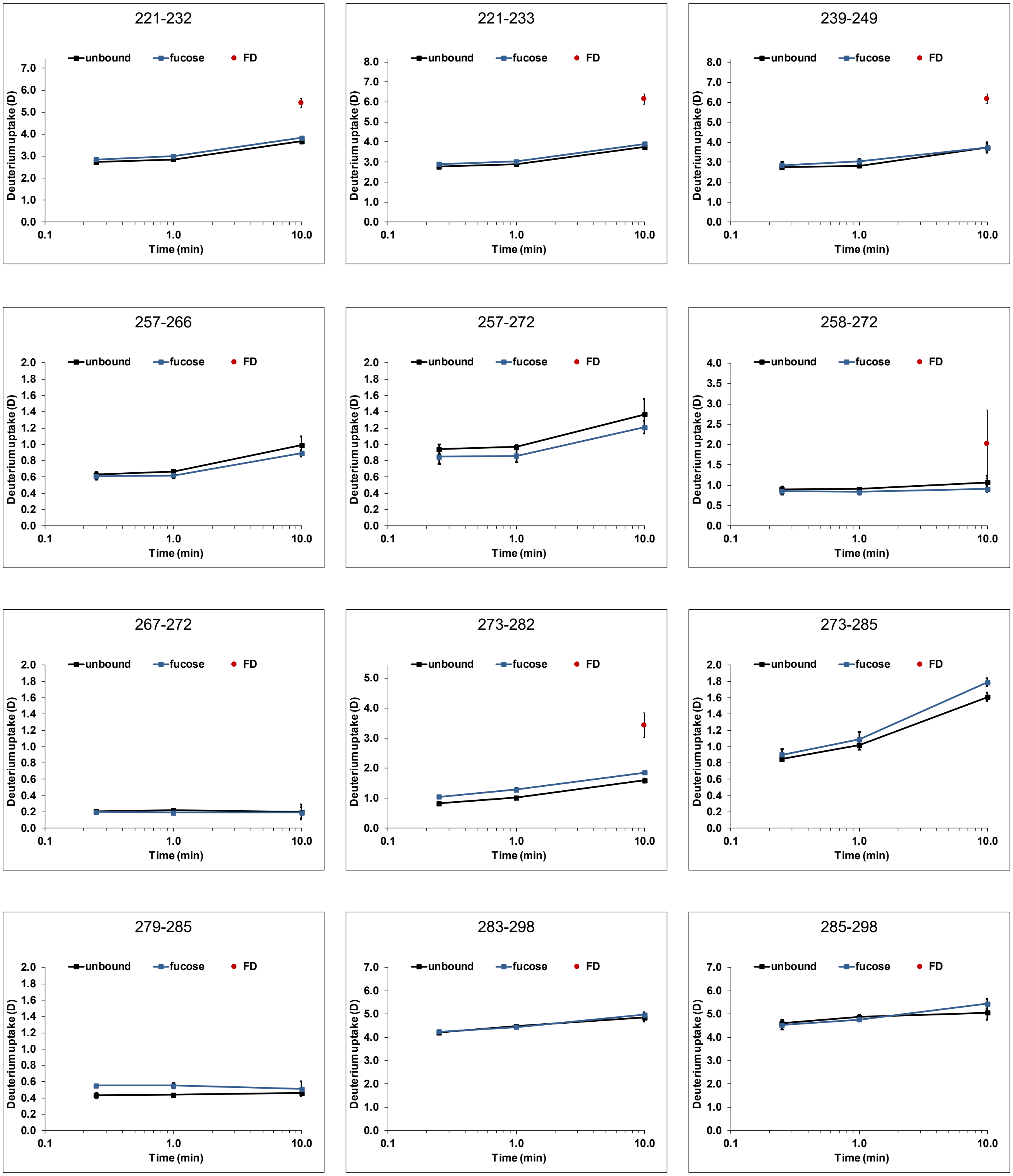


Figure S 32: Deuterium uptake plots for partially deamidated Saga P dimers (black) with 100 mM fucose (blue) (dataset T10). All time points (15 s, 1 min and 10 min) were performed in triplicate, the fully deuterated control (FD, red) represents two measurements. Error bars indicate the standard deviation (SD).

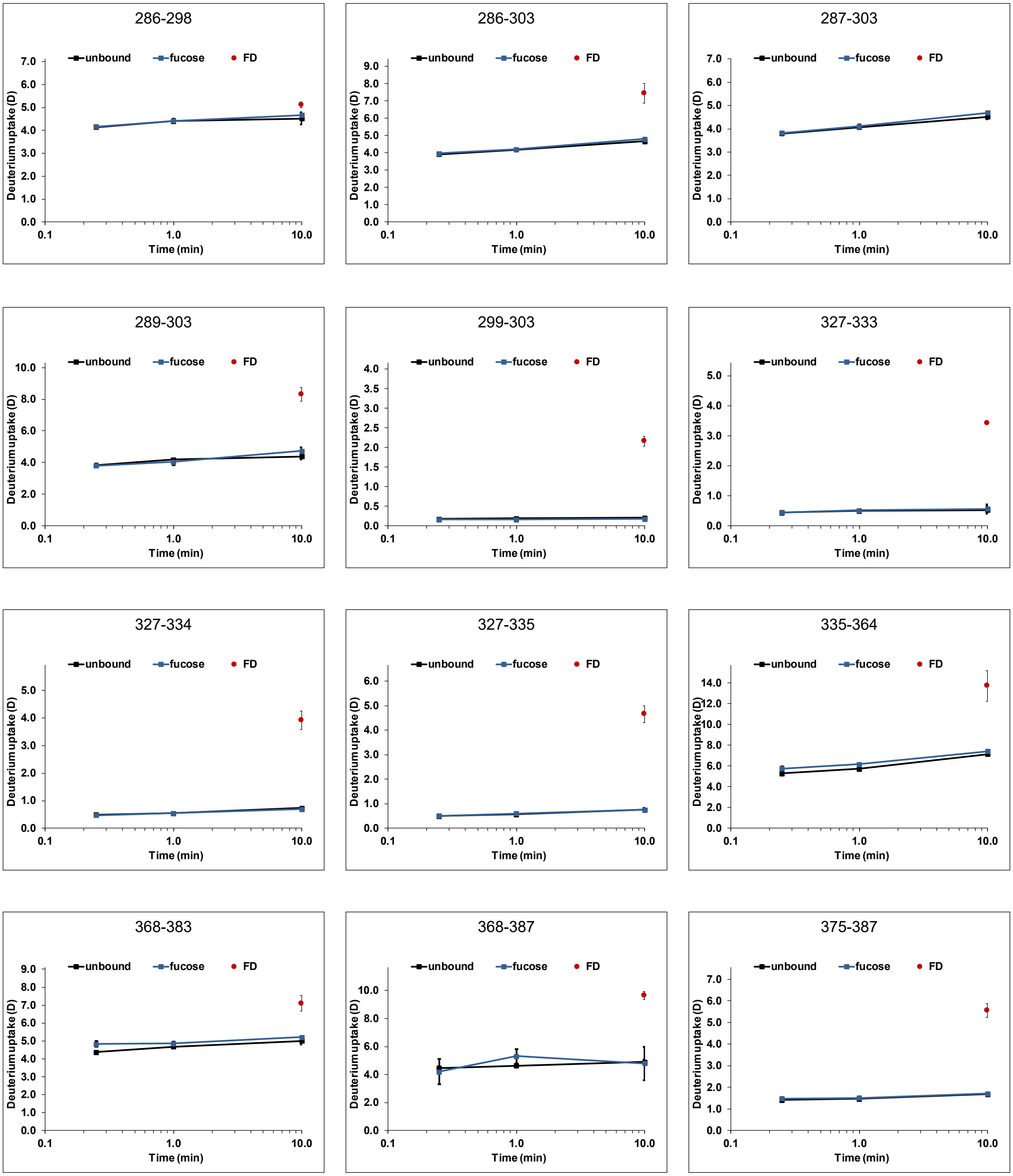


Figure S 32: continued

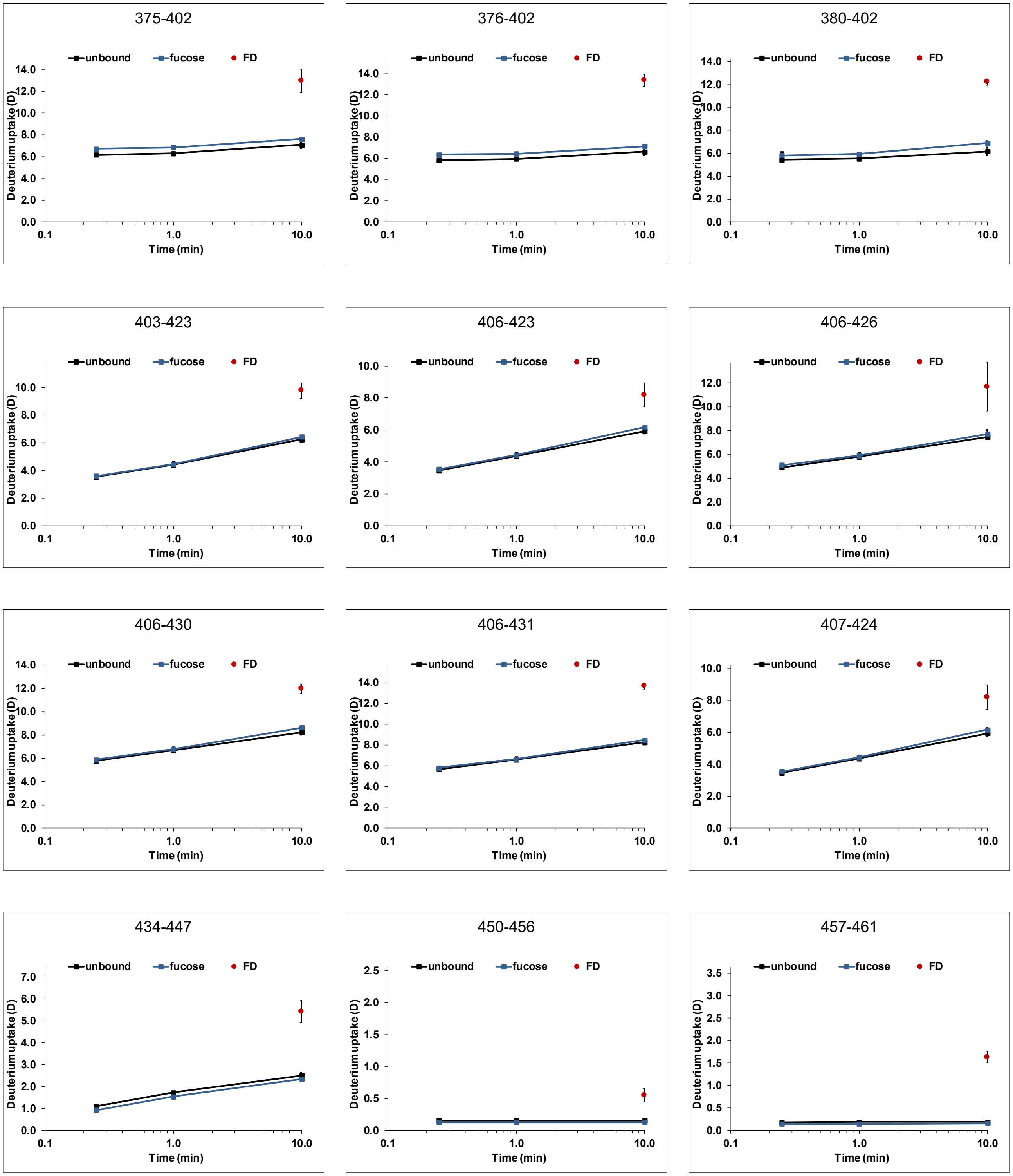


Figure S 32: continued

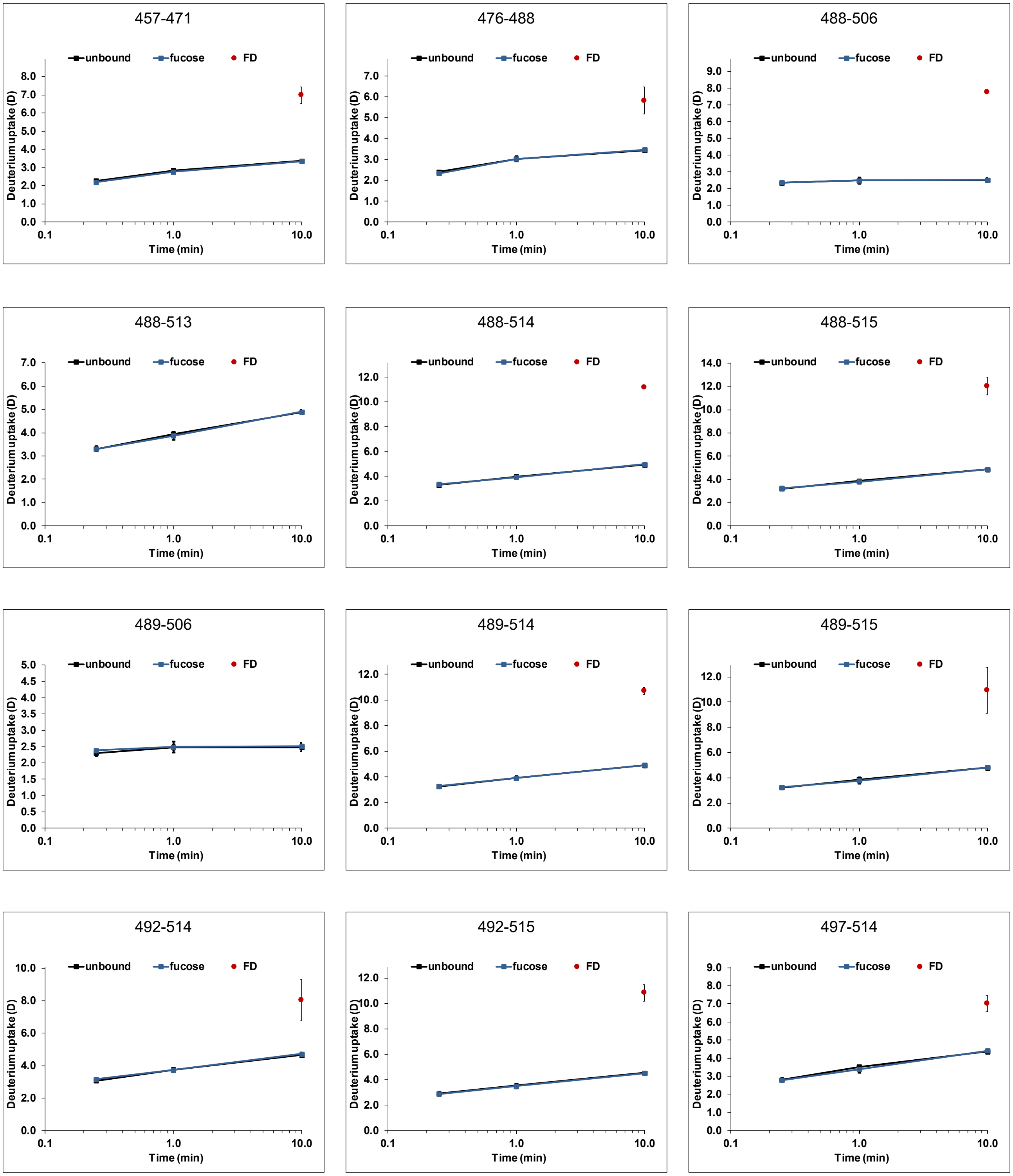


Figure S 32: continued

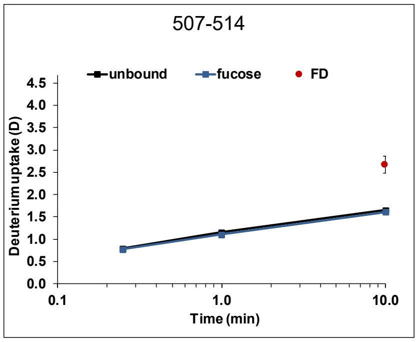


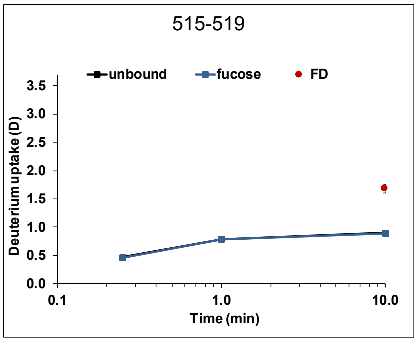


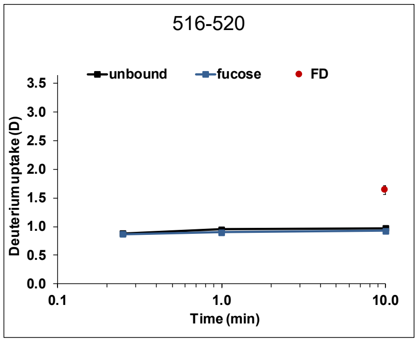


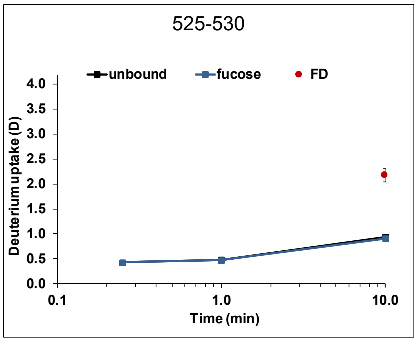


Figure S32: continued
